# A Denisovan-derived *Alu* insertion in *OCA2* contributes to pigmentation diversity in present-day Melanesians

**DOI:** 10.64898/2026.03.18.712481

**Authors:** Kwondo Kim, Aaron Pfennig, Sabriya A. Syed, Nicholas Moskwa, Nélio A.J. Oliveira, Quynh-Mai Pham, Pille Hallast, Feyza Yilmaz, Justin A. McDonough, Heather L. Norton, Joshua M. Akey, Charles Lee

## Abstract

Modern humans inherited DNA from Neanderthals and Denisovans, but the contribution of introgressed structural variants (SVs) to present-day human phenotypes and adaptation remains poorly understood. Here, we used a graph-genome approach to genotype 96,277 SVs in 3,332 present-day humans and three high-coverage archaic hominin genomes, identifying 153 candidate introgressed SVs. These SVs are enriched for signatures of local adaptation compared to non-introgressed SVs (*p-value* = 3.04 × 10^-7^). Among these, we focused on a Denisovan-derived *Alu* insertion located in intron 18 of *OCA2*, a gene central to pigmentation. This introgressed *Alu* insertion is most frequently observed (> 60%) in Indigenous people from Bougainville Island of Melanesia, and is significantly associated with increased skin pigmentation in this region. To assess its functional impact, the *Alu* insertion was introduced into human induced pluripotent stem cells (iPSCs), which were subsequently differentiated into melanocytes. Melanocytes harboring the *Alu* insertion demonstrated elevated *OCA2* expression, increased pigmentation, and higher levels of enhancer activity compared to controls. Collectively, these findings highlight introgressed SVs as a significant source of adaptive and phenotypic diversity in modern humans and implicate the Denisovan-derived *Alu* insertion in *OCA2* in pigmentation variation among present-day Melanesian populations.

## INTRODUCTION

Modern humans admixed with archaic hominins, leaving detectable Neanderthal and Denisovan ancestry in the genomes of present-day individuals ^1,2^. Non-African individuals can trace approximately 2% of their genomes back to Neanderthal ancestors, and Oceanic individuals have an additional 3.5% of their genomes inherited from Denisovans ^1,3^. Although many introgressed alleles were likely deleterious and were removed by purifying selection ^4,5^, a subset of introgressed loci persisted and contributed to adaptation ^6^, influencing traits such as immune response ^7,8^, high-altitude adaptation ^9,10^, pigmentation ^11,12^, and susceptibility to infectious diseases ^13,14^. These findings have established archaic introgression as an important source of human evolutionary innovation.

Despite this progress, most studies of archaic introgression have focused almost exclusively on single-nucleotide variants (SNVs). By contrast, the role of introgressed structural variants (SVs) - defined as genomic alterations of 50 base pairs (bp) or more ^15^-remains poorly understood, with only a small number of studies addressing this question directly ^16–21^. This gap is notable because SVs can exert substantial functional effects, especially when they modify gene structure or regulatory elements. The under-representation of SVs in previous research is partly attributable to technical challenges. Ancient DNA is often damaged and highly fragmented ^22,23^, making SV discovery difficult. However, recent developments in long-read sequencing methods, pangenome-scale SV catalogs, and graph-based genotyping now enable more reliable detection and genotyping of SVs in both modern and archaic genomes ^24^. These advances provide an opportunity to evaluate SVs’ contributions to introgression, adaptation, and phenotypic diversity ^19,20^.

Here, we use a graph-based approach to genotype 96,277 SVs in 3,332 present-day humans and three high-coverage archaic hominin genomes, identifying 153 candidate introgressed SVs inherited from Neanderthals or Denisovans. We show that these introgressed SVs are enriched for signatures of local adaptation compared to non-introgressed SVs (*p-value* = 3.04 × 10^-7^), suggesting their potential contribution to human phenotypic diversity. Among these, we focus on a Denisovan-derived *Alu* insertion in intron 18 of *OCA2*, a key pigmentation gene, due to its strong population differentiation and biological plausibility. This *Alu* insertion is found at an unusually high frequency (> 60%) in Indigenous people from Bougainville Island of Melanesia, who have the highest level of skin pigmentation observed across the world ^25^. By combining population genetic analyses, pigmentation measurements across Bougainville-area populations, and genome editing in human induced pluripotent stem cells (iPSCs), we identify this introgressed regulatory insertion as a functional contributor to pigmentation variation and highlight the broader value of studying archaic SVs in human genomes.

## RESULTS

### Identification of introgressed structural variants (SVs)

To identify SVs introgressed from archaic hominins, we first genotyped SVs using a *k*-mer-based approach on a graph genome constructed from the Human Genome Structural Variation Consortium (HGSVC) ^24^. This analysis yielded a callset of 30,889 insertions (≥ 50 bp), 19,962 deletions (≥ 50 bp), 748,812 small indels (< 50 bp), and 12,328,706 SNVs, with a mean genotyping rate of 99.84%. We next searched for SVs with the expected signature of archaic introgression ^4,5^. Specifically, we identified SVs present in at least one archaic genome and one non-African genome, but absent from great ape (n = 34) and African genomes (n = 592) (**Fig. 1A**; see Materials and Methods). This strategy enriches for introgressed SVs by removing ancestral polymorphisms that predate the split between modern humans and archaic hominins. After filtering, we identified 153 candidate introgressed SVs: 97 likely of Neanderthal origin, 37 of Denisovan origin, and 19 shared between both archaic lineages (**Fig. 1B** and **Table S1**).

**Figure 1.**
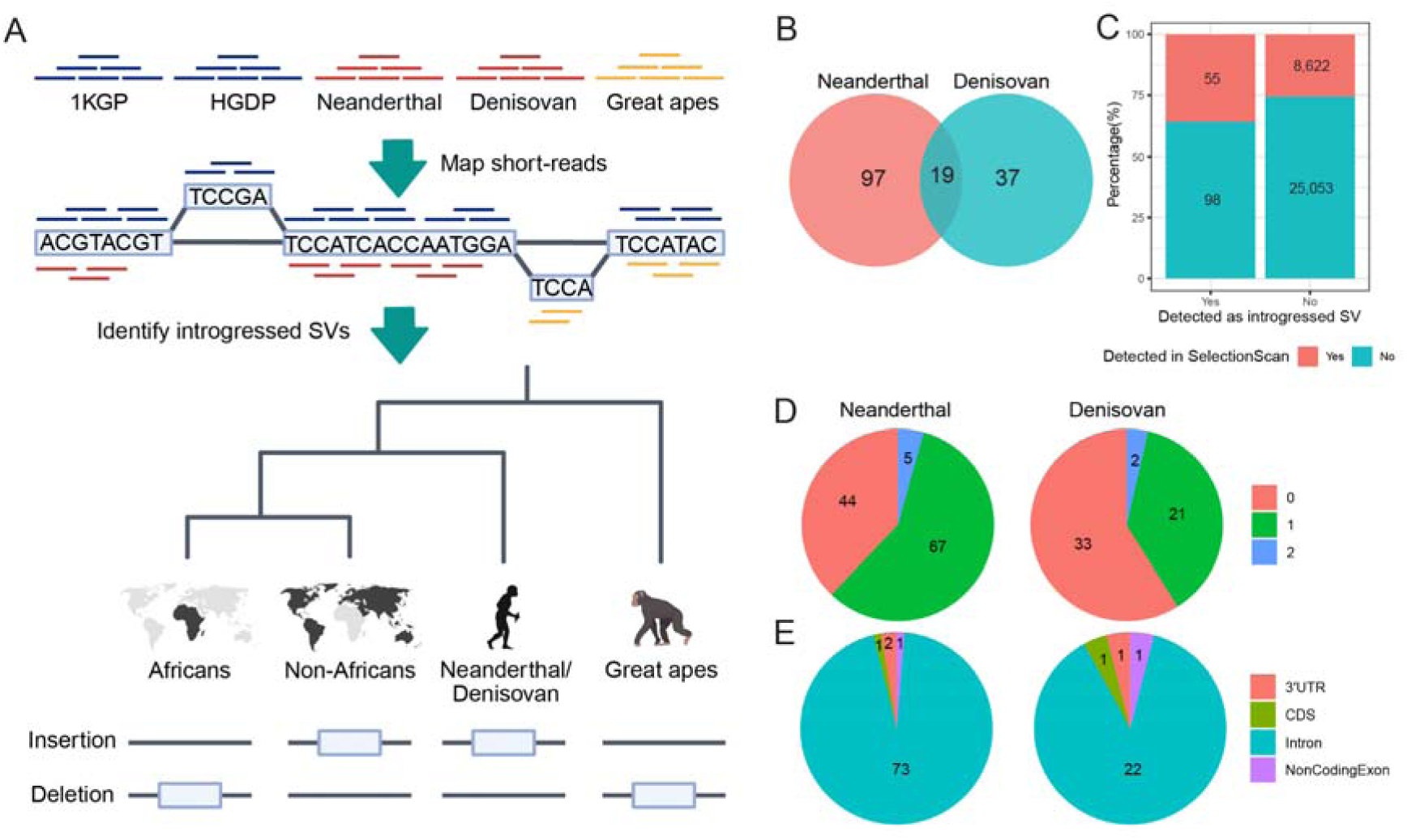
Identification and characterization of introgressed structural variants (SVs). (**A**) A schematic overview of the process to identify introgressed SVs. (**B**) Inferred source of introgressed SVs. (**C**) Enrichment of the selection signal in the introgressed SVs. Introgression and selection signals were significantly associated (Fisher’s exact test: odds ratio = 2.49; *p-value* = 3.04 x 10^-7^). (**D**) Distribution of introgressed SVs annotated with 0, 1, or 2 protein□coding genes. (**E**) Distribution of introgressed SVs overlapping 3′ UTRs, coding sequences (CDS), introns, and non□coding exons. [Figure created with BioRender]

We then asked whether the global distribution of these SVs recapitulates established patterns of archaic ancestry. For each individual, we summed the length of candidate introgressed SVs and averaged these values within populations. Population-level SV length was strongly correlated with Neanderthal ancestry estimated from SNVs ^5^ (Spearman’s correlation coefficient ρ = 0.72; *p-value* = 7.27 × 10^-9^), and more modestly, but still significantly, with Denisovan ancestry (Spearman’s correlation coefficient ρ = 0.59; *p-value* = 1.25 × 10^-5^), likely reflecting the absence of Oceanian haplotypes from the graph genome (**Figs. S1-S3**).

As an independent test, we mapped Neanderthal and Denisovan sequences to the candidate introgressed insertions and compared their divergence with that of non-introgressed insertions (see Materials and Methods). Candidate introgressed insertions carried significantly fewer variants per bp than non-introgressed insertions for both Neanderthal-derived candidates (1.00 × 10^-4^ variants per bp in introgressed insertion vs. 3.02 × 10^-4^ variants per bp in non-introgressed insertion, *p-value* = 5.56 × 10^-8^) and Denisovan-derived candidates (3.27 × 10^-4^ vs. 6.27 × 10^-4^ variants per bp, *p-value* = 3.61 × 10^-3^); (**Fig. S4**). This reduced divergence supports closer sequence similarity to archaic homologs and is consistent with inheritance through introgression.

Several candidate introgressed SVs reached high frequencies (> 0.8) in specific present-day populations (**Fig. S5**), prompting us to test whether introgressed SVs are enriched for local adaptation. Using Ohana ^26^, a maximum likelihood framework that models a genome as mixtures of *K* ancestry components and tests for deviations from neutral allele frequency (AF) expectations, we found that introgressed SVs were depleted in the ancestry component maximized in African populations (*K* = 6) and enriched across multiple non-African ancestry components when modeled with eight components (**Figs. S6** and **S7**). Introgressed SVs were also significantly more likely to be under local adaptation than non-introgressed SVs (Fisher’s exact test, odds ratio = 2.49; *p-value* = 3.04 × 10^-7^; **Fig. 1C**), suggesting that archaic SVs have been a disproportionate source of adaptive variation in present-day humans.

We next examined the repeat composition and genomic context of the 153 candidate introgressed SVs. Most insertions (107 from Neanderthal and 54 from Denisovan) clustered around 300 bp in length (**Fig. S8a**), and were predominantly composed of *Alu* elements (> 64%), followed by simple repeats (> 14%) (**Fig. S8b**). Gene annotation showed that 72 Neanderthal-derived SVs and 23 Denisovan-derived SVs overlapped protein-coding genes (ENSEMBL), with > 90% mapping to introns (**Fig. 1D, E**). Two SVs were predicted to introduce coding frameshifts: a 54 bp Neanderthal-derived insertion (chr10-24594994-INS-54) in *ARHGAP21* and a 92 bp Denisovan-derived insertion (chr1-46270709-INS-92) in *RAD54L* (**Fig. S9**).

Together, these analyses define a set of SVs with multiple independent hallmarks of archaic introgression and show that introgressed SVs are enriched for loci under local adaptation. These results implicate archaic SVs as an important and underappreciated source of adaptive variation in local human populations.

### Denisovan-derived *Alu* insertion in *OCA2* is highly enriched in Bougainville Island of Melanesia

Among the 153 candidate introgressed SVs, we selected a Denisovan-derived *Alu* insertion in *OCA2* (chr15-27934219-INS-332) for in-depth analyses. We prioritized this SV for three reasons: it lies within a well-established pigmentation gene, shows strong population differentiation, and is present in the Denisovan genome but absent from great ape and African genomes. *OCA2* encodes the P protein, a transmembrane transporter central to melanogenesis ^27^. By regulating melanosomal pH and tyrosine levels, *OCA2* influences tyrosinase activity and eumelanin production ^27^. Given this biology, an introgressed regulatory variant in *OCA2* is a strong candidate to affect pigmentation-related phenotypes ^27^.

The variant is a 332 bp *AluY* element, identified by comparing the GRCh38 reference genome with a haplotype from HG00096, and is located in intron 18 of *OCA2* (chr15:27,934,218; **Fig. 2A**). Graph-based genotyping showed that the *Alu* insertion is homozygous in the Denisovan genome but absent from the Altai Neanderthal and great apes (**Figs. 2A** and **S10**). Analysis of four high-coverage archaic hominin genomes (Altai Neanderthal, Chagyrskaya Neanderthal, Denisovan, and Vindija Neanderthal) further supported a Denisovan-specific origin (**Fig. S11**).

**Figure 2.**
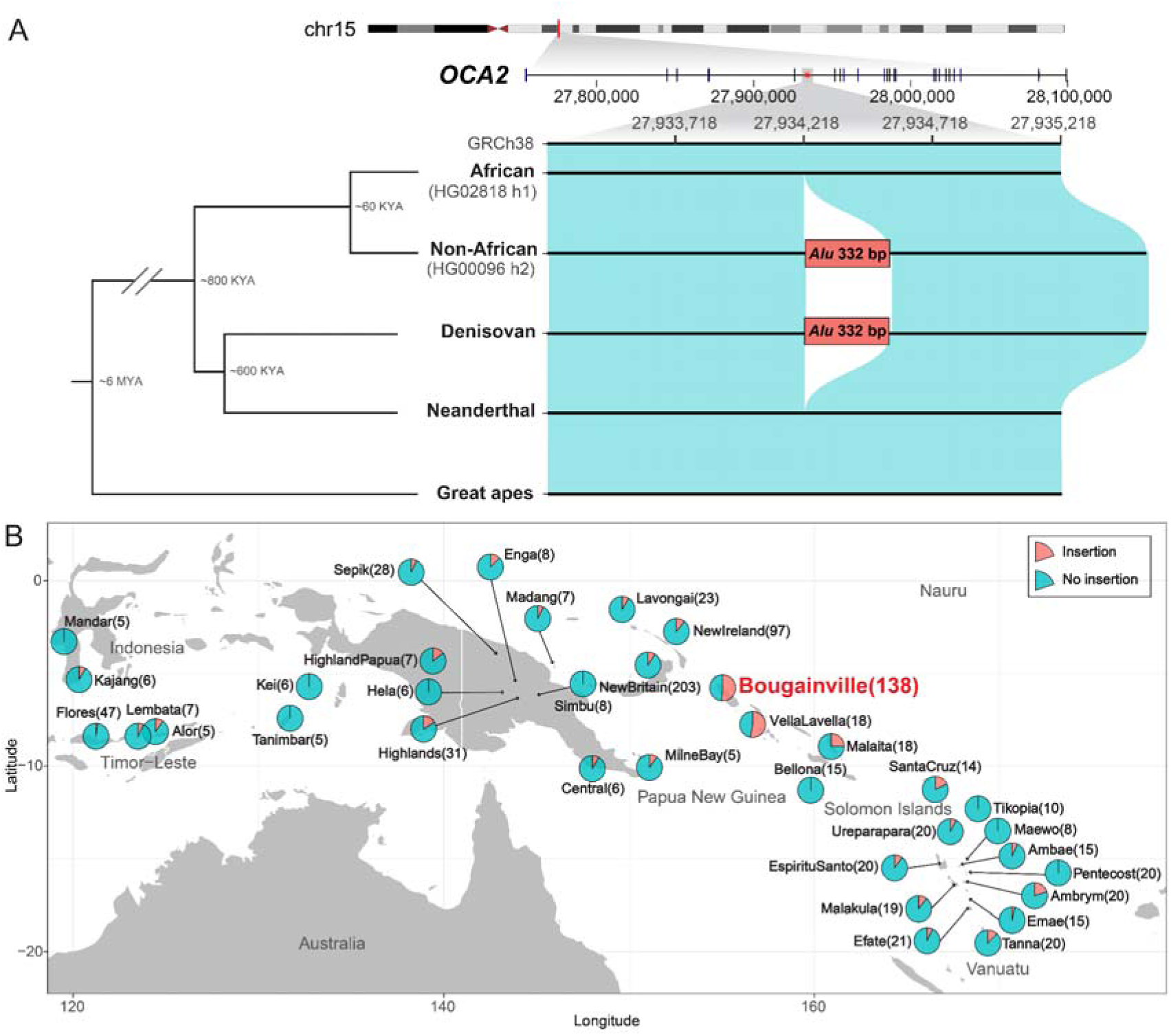
Geographic and population distribution of a Denisovan-derived *Alu* insertion in *OCA2* (chr15-27934219-INS-332; GRCh38). (**A**) Genomic context and population presence/absence. Schematic of the *OCA2* locus in the GRCh38 reference showing the position of the *Alu* insertion (chr15:27,934,219; INS-332) and whether the *Alu* insertion is detected in each population group. (**B**) Regional allele frequency map of the *OCA2 Alu* insertion around Bougainville Island. Pie charts show *Alu* insertion allele frequency (red) for populations sampled in and around Bougainville Island using data from the Human Genome Diversity Project (HGDP; n=828), the Southeast Asia and Oceania dataset (SAO; n=586), and genotyping PCR results (n=426). Individuals from Bougainville across the three datasets were consolidated into a single “Bougainville” group. To harmonize labels across datasets, East Sepik (SAO), West Sepik (SAO), and Papuan Sepik (HGDP) were merged into the “Sepik” group, while Eastern Highlands (SAO), Western Highlands (SAO), and Papuan Highlands (HGDP) were combined into the “Highlands” group. Sample sizes for each region are indicated in parentheses. Regions with fewer than five individuals are not shown. For sequence-based genotypes, calls with genotype quality scores (GQ) < 30 were excluded before calculating allele frequencies.

Using data from the 1000 Genomes Project (1KGP) and the Human Genome Diversity Project (HGDP), we found that the *OCA2 Alu* insertion reaches an AF of 0.64 in the Nasioi population of Bougainville Island (4 homozygotes and 6 heterozygotes among 11 individuals), while averaging only 0.056 across 61 other non-African populations (14 homozygotes and 289 heterozygotes among 2,572 individuals) and is absent from East Asia (**Fig. S12**). When filtering 33,828 SVs for those rare in Africans (AF < 0.01) and non-Africans (AF < 0.1) but common in the Nasioi (AF > 0.5), only 18 variants (0.05%) met these criteria. Among them, the *OCA2 Alu* insertion was the only SV absent in great apes but present in the Denisovan genome. Genotyping 586 individuals from 57 regions in and around Bougainville Island shows a similar geographic pattern. The *Alu* insertion was enriched on Bougainville (AF = 0.50; 2 homozygotes and 1 heterozygote among 5 individuals) and on nearby Vella Lavella Island (AF = 0.53; 5 homozygotes and 9 heterozygotes among 18 individuals), with AF declining as geographic distance from Bougainville increases (**Table S2, Figs. 2B** and **S13**).

Overall, these results identify a putatively Denisovan-derived *Alu* insertion in *OCA2* that has reached unusually high frequency in the Bougainville region. Since such strong geographic enrichment can arise through multiple evolutionary processes, including demographic history and positive selection, we next investigated the evolutionary history of this locus to understand forces shaping its present-day distribution.

### Evolutionary history of the *OCA2 Alu* insertion

To further test whether the *OCA2 Alu* insertion resulted from archaic introgression, we compared its location with known Denisovan-like segments ^28^. The *OCA2 Alu* insertion is found in 164 Denisovan-like introgressed segments with high posterior probability (> 0.8). These segments are much more common among individuals with the *Alu* insertion than in those without it (*X*^2^-contingency test *p-value* = 1.80 × 10^-132^). This pattern suggests the *Alu* insertion is part of an introgressed haplotype from Denisovan.

We then investigated the haplotype structure around 100 kb of the *Alu* insertion using phased genotypes from both present-day and archaic individuals. The *Alu* insertion is located in a recombination hotspot, and its strongest LD (r^2^ = 0.48) is with an SNV located 515 bp downstream at chr15:27,934,733 (chr15-27934733-SNV-T-C; **Fig. 3A**). Because of the high recombination rate, no nearby SNV had an r^2^ > 0.50 with the *Alu* insertion. This indicates that no single marker can reliably represent the *Alu* insertion haplotype, leading us to haplotype-based analyses.

**Figure 3.**
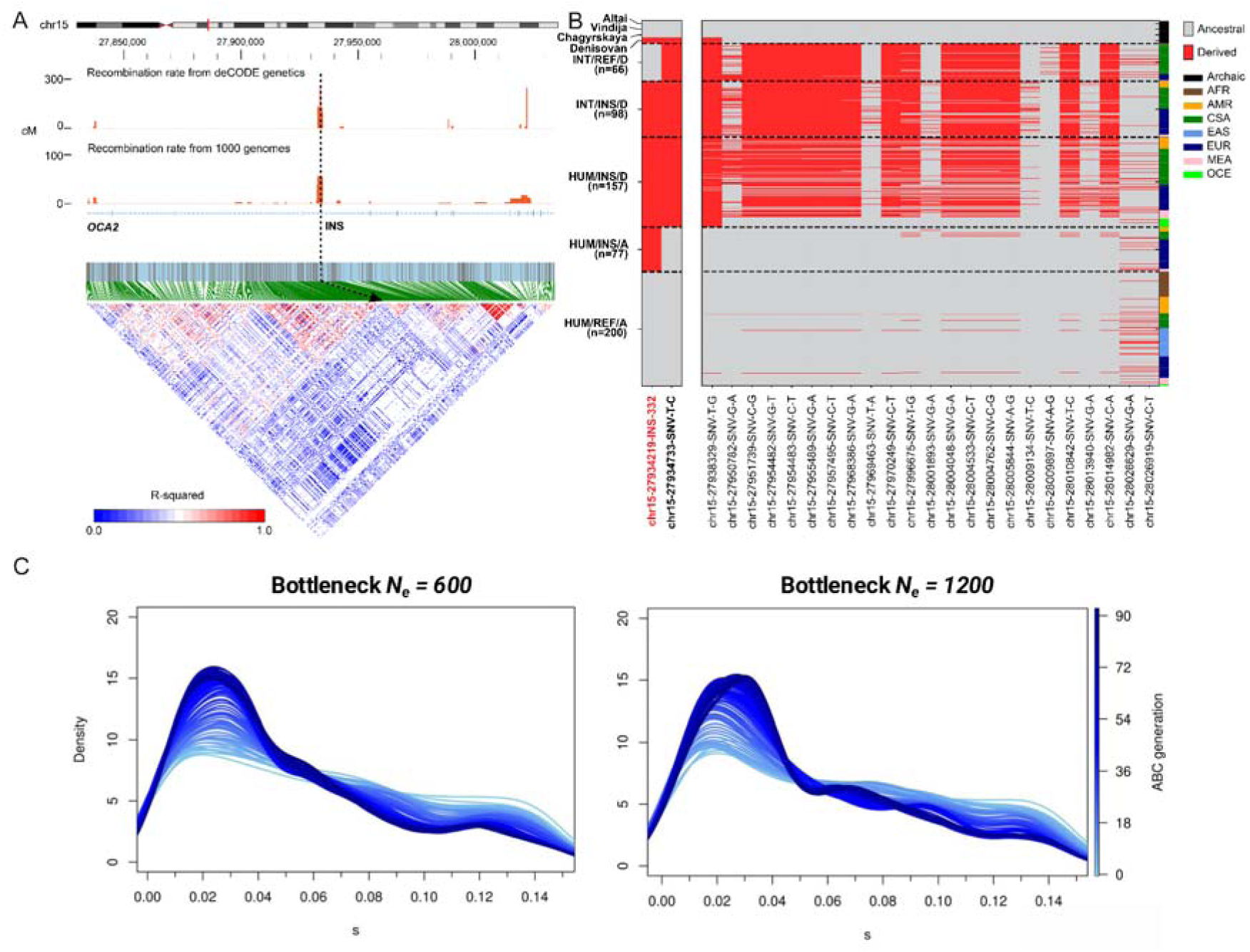
Haplotype structure and selection around the Denisovan-derived *Alu* insertion in *OCA2*. (**A**) Local recombination landscape (top) and linkage disequilibrium (LD; bottom) across 100 kb of the *Alu* insertion site. Recombination rates were obtained from the deCODE Genetics and 1000 Genomes Project via the UCSC Genome Browser. The *Alu* insertion lies within a recombination hotspot and shows stronger LD with variants on the downstream of the locus than the upstream. (**B**) Phased genotype matrix at the *Alu* insertion and 24 single-nucleotide variants (SNVs) identified by hmmix ^28^ as informative of introgression. Haplotypes carrying the *Alu* insertion show elevated divergence and are most strongly associated with the downstream SNV, chr15-27934733-SNV-T-C (r² = 0.48). For visualization, 200 haplotypes lacking the *Alu* insertion were randomly sampled from present-day humans (HUM/REF/A). All insertion-carrying haplotypes were included. (**C**) Selection coefficient (s) density from Approximate Bayesian Computation (ABC) simulations with severe (N_e_ = 600, left) and moderate (N_e_ = 1200, right) bottlenecks in the Nasioi population of Bougainville Island.

We focused on 25 sites informative for introgression, including SNVs identified by hmmix ^28^ and the *Alu* insertion itself, and compared haplotypes from present-day and archaic hominin genomes. Haplotypes with the *Alu* insertion were grouped with those predicted to be Denisovan-like and were different from most present-day haplotypes without the *Alu* insertion (**Fig. 3B**). Only 3 of the 25 informative sites, including the *Alu* insertion itself, match the Denisovan allele. However, empirical resampling and forward simulations showed that this limited allele sharing is expected for Denisovan-like segments in this region (**Fig. S14**).

Although many insertion haplotypes matched those found in the Denisovan-like haplotypes, the match was not perfect. Some individuals with the *Alu* insertion did not show signs of Denisovan ancestry, and some Denisovan-like haplotypes lacked the *Alu* insertion (**Fig. 3B**). We first ruled out genotyping errors by visual inspection of read alignments. We then sorted haplotypes into five groups based on Denisovan ancestry (INT; Introgression vs. HUM; Human), whether the *Alu* insertion was present (INS; Insertion vs. REF; Reference), and the allele state at the SNV with the highest LD (D; Derived vs. A; Ancestral): HUM/REF/A, (n=200), HUM/INS/A (n=77), HUM/INS/D (n=157), INT/INS/D (n=98), INT/REF/D (n=66). In a haplotype network, HUM/REF/A clustered with HUM/INS/A, while HUM/INS/D clustered with INT/REF/D, INT/INS/D, and the Denisovan haplotypes (**Fig. S15**). This pattern at first seemed to be consistent with recurrent insertions. However, all insertion haplotypes share identical breakpoints in read alignments (**Fig. S16**), arguing against multiple independent insertions. Furthermore, in the marginal tree at chr15-27934733-SNV-T-C, HUM/INS/A haplotypes were scattered across the tree (**Fig. S17**), which does not fit with recurrent insertion events. Along with the geographic distribution of the *Alu* insertion, these findings support a model in which Denisovan introgression was followed by rapid erosion of the surrounding haplotype due to the high local recombination.

We then examined whether the high frequency of the *Alu* insertion in the Nasioi population (AF = 0.64) was shaped by positive selection or neutral demographic processes. Chromosome-wide normalized XP-EHH did not show a strong signal at the insertion site (mean *XP-EHH(CSA, Nasioi)* = -0.66 and mean *XP-EHH(EAS, Nasioi)* = 0.65; **Fig. S18a**, **c**). However, after adjusting XP-EHH for local recombination rate, the insertion site stood out as an outlier when Central and South Asian (CSA) populations were used as a reference (mean *XP-EHH_rec_(CSA, Nasioi)* = -3.55; **Fig. S18b**), but not when using East Asian (EAS) populations as a reference (mean *XP-EHH_rec_(EAS, Nasioi)* = 0.29; **Fig. S18d**). This suggests that rapid local recombination may partly mask any haplotype-based selection signal. To further test this observation and estimate the strength of selection, we used Approximate Bayesian Computation (ABC) with several summary statistics (see Materials and Methods). In 15,000 simulations with selection coefficients from 0.0 to 0.15, the results supported positive selection with a median selection coefficient of 0.0363 (95% CI: 0.0027 - 0.1385; **Fig. 3C**). These estimates account for local recombination rates and were similar under both severe (N_e_ = 600) and moderate (N_e_ = 1,200) bottleneck models for the Nasioi population.

In summary, several independent analyses support the idea that the *OCA2 Alu* insertion was introgressed from Denisovans and that the surrounding haplotype has been broken down by recombination. Also, the combined haplotypic, geographic, and recombination-aware selection analyses best fit a model of Denisovan introgression followed by positive selection in Bougainville Island, which led us to test the effects of the *Alu* insertion on pigmentation traits and gene regulation.

### Denisovan-derived *OCA2 Alu* insertion is associated with skin pigmentation variation in Bougainville Island

To assess whether the *OCA2 Alu* insertion is linked to pigmentation-related traits, we first examined published genome-wide association study (GWAS) data for the 24 introgression-informative SNVs surrounding the *Alu* insertion (**Fig. 4A**). Twenty of these SNVs have previously been associated with pigmentation traits (**Table S3**). The SNV in strongest LD with the *Alu* insertion, chr15-27934733-SNV-T-C (r^2^ = 0.48), is associated with multiple pigmentation traits, including “ease of skin tanning” (*p-value* = 1.34 × 10^-11^), “Blonde hair color” (*p-value* = 9.90 × 10^-9^), “Skin color” (*p-value* = 8.08 × 10^-8^), and “Light brown hair color” (*p-value* = 3.13 × 10^-4^) (**Fig. 4B**). These observations indicate that the introgressed haplotype lies in a pigmentation-relevant regulatory landscape.

**Figure 4.**
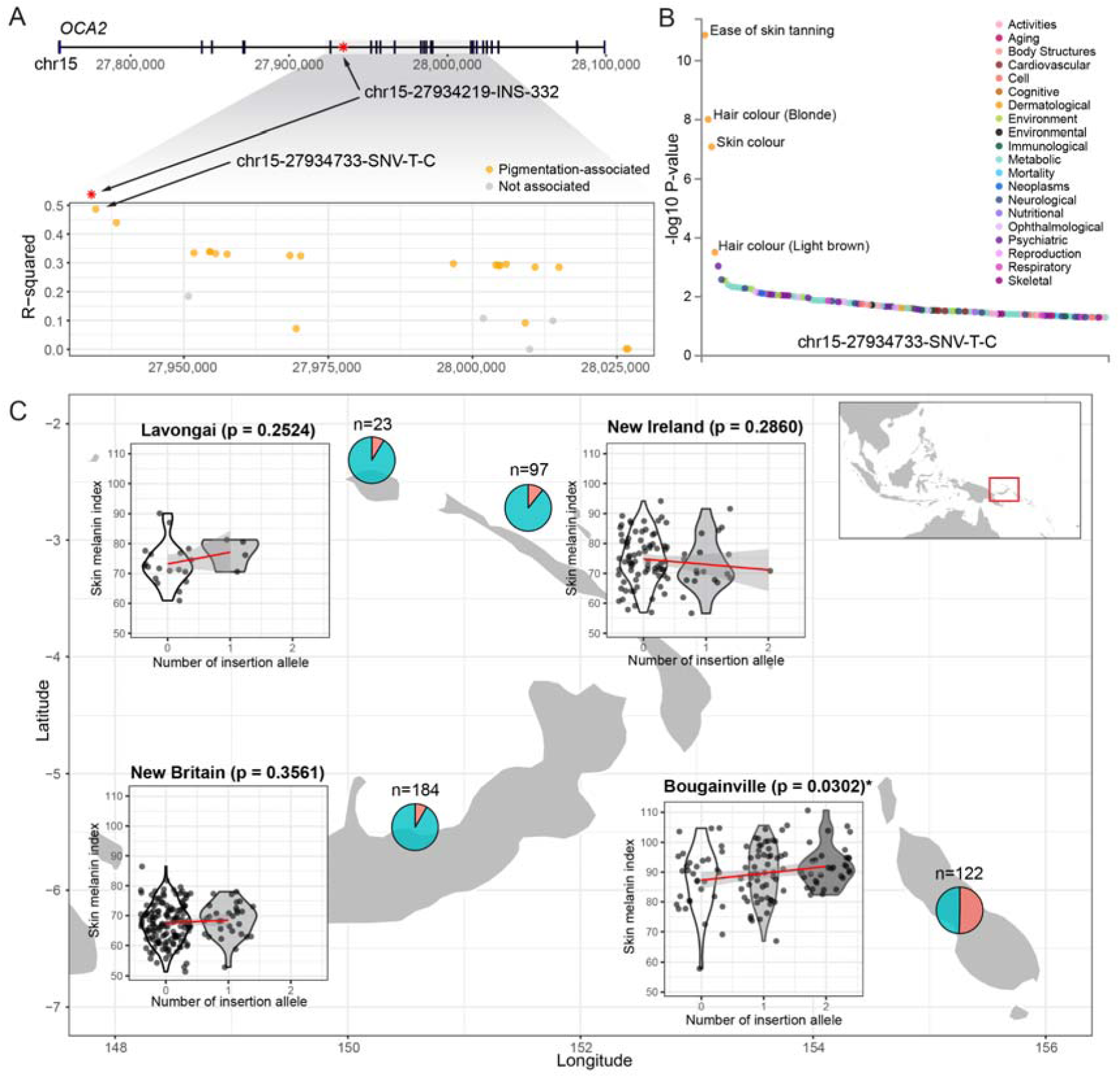
Association of the Denisovan-derived *Alu* insertion in *OCA2* with skin pigmentation in Bougainville Island. (**A**) Gene structure of *OCA2* (top) and pairwise linkage disequilibrium (LD; r²) between the *Alu* insertion and 24 introgression-informative variants (bottom). Variants previously reported to be associated with pigmentation traits are shown in orange. Variants with no reported pigmentation association are shown in gray. (**B**) Strength of association with published phenotypes for the introgression-informative variant in the strongest LD with the *Alu* insertion (chr15-27934733-SNV-T-C, r^2^ = 0.48). Data points corresponding to hair or skin color phenotypes are highlighted in orange with corresponding labels. (**C**) Association between insertion allele dosage (0-2 copies) and skin melanin index across four islands in and around Bougainville Island. The red line shows the smoothed trend (ggplot ^104^ using the formula = y ∼ x). The pie chart for each island indicates the *Alu* insertion allele frequency for each island (red = *Alu* insertion allele).

We next tested the association between the *Alu* insertion genotype and measured pigmentation in populations from the Bougainville region. We genotyped 426 individuals from four islands: Bougainville (n = 122), Lavongai (n = 23), New Britain (n = 184), and New Ireland (n = 97), for whom skin and hair melanin measurements were available ^29,30^. The *Alu* insertion allele showed a strikingly uneven geographical distribution: it was common on Bougainville (AF = 0.50), but rare on the other islands (maximum AF = 0.11), and only one individual outside Bougainville was homozygous for the *Alu* insertion (**Fig. 4C**). Skin melanin index varied considerably between islands (F = 129.21; *p-value* = 2.56 × 10^-26^), primarily driven by the high melanin index observed in Bougainville. Hair melanin index, however, did not differ much between islands (F = 0.57; *p-value* = 0.45; **Fig. S19**).

On each island, we tested whether the number of *Alu* insertion alleles (i.e., insertion dosage) was associated with melanin levels. We used two models: a simple linear model with insertion dosage as the predictor, and a full model that also included insertion x age and insertion x sex interactions (**Table S4**). In Bougainville, the simple model showed a significant association between insertion dosage and skin melanin index, with an increase of 2.26 melanin units per insertion allele (*p-value* = 0.0302; **Fig. 4C**). No significant association was observed for hair melanin index (*p-value* = 0.6453; **Fig. S20**). In the full model, the main effect of insertion dosage on skin melanin index was not significant (*p-value* = 0.0848), but the interaction between insertion and sex was significant (*p-value* = 0.0085), with a stronger effect in females than in males (**Table S4** and **Fig. S21**), consistent with the sex-specific effects occasionally observed for pigmentation traits ^31^. We did not find significant associations for skin or hair melanin indices in Lavongai, New Britain, or New Ireland islands in either model, except for the insertion x age interaction in New Ireland (**Table S4**).

Overall, these results link the Denisovan-derived *OCA2 Alu* insertion to a higher skin melanin index in Bougainville Island. The geographic restriction of this effect, together with the evidence for sex-specific differences, suggests that the phenotypic impact of the *Alu* insertion depends on local genetic or environmental context.

### Denisovan-derived *Alu* insertion increases *OCA2* expression and pigmentation in human melanocytes

To test whether the Denisovan-derived *Alu* insertion affects *OCA2* function, independent of genetic background, we introduced a single copy of the 332 bp *Alu* element into the KOLF2.1J iPSC line ^32^, generating heterozygous (HET) edited cells. Edited and wild-type (WT) cells derived from the same parental line were then differentiated into melanocytes (**Fig. 5A**). We measured gene expression at seven time points during melanocyte development. HET melanocytes consistently showed higher *OCA2* expression than WT in independent qPCR assays, with differences seen as early as day 16 (**Fig. 5B**). By contrast, melanocyte marker genes, including *MiTF*, were expressed at similar levels in both genotypes, indicating that cells developed similarly and that the effect was specific to *OCA2.* Consistent with the established relationship between *OCA2* expression and melanin production ^27^, HET melanocytes also showed visibly greater pigmentation than WT (**Fig. 5C**). Together, these results indicate that the Denisovan-derived *Alu* insertion enhances melanogenesis by upregulating *OCA2*.

**Figure 5.**
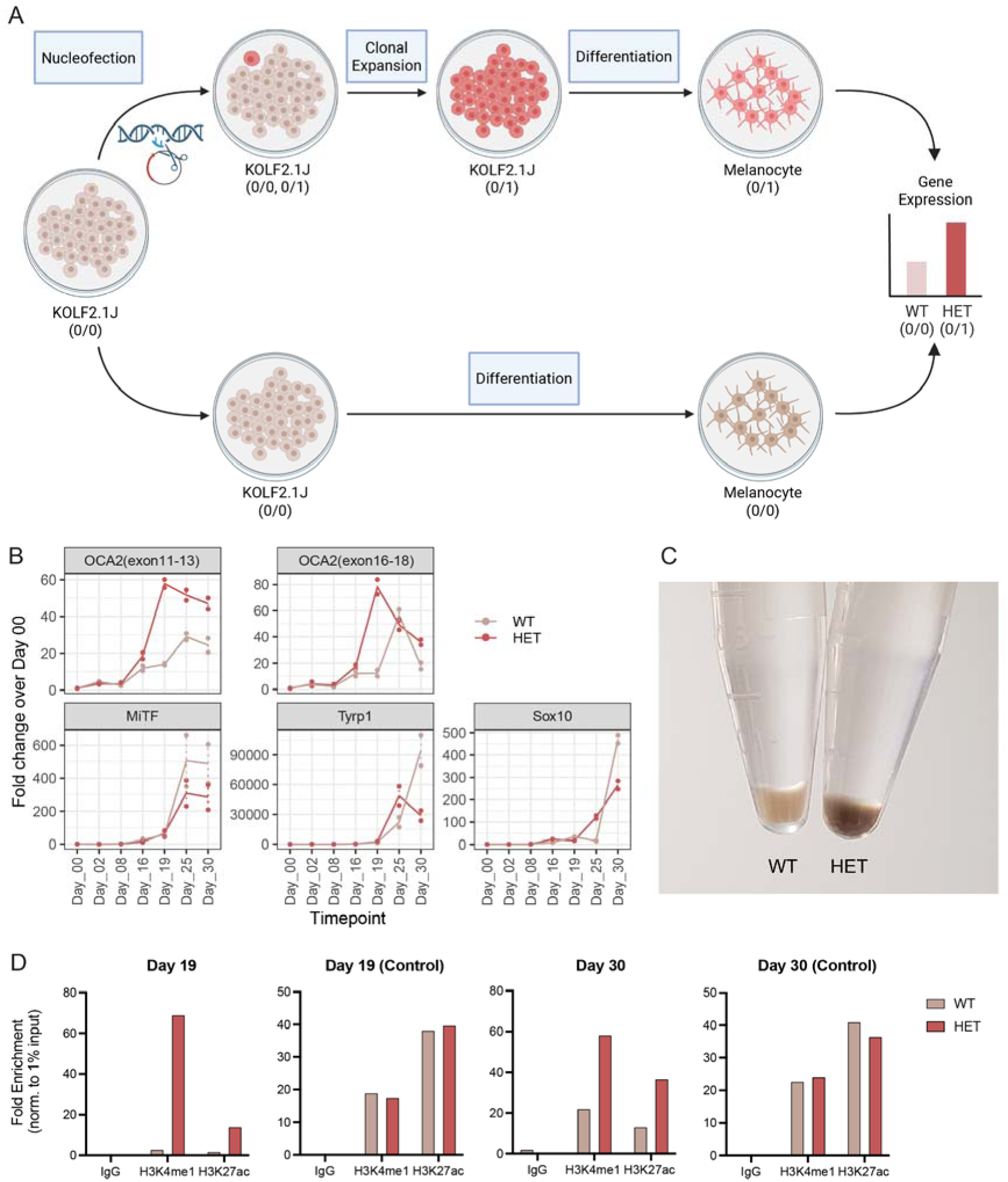
Functional effects of the Denisovan-derived *Alu* insertion in *OCA2.* (**A**) Schematic overview of the *in vitro* experimental design to test the *Alu* insertion’s functional impact in the human induced pluripotent stem cell (iPSC) line, KOLF2.1J. (**B**) Expression of *OCA2* and melanocyte marker genes across seven stages of melanocyte differentiation (days 0, 2, 8, 16, 19, 25, and 30). Each dot represents a technical replicate (heterozygous insertion carriers, red; wildtype, gray). Lines indicate the mean across two replicates at each time point. Two *OCA2* amplicons are shown: exon 11-13 and exon 16-18 (exon numbering from RefSeq NM_000275.3 as displayed in the UCSC Genome Browser). (**C**) Representative image illustrating pigmentation differences between wild-type (WT; homozygous reference) and heterozygous (HET; one *Alu* insertion allele) melanocyte cultures. (**D**) Enrichment of histone marks, H3K4me1 and H3K27ac, at days 19 and 30. “IgG” and “Control” indicate negative controls for the enrichment assay and locus-specific signal, respectively. The locus “Control” is derived from the *B2M* gene. Fold enrichment values were normalized to 1% of the input control. [Figure created with BioRender]

To understand how the *Alu* insertion increases *OCA2* expression, we considered two possible explanations: it could change splicing or affect cis-regulatory activity. RNA□seq showed higher *OCA2* expression in HET melanocytes compared to WT (**Fig. S22**), but there were no exon-specific changes that would suggest altered splicing (**Fig. S23**). This indicates that the *Alu* insertion likely does not change the transcript structure, but instead affects overall transcriptional regulation. We then checked if the *Alu* insertion lies in a regulatory element for melanocytes. Public epigenomic data from ChIP-Atlas ^33^ showed that the *Alu* insertion overlaps a region marked by H3K27ac in melanocyte-related cells (i.e., 501mel, CHL1, and NHEM; **Fig. S24**), suggesting enhancer-like activity in pigment cells. To test this directly, we performed ChIP-qPCR for histone marks associated with active enhancers. Both H3K4me1 and H3K27ac enrichments were significantly elevated at the *Alu* insertion site in HET melanocytes compared to WT, with fold enrichment differences up to 66. The pattern of H3K4me1 enrichment matched the expression results, with a higher difference at day 19 than at day 30 (**Figs. 5D** and **S25**). These results support the idea that the *Alu* insertion boosts local enhancer activity, thereby increasing *OCA2* expression and melanocyte pigmentation.

In summary, population genetic analyses and functional assays converge on the conclusion that the Denisovan-derived *Alu* insertion in *OCA2* contributes to the higher skin pigmentation in Bougainville Island by elevating *OCA2* expression, offering a rare example of introgressed SVs with a measurable phenotypic impact.

## DISCUSSION

Archaic introgression has emerged as an important source of genetic variation in modern humans, but most work in this area has focused on SNVs ^1–3^. By contrast, the contribution of archaic hominin-derived SVs, particularly to present-day human phenotypes, has remained largely unexplored. Using graph genome-based genotyping, this study systematically identified 153 candidate SVs inherited from Denisovans or Neanderthals. Most of these candidate introgressed SVs were insertions (n = 143), with *Alu* elements dominating (87/143; **Figs. S8b** and **S26b**), suggesting that transposable element variation is an important but understudied source of archaic introgression. Among these candidates, we identified a 332 bp Denisovan-derived *AluY* insertion in intron 18 of *OCA2*, a key pigmentation gene, that reaches unusually high frequency in Bougainville Island and is associated with increased skin pigmentation in that region. In melanocytes derived from edited human iPSCs, this *Alu* insertion led to increased *OCA2* expression, greater pigmentation, and higher levels of enhancer-associated chromatin marks at the insertion site. These findings implicate introgressed SVs as a functional source of phenotypic variation in present-day humans and identify a Denisovan-derived regulatory insertion in *OCA2* as a contributor to pigmentation diversity in Melanesian populations (**Fig. 6**).

**Figure 6.**
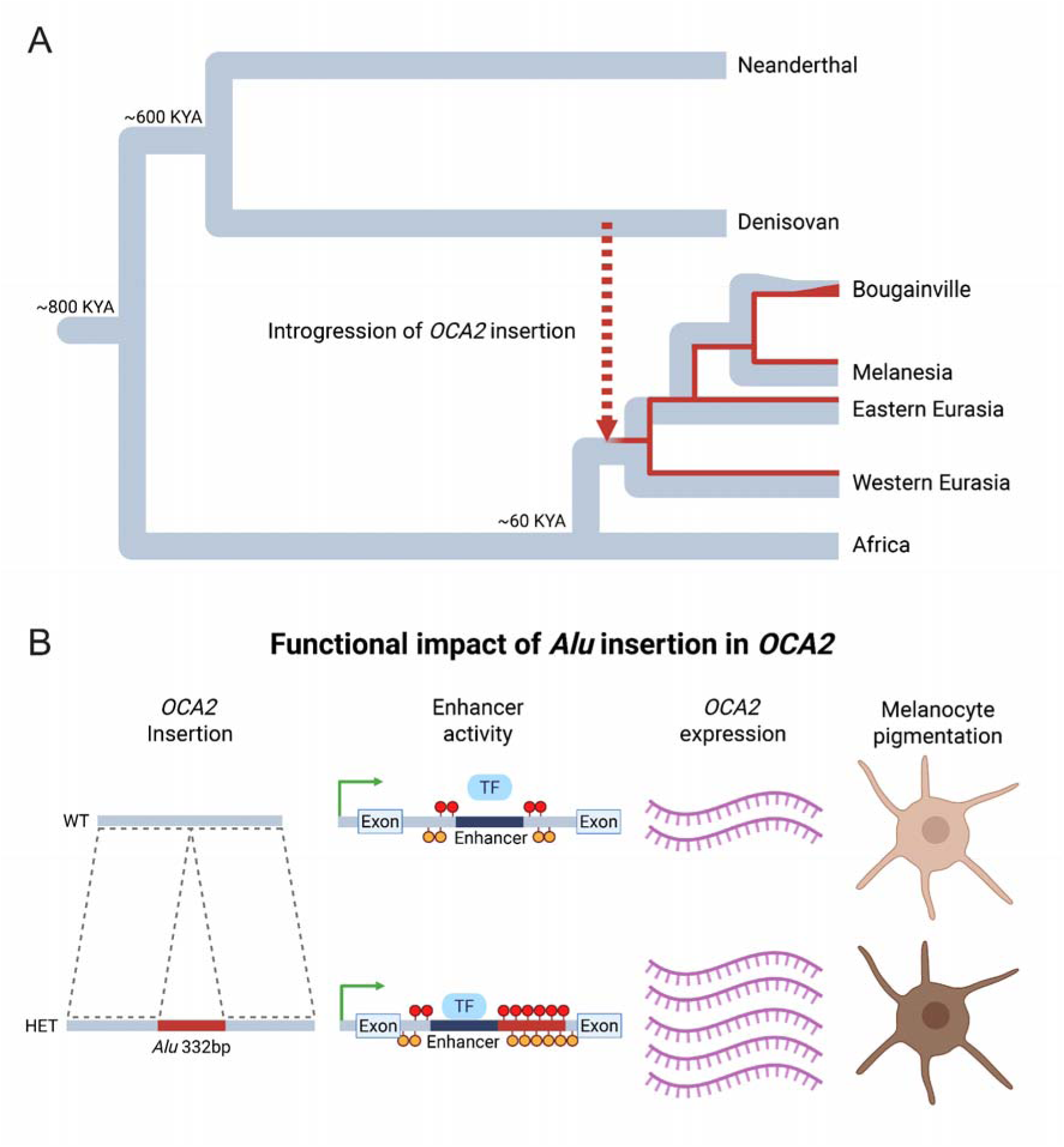
Model of the Denisovan-derived *Alu* insertion in *OCA2*: Evolutionary history and functional impact. (**A**) Schematic phylogeny summarizing the inferred origin and spread of the Denisovan-derived *Alu* insertion in *OCA2*. The *Alu* insertion is modeled as existing in Denisovans and introgressing into non-African populations. Present-day allele frequency differences (particularly on Bougainville Island) are consistent with effects of local demographic history and possible positive selection. (**B**) Summary of functional consequences of the Denisovan *Alu* insertion in *OCA2* based on *in vitro* experiments. [Figure created with BioRender]

A primary challenge in interpreting this locus is that the *Alu* insertion occurs within a recombination hotspot. This led to only moderate linkage with nearby SNVs (r² = 0.48 at the best proxy) and indicates that carrying the *Alu* insertion does not always match up with introgression labels. Nevertheless, multiple independent lines of evidence indicate a Denisovan origin for the *Alu* insertion: it appears in the Denisovan genome but not in Africans, Neanderthals, or great apes; haplotypes carrying the *Alu* insertion cluster with Denisovan-like haplotypes; and all carriers exhibit identical breakpoints, consistent with a single historical insertion event. Collectively, these results support a model in which the *Alu* insertion entered present-day humans via Denisovan introgression, followed by rapid local recombination that eroded the surrounding haplotype.

Our data also provide initial insight into the molecular mechanism by which this introgressed SV may act. The *Alu* insertion is located in an intronic region that already shows enhancer□associated chromatin marks (H3K27ac) in melanocyte-related datasets, even in the absence of the *Alu* insertion. This suggests the locus already had regulatory potential. *Alu* elements themselves can develop enhancer□like chromatin states and alter transcription factor binding sites ^34^, providing a plausible route by which the *Alu* insertion could modulate local gene regulation. Consistent with this idea, the *Alu* insertion increased H3K4me1 and H3K27ac enrichment at the locus in edited melanocytes, accompanied by higher *OCA2* expression and increased pigmentation. These changes were most noticeable around day 19 of differentiation ^35^, when neural crest-derived precursors are transitioning into pigment□producing melanocytes. Overall, these findings suggest that the Denisovan-derived *Alu* insertion strengthens an existing regulatory element, thereby increasing *OCA2* expression during melanocyte development.

The association analyses suggest that the phenotypic effects of the *Alu* insertion are context dependent. In Bougainville, individuals carrying more copies of the *Alu* insertion tend to have increased skin melanin levels. In contrast, little to no effect was observed on nearby islands, where the allele is rarer, and the genetic background may differ. The association in Bougainville was also stronger in females in models with interaction terms. These results, however, should be interpreted cautiously, as they may be influenced by factors beyond the sex and age we accounted for, including differences in sample size, allele frequency, environmental exposure, and unmeasured genetic effects. Nonetheless, sex-specific effects have been reported for certain pigmentation□related traits ^31^, and our results raise the possibility that the impact of the *OCA2* regulatory variant differs across demographic or physiological contexts. More broadly, they underscore that the phenotypic consequences of introgressed variants may depend not only on the variant itself, but also on the population in which it occurs.

The unusually high frequency of the *Alu* insertion in Bougainville raises a question of whether this geographic concentration reflects positive selection, random genetic drift, or both. Recombination□adjusted XP-EHH and ABC analysis support the idea that positive selection acted on the *Alu* insertion in Bougainville Island, but the strength and timing of this selection depend on assumptions about the population and on uncertainty about the local recombination rate. The environmental cause is also unclear: ultraviolet radiation is about the same across nearby islands at similar latitudes ^29^. Instead, the unusually high AF of the *Alu* insertion on Bougainville is likely the outcome of positive selection acting in combination with demographic factors, including founder effects, drift, and long□term population isolation ^36,37^. Given that the effect of *Alu* insertion likely interacts with the genetic background in which it lies, a plausible scenario is that the *Alu* insertion entered the region at low frequency, gained an early foothold in Bougainville due to stochastic demographic events, and then increased under selection due to its stronger functional effect than elsewhere. Denser regional sampling and more explicit demographic modeling will be needed to fully resolve the evolutionary history of this locus.

More generally, some Melanesian populations are known for having a striking phenotype: an increased skin pigmentation combined with naturally blonde hair, found in up to 10% of individuals in some groups ^29^. While our results identified a contributor to increased skin pigmentation, they do not explain the occurrence of blonde hair in Bougainville Island. Previous work showed that a *TYRP1* missense variant (rs387907171) is a major determinant of blonde hair in the Solomon Islands ^38^, but this allele is absent in Bougainville Island, in both our data (**Fig. S27**) and a prior study ^39^, despite the presence of light hair color on the island ^29^. This discrepancy indicates that additional, yet unidentified, variants contribute to hair color variation in Bougainville, potentially through interaction with *OCA2* or other pigmentation loci. Identifying these variants will deepen our understanding of how pigmentation traits evolved in genetically distinct Melanesian populations.

Our study also provides a rare functional glimpse into Denisovan biology ^40^. Introgressed variants preserved in present-day human genomes can serve as molecular traces of archaic hominin phenotypes, particularly when direct physical remains are limited. Phenotypic predictions from Denisovan SNVs have suggested that Denisovans had brown eyes, brown hair, and increased skin pigmentation ^41–43^. The observation that the Denisovan-derived *OCA2 Alu* insertion increases pigmentation in human melanocytes is consistent with the possibility that it contributed to elevated pigmentation in Denisovans themselves. Although such inferences remain indirect, they illustrate how functional analysis of introgressed variants, especially SVs, can complement comparative genomics and recent genome-editing studies that experimentally model archaic alleles in human cells ^44–47^.

Finally, our results highlight the value of studying archaic introgression through an SV-centered lens. The current capacity to detect introgressed SVs remains limited by the restricted diversity in long-read SV catalogs and technical challenges associated with graph-based genotyping. These constraints are particularly relevant for Oceanian populations, which are underrepresented in existing genomic resources. Expanding pangenome references ^48^ to include more individuals from Oceania and other understudied regions will improve the discovery of archaic-derived SVs, refine their evolutionary histories, and broaden the set of candidates available for functional study. As these resources mature, SV-focused approaches should provide a more complete picture of how archaic introgression has shaped present-day human diversity and offer new insight into the biology of our extinct relatives.

## MATERIALS AND METHODS

### Processing of raw sequence reads

High coverage (∼30x) raw sequencing reads for archaic hominin genomes were obtained from three previously published studies: Altai Neanderthal ^49^, Vindija Neanderthal ^50^, and Denisovan ^41^ (**Table S5**). To mitigate the effects of ancient DNA damage on k-mer-based analyses, we first evaluated the damage profiles of all sequencing libraries (**Fig. S26a**). Initial quality control was performed using FastQC v0.11.9 (https://www.bioinformatics.babraham.ac.uk/projects/fastqc/). Low-quality bases (Phred base quality score < 20) and adapter sequences were removed using AdapterRemoval v2.3.2 ^51^ with the “--collapse” option. For each library, one million reads were randomly selected and aligned to the GRCh37 reference genome (hs37d5) using bwa aln v0.7.17 ^52^ with the parameters “-n 0.01”, “-l 16500”, and “-o 2”. PCR duplicates were identified and removed using the Picard MarkDuplicates v2.26.11 (http://broadinstitute.github.io/picard). DNA damage profiles were generated using Damageprofiler v1.1 ^53^ using reads with a mapping quality score ≥ 30. Of 27 libraries analyzed, 16 exhibited characteristic ancient DNA damage patterns, namely, elevated C → T or G → A mismatch within 5 bp of read termini. To reduce the influence of damage on k-mer genotyping, we trimmed 5 bp from both ends of reads in these libraries using fastp v0.23.2 ^54^ with the deduplication (--dedup) and minimum read length of 31 bp (-l 31) options. Libraries showing abnormal mismatch rates within internal read regions (B1107 and B1109 for Denisovan, and A9368, A9369, A9401, A9402, A9403, A9404, B8747, and R5473 for Vindija Neanderthal) were excluded. The remaining libraries for each archaic hominin genome were subsequently merged into a single FASTQ file per individual. Sequencing reads from present-day humans (**Table S5**), excluding samples from the 1KGP dataset, were processed analogously (**Fig. S26a**). Reads were quality filtered and deduplicated using fastp v0.23.2 ^54^, and high-quality reads were merged into individual FASTQ files.

### Structural variant genotyping using PanGenie

SVs and other variants were genotyped using PanGenie v1.01 ^55^, a k-mer-based genotyping framework, with default parameters. Inputs included: (i) the graph genome from the Human Genome Structural Variation Consortium (HGSVC) ^24^, (ii) preprocessed high-quality reads, and (iii) the human reference genome, GRCh38 (excluding alternate haplotypes; http://ftp.1000genomes.ebi.ac.uk/vol1/ftp/data_collections/HGSVC2/technical/reference/20200513_hg38_NoALT/hg38.no_alt.fa.gz) (**Fig. S26a**). PanGenie successfully genotyped 95.83%, 95.82%, and 92.77% of variants in the graph genome for Altai Neanderthal, Denisovan, and Vindija Neanderthal, respectively. For 1KGP samples, publicly available PanGenie genotype calls were incorporated (https://ftp.1000genomes.ebi.ac.uk/vol1/ftp/data_collections/HGSVC2/release/v2.0/PanGenie_results/20201217_pangenie_merged_bi_all.vcf.gz). Across archaic and present-day samples, we genotyped a total of 60,418 insertions (≥50 bp) and 35,859 deletions (≥ 50 bp). Locus-wise filtering was applied using HGSVC “lenient” criteria ^24^, retained genotypes with quality scores (GQ) > 30, and excluded loci with a missing genotype rate > 0.05. The resulting dataset comprised 30,889 insertions (≥ 50 bp) and 19,962 deletions (≥ 50 bp), 748,812 small indels (< 50 bp), and 12,328,706 SNVs with a mean genotyping rate of 99.84% (**Fig. S26b**).

Additionally, we analyzed 586 genomes from Southeast Asian and Oceanian populations (PRJEB9586, EGAD00001001634, EGAD00001004156, EGAD00001007783, and EGAD00001006880, **Tables S2** and **S5**), using the same PanGenie pipeline.

### Genotype quality assessment

To benchmark the quality of archaic genome genotypes relative to present-day genomes, we randomly selected 10 individuals each from 1KGP, HGDP, and great ape datasets. For each sample, we examined k-mer distributions and quantified k-mer support per variant. The Vindija Neanderthal genome exhibited a high proportion of erroneous k-mers (41.86%) and lower genotype quality (**Figs. S28** and **S29**). Consequently, it was excluded from downstream analyses.

Further evaluation included transition-to-transversion (Ts:Tv) ratios and heterozygosity estimates. As previously reported ^56^, Ts:Tv ratios for SNVs were approximately 2.1 across present-day humans and archaic hominins, while great ape genomes displayed slightly elevated Ts:Tv ratios (**Fig. S30**). Consistent with prior studies ^41,49,50^, heterozygosity was reduced in the Altai Neanderthal, Denisovan, and great ape compared to present-day humans (**Fig. S31**).

### Identification of introgressed structural variants

To identify SVs introgressed from archaic hominins, we applied a three-step filtering (**Fig. S26a**). First, we selected SV alleles absent in all great ape genomes but present as a homozygous state in at least one archaic hominin genome. Second, we excluded SVs present in African populations without recent admixture (i.e., Bantu Kenya, Bantu South Africa, Biaka, Esan, Luhya, Mandinka, Mbuti, Mende, San, and Yoruba, n=592), assuming that these African populations have no introgression from archaic hominins. Finally, we retained SVs present in at least one non-African individual.

### Variant calling within non-reference insertions

To support the notion that candidate SVs were introgressed from archaic hominins rather than represent ancestral variants, we compared sequence divergence between archaic hominins and non-African present-day humans. High-quality archaic reads (without internal mismatches) were aligned to haploid assemblies from HGSVC ^24^ using bwa mem v0.7.17 ^52^. PCR duplicates were marked using samblaster v0.1.26 ^57^, and only primary alignments (mapping quality > 30, read length > 35 bp) were retained. Terminal trimming (5 bp from both ends) was performed using BamUtil v1.0.15 ^58^ to minimize ancient DNA-specific errors.

Variants were called using GATK v3.8 (Genome Analysis Tool Kit 3.8-1-0-gf15c1c3ef) ^59^. Indel realignment was performed with “IndelRealigner”, and genotypes were called with “UnifiedGenotyper”. Filtering followed GATK best practices: (1) variant clusters with “-clusterSize 3” and “-clusterWindowSize 10” options; (2) variants with depth < 1/3x and > 3x (x, overall mean sequencing depth across all variant sites); (3) quality by depth, QD □<□ 2; (4) phred-scaled variant quality score, QUAL□<□30; (5) strand odds ratio, SOR□>□3 (for SNV only); (6) Fisher strand, FS□>□60; (7) mapping quality, MQ□<□40 (for SNV only); (8) mapping quality rank sum test, MQRankSum□< □-12.5 (for SNV only); and (9) read position rank sum test, ReadPosRankSum□<□-8 were filtered.

Callable loci were defined using GATK’s “CallableLoci,” and only insertions within callable regions were analyzed. Divergence was quantified as variant counts per bp in each insertion, and introgressed versus non-introgressed insertions were compared using a likelihood ratio test (R package “lmtest”).

### Retrieval of short-read SNV data from gnomAD

SNV data were obtained from the HGDP and 1KGP call set available in the Genome Aggregation Database (gnomAD v3.1.2) ^60^. We retained only those samples for which corresponding genotype information was present in our filtered PanGenie dataset. To ensure consistency and quality, we restricted analyses to autosomal SNVs that passed all gnomAD quality filters (“PASS”) and exhibited a missing genotype rate < 5% across the retained samples.

### Detection of selection signatures in introgressed structural variants

Signals of selection acting on introgressed SVs were investigated using Ohana ^26^, a maximum likelihood-based framework that models individual genomes as mixtures of *K* ancestry components. This method tests whether the observed AF of each variant is better explained by a selection model relative to a genome-wide null model.

To construct the null model, we first pruned SNVs from the gnomAD dataset using PLINK v1.9 ^61^ with “--indep-pairwise 50 10 0.1” and removed loci with missing genotypes with “--geno 0.0”. We then randomly selected 100,000 SNVs to infer population structure and admixture proportions across present-day humans. The resulting covariance matrix of ancestry components served as the null expectation under neutrality.

For each SV of interest, we compared the likelihood of the observed ancestry component-specific AFs under the neutral and selection models. The strength of evidence for selection was quantified using a likelihood ratio statistic (LRS). Likelihood ratio tests were performed for each SV, and SVs with Bonferroni□corrected p□values < 0.05 were considered significant. We performed these tests iteratively across ancestry components, with *K* varying from 6 to 10 to check consistency (**Figs. S32** and **S33**).

### Annotation of introgressed structural variants

Introgressed SVs were annotated using AnnotSV v3.1.2 ^62^ with ENSEMBL annotation. Repeat elements within introgressed insertions were annotated using RepeatMasker v4.1.2-p1 (http://www.repeatmasker.org; rmblastn engine (“-e nbci”) and “-species human”).

Phenotypic associations for the introgression-informative SNVs were queried using GWAS ATLAS (https://atlas.ctglab.nl/PheWAS), with a focus on pigmentation-related traits. The introgression-informative SNVs were lifted over from GRCh38 to GRCh37 using the UCSC LiftOver tool (https://genome.ucsc.edu/cgi-bin/hgLiftOver). Genome-wide significant associations (*p-value* < 5.0 × 10^-8^) were only considered.

### Breakpoint characterization of the *OCA2 Alu* insertion

Human haplotypes, HG00096 h2 and HG02828 h1 from HGSVC ^24^, and ape genomes ^63^ were aligned to the GRCh38 reference genome using minimap v2.29-r1283 ^64^ with the “-x asm20” option.

To visualize the breakpoints of the *OCA2 Alu* insertion in archaic hominin genomes, we retrieved raw reads from four high-coverage archaic hominin genomes (**Table S5**). Adapter sequences were removed, and overlapping paired-end reads were merged using leeHom v1.2.17 ^65^. For the Chagyrskaya Neanderthal genome (Chagyrskaya 8), we utilized unmapped reads (http://ftp.eva.mpg.de/neandertal/Chagyrskaya/rawBAM/) that had been pre-trimmed and merged. The clean reads were mapped to GRCh38 using bwa mem v0.7.17 ^52^, and breakpoint regions were visualized using IGV v2.12.3 ^66^.

To assess breakpoint consistency across individuals, we examined mapped reads from all present-day human genomes carrying the *Alu* insertion (downloaded from the International Genome Sample Resource; IGSR; https://www.internationalgenome.org/) as well as from the Denisovan genome. Soft-clipped reads supporting the *Alu* insertion were counted to evaluate breakpoint reproducibility across insertion carriers.

### Preparation of data for evolutionary analysis of the *OCA2 Alu* insertion

Hmmix calls of introgressed segments for 1KGP and HGDP samples relative to GRCh38 were downloaded from: https://doi.org/10.5281/zenodo.14136628 ^28^. High-coverage archaic hominin genotypes (Altai Neanderthal, Vindija Neanderthal, Chagyrskaya Neanderthal, and Denisovan) were retrieved from https://zenodo.org/records/13368126 ^28^. Ancestral sequences and recombination maps for chromosome 15 (GRCh38) were obtained from https://doi.org/10.5281/zenodo.11212339 ^28^.

Archaic hominin and present-day human genotypes were merged using bcftools v1.21 ^67^, retaining only polymorphic sites. Genotypes were phased using SHAPEIT v5 ^68^, with common (minor allele frequency (MAF) > 0.001) and rare variants (MAF ≤ 0.001) being phased separately. While higher switch error rates are expected in archaic hominin genomes, phasing accuracy is sufficient for analyses that average across haplotypes ^69,70^. Pairwise variant correlations were computed using PLINK2 ^71^ with the options “--r2-phased ref-based”, “--ld-window 9999”, and “--ld-window-kb 1000”. The visualization of LD structure was generated using LDblockShow ^72^ with “-SeleVar 2”, “-MAF 0.05”, and “-Miss 0.05” parameters.

### Analyses of introgressed segments overlapping the *OCA2* insertion

Introgressed segments were intersected with the genomic position of the *OCA2 Alu* insertion (chr15:27,934,218; GRCh38). Overrepresentation of introgressed segments among insertion carriers was assessed using a *X*^2^-contingency test implemented in scipy v1.15.2 ^73^. For each individual, phased genotypes were used to assign introgressed segments to the haplotype with the fewest mismatches in derived alleles across the *OCA2 Alu* insertion and 24 introgression-informative SNVs identified by hmmix.

### Comparison with empirical and simulated introgressed segments

To contextualize the observed poor matching of putatively Denisovan introgressed segments at the *OCA2 Alu* insertion locus, we compared their properties to those of empirically resampled and simulated introgressed segments. Derived site densities, derived allele ages, and derived allele sharing with archaic references were computed. Empirical distributions were obtained from 10,000 predicted introgressed segments on chromosome 15 with posterior probability > 0.8. Allele ages were extracted from ancestral recombination graphs (ARGs) inferred using RELATE ^74^ (see below).

Simulated distributions were generated using msprime v1.3.3 ^75^. Parameters included: human-archaic hominins split 28,000 generations ago; Denisovans and Neanderthals split 20,000 generations ago; Denisovan branching into two sublineages 12,000 generations ago; Out of Africa (OOA) migration event 2,400 generations ago; and a single Neanderthal and Denisovan introgression event at 1,800 and 1,200 generations ago, respectively, each contributing 2% to the OOA population. Finally, low levels of migration were simulated between OOA and African populations. We assumed a mutation rate of 2.36 × 10^-8^ per bp per generation, and used the HapMap II recombination map for chromosome 15 (GRCh38) ^76^. A single Neanderthal genome was sampled from the introgressing lineage 4,400 generations ago (110 thousand years ago (kya), assuming a generation time of 25 years, corresponding the estimated age of the Altai Neanderthal ^49^), and a Denisovan genome was sampled from the divergent, non-introgressing Denisovan lineage 2,556 generations ago (63.9 kya, assuming a generation time of 25 years, corresponding to the age of the Denisovan individual sequenced to a high coverage ^41^). Additionally, 100 present-day human genomes from both OOA and African populations were sampled. For each of 100 replicates, summary statistics were computed using tskit v0.6.2 ^77,78^ and tspop v0.0.2 ^79^.

### Phylogenetic and ancestral recombination graph analyses of the *OCA2 Alu* insertion

To investigate the evolutionary relationships among haplotypes associated with the *OCA2 Alu* insertion, we first reconstructed a haplotype network using the *OCA2 Alu* insertion and 24 introgression-informative SNVs identified by hmmix. This network was generated using the minimum spanning tree algorithm implemented in PopART ^80^.

To place these haplotypes within a temporal and genealogical framework, we next inferred an ARG for the genomic region surrounding the *OCA2 Alu* insertion using RELATE v1.2.2 ^74^. First, we extracted all biallelic SNVs in the phased VCF using the utilities “ConvertFromVCF” and “PrepareInputFiles.” Second, the ARG was inferred using the “RelateParallel.sh” script under the assumptions of a mutation rate of 1.25 × 10^-8^ per bp per generation, and an effective population size (N_e_) of 30,000. We then extracted the marginal tree corresponding to the SNV most strongly associated with the *OCA2 Alu* insertion (chr15:27,934,733; chr15-27934733-SNV-T-C) using “RelateExtract”. The resulting tree was visualized with a customized version of “TreeViewMutation”.

### Selection tests on the locus of *OCA2 Alu* insertion

To evaluate whether the *OCA2 Alu* insertion locus exhibits evidence of positive selection, we first examined haplotype-based evidence for selection using cross-population extended haplotype homozygosity (XP-EHH) ^81^. XP-EHH was calculated with the Nasioi population as the target population and Central and South Asian populations as a reference. To control for local variation, raw XP-EHH scores were standardized to Z-scores relative to the chromosome-wide background. In addition, given that the *OCA2* region overlaps a known recombination hotspot, XP-EHH scores were further normalized relative to sites within the highest percentile of recombination rates based on the HapMap II recombination map.

To obtain additional evidence for selection and estimate the strength of selection, we used an ABC approach. First, we extended the demographic model for Papuans from Malaspinas et al. 2016 ^82^ to include Nasioi individuals. We assumed the Nasioi population to have split off from the Papuan population 700 generations ago, going through a 90-generation bottleneck of variable size (N_e_ = 600 and N_e_ = 1,200) before expanding back to 8,834, the same size inferred for Papuans. We also added a mutation at the site of the *OCA2 Alu* insertion to the Denisovan population 1,500 generations ago. This mutation was initially neutral, but became beneficial in the Nasioi population 500 generations ago. We varied the selection coefficient between 0.0 and 0.15, performing a total of 15,000 simulations. We simulated a ± 250 kb region around the *OCA2 Alu* insertion, assuming the HapMapII recombination map using stdpopsim v0.1.dev1620+gbae9a69 ^83^ with SLiM v4.0 ^84^ as its engine and sampled the same number of individuals from the African (n = 592), European (n = 640), East Asian (n = 697), Papuan (n = 14), and Nasioi (n = 11) populations as present in 1KGP and HGDP datasets. Using tskit v0.6.4 ^85^ and scikit-allel v1.3.13 ^86^, we then extracted a ± 100 kb region around the *OCA2* insertion and calculated the following summary statistics for the ABC: AF, iHS, genetic diversity (□), and the number of segregating sites, Fay and Wu’s H in the Nasioi population, as well as *F*_ST_ for the Papuan vs. Nasioi, and the East Asian vs. Nasioi populations, respectively. Similarly, we calculated the same summary for the empirical data for a 200 kb genomic region surrounding the *OCA2 Alu* insertion using scikit-allel. Assuming a uniform prior for selection coefficient, we finally estimated the posterior distribution of selection coefficients for the *OCA2* locus from simulations most closely matching the empirical data using the Lenormand sequential sampling scheme ^87^ for ABC implemented in EasyABC ^88^ in R with a predetermined stopping criterion (*p_acc_* = 0.01).

### Genotyping PCR for *OCA2 Alu* insertion in individuals around Bougainville Island

To perform genotyping PCR, primers were designed flanking the *OCA2 Alu* insertion using NCBI primer blast ^89^, approximately 730-930 bp from its breakpoint. Primer sequences can be found in **Table S6**. Q5 Hot Start High-Fidelity 2X Master Mix (New England Biolabs M0494S) was used for PCRs, supplemented with Bovine Serum Albumin (New England Biolabs) to counter melanin inhibition. PCR product was separated by gel electrophoresis on a 1% agarose gel (ThermoFisher Scientific 16500100) supplemented with SYBR Safe DNA Gel Stain (Invitrogen S33102). A representative gel image for genotyping PCR can be found in **Fig. S34**.

Pigmentation measurements and genetic material used in this study were originally collected as part of a larger study investigating the population history and settlement of Island Melanesia ^30,39^. All participants gave their informed consent to participate as research subjects. Institutional review board (IRB) approval for the collection and analysis of data was obtained from Temple University (IRB 99-226), Pennsylvania State University (IRB 00M558-2), and the Papua New Guinea Medical Research Advisory Committee for this collaborative project with the PNG Institute of Medical Research.

In total, 458 individuals from the Bismarck Archipelago and Bougainville in the Solomon Islands were genotyped for the *OCA2 Alu* insertion using primers described above. 32 of them were excluded from downstream analyses due to unreliable genotyping results or because their demographic information was unavailable. Skin and hair pigmentation were measured using a Dermaspectrometer (Cortex Technologies, Hadsund, Denmark). Pigmentation measurements were reported as melanin index, as described in Norton et al. 2016 ^39^. Additional demographic information was obtained for each individual, including sex, age, island affiliation, linguistic grouping (Austronesian or Papuan), and neighborhood. Briefly, “neighborhood” designations were an indication that an individual and both parents were from the same village and spoke the same language.

### CRISPR/Cas9 editing of human iPSC

We introduced the *Alu* element of *OCA2* into KOLF2.1J iPSCs using previously published methods ^32,90^. Briefly, Cas9 sgRNAs flanking the *Alu* insertion site were chemically synthesized with 2′-O-methyl and 3′-phosphorothioate end modifications (Synthego CRISPRevolution sgRNA) (**Table S6**) and resuspended in TE buffer at a concentration of 4 μg/μL. RNP was formed by combining SpCas9 nuclease (HiFi V3, IDT) with a sgRNA at a molar ratio of 1:4. To encourage insertion of the *Alu* element via homology-directed repair (HDR), we generated a template plasmid containing the *OCA2 Alu* element flanked with approximately 500 bps upstream and downstream of the insertion site (pUC57, Genscript) and resuspended in DPBS at a concentration of 200 pmol/μL (see **Table S6** for the insertion sequence). Cells were transfected using Amaxa nucleofection with P3 Primary Cell 4D-Nucleofector 16-well Strips (Lonza). Each well contained a single-cell suspension of 1.6 × 10^5^ cells in 20 μL of Primary Cell P3 buffer with supplement (Lonza) containing 2 μg Cas9, 1.6 μg sgRNAs, and 2 μg template plasmid. Nucleofection was performed using the Amaxa program CA-137. Immediately following electroporation, cells were distributed to a Synthemax-coated 6-well plate containing StemFlex, CloneR2, and 1 μM final Alt-R HDR enhancer (IDT). Cells were incubated at 32°C for 2 days before transfer to 37°C. At 24 h post-nucleofection, and every other day thereafter, the media were replaced with only StemFlex. Upon reaching near confluency, cells were single-cell plated into a vitronectin-coated 10-cm dish. At day 10, 48 colonies were manually picked and expanded to a vitronectin-coated 96-well plate for 4-5 days before being frozen down. A duplicate set of clones was cultured and lysed as described previously ^90^, after which cells were amplified by target site PCR and screened for the presence of the *Alu* element (**Fig. S35**). Selected clones were further validated by Sanger sequencing of the target site amplicon. We set up 2 independent RNPs with 2 individual sgRNAs, respectively, and isolated clones from both sets, of which only one yielded the edited clone. Clone H08 was identified as being heterozygous for the *Alu* insertion, with one allele containing the intended insertion and the other allele lacking the insertion but containing an unintended 11 bp deletion at the CRISPR site due to non-homologous end-joining. This was confirmed by amplicon NGS of clone H08.

### Differentiation of human iPSCs into melanocytes

We used the human iPSC line KOLF2.1J ^32^ (The Jackson Laboratory) for all differentiation experiments. Our protocol was adapted from Saidani et al. 2023 ^35^, with two key modifications: (1) differentiation plates were coated with fibronectin (10 µg/mL) to enhance embryoid body (EB) adhesion, and (2) EBs were plated on day 2 instead of day 7.

Briefly, EBs were generated by detaching iPSCs using Accutase for 5-7 minutes at 37□°C. After removing Accutase, cells were triturated in fresh mTeSR+ medium (STEMCELL Technologies) supplemented with 1× RevitaCell (Gibco, A26445-01). The single-cell suspension was counted using a hemocytometer, and 30,000 cells were seeded on day 0 media into ultra-low attachment 96-well V-bottom plates. Aggregation was achieved by centrifugation at 200 × g for 3 minutes.

Day 0 media consists of a 1:1 mixture of Neurobasal (Gibco, 21103049) and Ham’s F12 (Gibco, 1176505421103049) and is supplemented with CHIR99021 (3 µM) (Millipore), LDN-193189 (0.2 µM) (STEMCELL Technologies, 72147), SB431542 (40 µM), 2% B-27 without vitamin A (ThermoFisher, 12587010), 1% N-2 (ThermoFisher, 17502048), and 1x Revitacell (Gibco, A2644501) for 2 days at 37□°C.

On day 2, 10-12 EBs were transferred to fibronectin-coated 6-well plates and cultured in day 0 media lacking Revitacell. On day 4, attached EBs were fed day 4-6 media, a 1:1 mixture of Neurobasal/Ham’s F12, supplemented with CHIR99021 (3 µM), EDN3 (100 nM) (American Peptide or ENZO Life Sciences), SCF (50 ng/mL), human recombinant BMP4 (40 ng/mL), and ascorbic acid (0.3 mM). Cells were fed again with day 4-6 media on day 6.

From day 7 to day 30, EBs were cultured in MGM-4 medium (Lonza, CC-3249) supplemented with CHIR99021 (0.5 µM), EDN3 (100 nM), SCF (50 ng/mL), BMP4 (0.02 nM), and ascorbic acid (0.3 mM). Medium was refreshed every 2-3 days.

After 28-32 days, cells were dissociated using 0.05% trypsin and replated onto fibronectin-coated wells. These passaged melanocytes were either maintained in MGM-4 medium supplemented with 20 µM forskolin (Sigma-Aldrich, F3917-10MG), harvested for RNA extraction, or processed for immunocytochemistry (ICC) and flow cytometry (FC).

### Microscopy, immunocytochemistry, and flow cytometry analyses of iPSC-derived melanocytes

Morphological changes during the differentiation of iPSCs into melanocytes were monitored over time using either a Leica MD i1 or an EVOS® FL Inverted Fluorescence Microscope (Invitrogen™ AMF4300) (**Fig. S36**). Successful differentiation was confirmed by ICC analyses and FC (**Figs. S37** and **S38**).

ICC was done as described previously with some modifications ^91,92^. Cells were fixed using 4% PFA in 1×PBS for 20 min, followed by a 1×PBS wash for 10 min. Permeabilization was done using 0.4% Triton-X 100 (Sigma, T8787-250ml), followed by a 1 × PBS for 10 min wash. Cells were blocked using 4% BSA (New England Biolabs, B9000S) for 1.2 hours. All primary antibody incubations were overnight at 4°C. Secondary antibody incubations were overnight at 4°C. The primary antibodies and dilutions used were: TYRP1-488 (1:100; Biotechne, NBP2-54446AF488) and OCA2 (1:100; Thermofisher Scientific, PA5-102825). Secondary antibodies were Cyanine and Alexa dye-conjugated AffiniPure F(ab″)2 fragments purchased from Jackson ImmunoResearch Laboratories and used at a dilution of 1:250. DAPI (1 µg/ml: Life Technologies, D1306) was used for nuclei staining together with secondary antibodies.

For FC, cells were first dissociated and stained with LIVE/DEAD™ Fixable Aqua Dead Cell Stain in 1 × PBS for 20 minutes at room temperature, protected from light. After staining, cells were washed with 1 × PBS and centrifuged at 300 × g for 5 minutes. The resulting cell pellet was processed using the BD Cytofix/Cytoperm™ Fixation/Permeabilization Kit according to the manufacturer’s instructions. Following fixation and permeabilization, cells were incubated with either TYRP1-Alexa Fluor 488 or the corresponding Alexa Fluor 488-conjugated isotype control, diluted in FACS buffer (1 × PBS + 2% fetal bovine serum), for 30 minutes on ice in the dark. After incubation, cells were washed, centrifuged as described above, and resuspended in FACS buffer for analysis on a BD LSR Fortessa flow cytometer.

### Expression qPCR for *OCA2* and melanocyte markers

Two primer sets were designed to capture different portions of the *OCA2* gene exons. Previously published primer sets were used for assessing *Sox10*, *MiTF*, and *TYRP1* gene expressions ^35^ (**Table S6**). Melanocytes either had mRNA extracted fresh or were stored in RNAlater following standard protocols (ThermoFisher Scientific AM7021). mRNA was extracted using RNeasy Mini Kit (Qiagen #74104) for bulk samples or using Single Cell Lysis Kit (ThermoFisher Scientific 4458235) for small cell numbers. The mRNA was immediately reverse-transcribed into cDNA using QuantiTect Reverse Transcription Kit (QIAGEN 205311). To assess gene expression, Powerup Sybr Green Master Mix (ThermoFisher Scientific A25777) was mixed with cDNA diluted 1:10 and expression primers (0.5uM). Sample fluorescence during qPCR was measured using an Applied Biosystems QuantStudio 5 Dx Real-Time PCR System (ThermoFisher Scientific). Differential *OCA2* expression was calculated with the 2(-Delta Delta C(T)) method using *GAPDH* or β*-ACTIN* as housekeeping genes.

### RNA sequencing and analyses

RNA-Seq libraries were prepared using the RNA Library Prep with Polaris Depletion kit (Watchmaker) and Pure Beads (Roche). Sample inputs were normalized to 100 ng of RNA, and 1 μL of 1:500 ERCC 2 (Invitrogen) was spiked into samples. Samples were ribosomal RNA and globin-depleted and then bead-purified. Purified RNA was fragmented at 80°C for 1 min, targeting a fragment range of 300-350 bp. Fragmented RNA was reverse-transcribed, followed by second-strand synthesis and A-tailing. A-tailed, double-stranded cDNA fragments were ligated with xGen UDI-UMI (IDT) adaptors. Adaptor-ligated DNA was bead-purified. This was followed by 12 cycles of PCR amplification. The final libraries were bead-purified and sequenced on the Illumina Novaseq X platform (151 bp paired-end reads, ∼450 million reads per sample).

Initial quality control was performed using FastQC v0.11.9 (https://www.bioinformatics.babraham.ac.uk/projects/fastqc/). The paired-end reads were aligned to the human reference genome, GRCh38, with STAR v2.7.11b ^93^ and quantified using featureCounts v2.1.1 ^94^ with GENCODE v49 annotation ^95^. For the *OCA2* gene, normalized counts and dispersion parameters were estimated using DESeq2 v1.48.2 ^96^. HET and WT differences at each time point were evaluated using a parametric bootstrap test. For the *OCA2*, we estimated size-factor-normalized means and negative binomial dispersion parameters using DESeq2. Under a null model assuming no group effect, we generated 10,000 bootstrap datasets by simulating counts from a negative binomial distribution with these estimated parameters. For each bootstrap replicate, we computed the log□(HET) − log□(WT) difference, and empirical one□sided p□values were obtained as the proportion of simulated statistics greater than or equal to the observed value. Differential exon usage was quantified using DEXSeq v1.54.1 ^97^ with HT-seq ^98^ counts corresponding to non-overlapping exon regions prepared using the script “dexseq_prepare_annotation.py” from the package. Samples from days 16 and 19 were considered as one group for this analysis.

### Processing ChIP-seq data

H3K27ac ChIP-seq data (accessions: PRJNA377564, PRJNA723190, and PRJNA1138230) was reanalyzed with the ENCODE chip-seq-pipeline2 v1.3.5.1. Reads were trimmed using Trimmomatic v0.39 ^99^, aligned using Bowtie v2.2.6 ^100^, and deduplicated using Picard v1.126 (https://broadinstitute.github.io/picard/). Peaks were called using MACS v2 ^101^, and results were visualized using trackplot (https://github.com/PoisonAlien/trackplot).

### Chromatin immunoprecipitation (ChIP) and ChIP-qPCR

Day 19 and 30 hiPSCs differentiated to the melanocyte lineage (3-4 million cells) were fixed with 2% formaldehyde (Sigma-Millipore F1635-25ML) for 20 minutes at RT and quenched with 125 mM glycine (Sigma-Millipore 50046-250G). The cells were pelleted at 300 x G for 5 minutes at 4°C, the supernatant was removed, and rinsed with ice-cold phosphate-buffered saline (PBS). Cells were then snap frozen until use. ChIP was performed as previously described ^102,103^. Cells were resuspended in cell lysis buffer (10 mM Tris HCl [pH 7.5], 10 mM NaCl, 0.5% NP-40, and protease cocktail inhibitor (PCI) (Pierce, A32965)), and incubated on ice for 15 minutes. After dounce homogenization on ice, the nuclei were pelleted at 300 x G for 5 minutes at 4°C and incubated in MNase digestion buffer (20 mM Tris HCl [pH 7.5], 15 mM NaCl, 60 mM KCl, 1 mM CaCl_2,_ and PCI supplemented with 1,000 gel units of micrococcal nuclease per 5 million cells (NEB, M0247S). The nuclei were incubated for 20 min at 37°C. The reaction was stopped by adding an equal volume of sonication buffer (100 mM Tris HCl [pH 8.1], 20 mM EDTA, 200 mM NaCl, 0.2% sodium deoxycholate, 2% Triton X-100, and PCI). Samples were sonicated using Diagenode Bioruptor (Diagenode), 10 cycles of 30”-on, 30”-off at 4°C, and centrifuged at 21,000xg for 5 min. Histone ChIP was performed with ∼1 ug of DNA and 2 μg of anti-Histone H3 (acetyl K27) (Abcam, ab4729), anti-Histone H3 (mono methyl K4) (Abcam, ab8895), and normal Rabbit IgG (Cell Signaling Technology, 2729) antibodies overnight at 4°C. This was followed by enrichment with protein G-Dynabeads (ThermoFisher Scientific, 10004D) at 4°C for 3 hours. Sequential washes were performed with ChIP buffer (50 mM Tris [pH 8.1], 10 mM EDTA, 100 mM NaCl, 1% Triton X-100, and 0.1% sodium deoxycholate), ChIP buffer with 0.5 M NaCl, Tris/LiCl buffer (10 mM Tris [pH 8.1], 1 mM EDTA, 0.25 M LiCl_2_, 0.5% NP-40, and 0.5% sodium deoxycholate) and Tris/EDTA buffer (50 mM Tris [pH 8.0] and 10 mM EDTA). Immune complexes were eluted in 100 ul of elution buffer (10 mM Tris HCl [pH 8.0], 10 mM EDTA, 150 mM NaCl, 1% SDS, 5 mM DTT), incubated for 30 minutes at 65°C. Eluates were reverse cross-linked overnight at 65°C with 5 ul of Proteinase K (20 mg/ml, Invitrogen, AM2548). DNA was purified using a ChIP DNA Clean & Concentrator kit (Zymo Research, D4014). Enrichment was measured by qPCR relative to 1% of input DNA using iTaq Universal SYBR Green Supermix (Biorad, 1725121). Primer sequences are listed in **Table S6**.

## Data availability

The raw sequence reads from archaic hominins, HGDP, and great apes were retrieved from Sequence Reads Archive (SRA) with the following accessions: PRJEB1265 (Altai Neanderthal), PRJEB21157 (Vindija Neanderthal), PRJEB3092 (Denisovan), PRJEB6463 (HGDP), and PRJNA189439 (great apes). The sequence reads from the Chagyrskaya Neanderthal were retrieved from http://ftp.eva.mpg.de/neandertal/Chagyrskaya/rawBAM/. For the genotypes of 1KGP, we downloaded publicly available PanGenie calls (https://ftp.1000genomes.ebi.ac.uk/vol1/ftp/data_collections/HGSVC2/release/v2.0/PanGenie_results/20201217_pangenie_merged_bi_all.vcf.gz). For the short-read SNV data, we retrieved the HGDP and 1KGP callset from gnomAD v3.1.2 (https://gnomad.broadinstitute.org/downloads#v3-hgdp-1kg). The raw sequence reads from Southeast Asian and Oceanian populations were retrieved from SRA and European Genome-phenome Archive (EGA) with the following accessions: PRJEB9586, EGAD00001001634, EGAD00001004156, EGAD00001007783, and EGAD00001006880. The great apes’ genome sequences were retrieved from https://github.com/marbl/Primates. The ChIP-seq data for melanoma and melanocyte were retrieved from SRA with the following accessions: PRJNA377564, PRJNA723190, and PRJNA1138230.

## Code availability

No custom codes were used for this study.

## DECLARATIONS

## Acknowledgments

We thank Omer Gokcumen, Emilia Huerta-Sanchez, and Nolan Kamitaki for helpful discussions, suggestions, and feedback on the manuscript.

## Funding

K.K. was supported in part by a Jackson Laboratory Postdoctoral Scholar Award.

## Competing interests

The authors declare that they have no competing interests.

## Authors’ contributions

K.K. and C.L. conceived the study. K.K., A.P., J.M.A., and C.L. drafted the manuscript. K.K., P.H., and F.Y. performed genotyping analyses. K.K., A.P., and J.M.A. performed evolutionary analyses. K.K. performed all other genomic analyses. S.A.S. and H.L.N. performed genotyping PCR. Q.P. and J.A.M. performed nucleofection of KOLF2.1J. N.M., S.A.S., and N.A.J.O. performed melanocyte differentiation. N.M. performed expression qPCR and ICC. S.A.S. performed ChIP-seq analysis and ChIP-qPCR. C.L. managed the project. All authors critically revised and approved the final manuscript.

## Notes

### Competing Interest Statement

The authors have declared no competing interest.

