## Supplementary figures for "A Denisovan-derived *Alu* insertion in *OCA2* contributes to pigmentation diversity in present-day Melanesians"

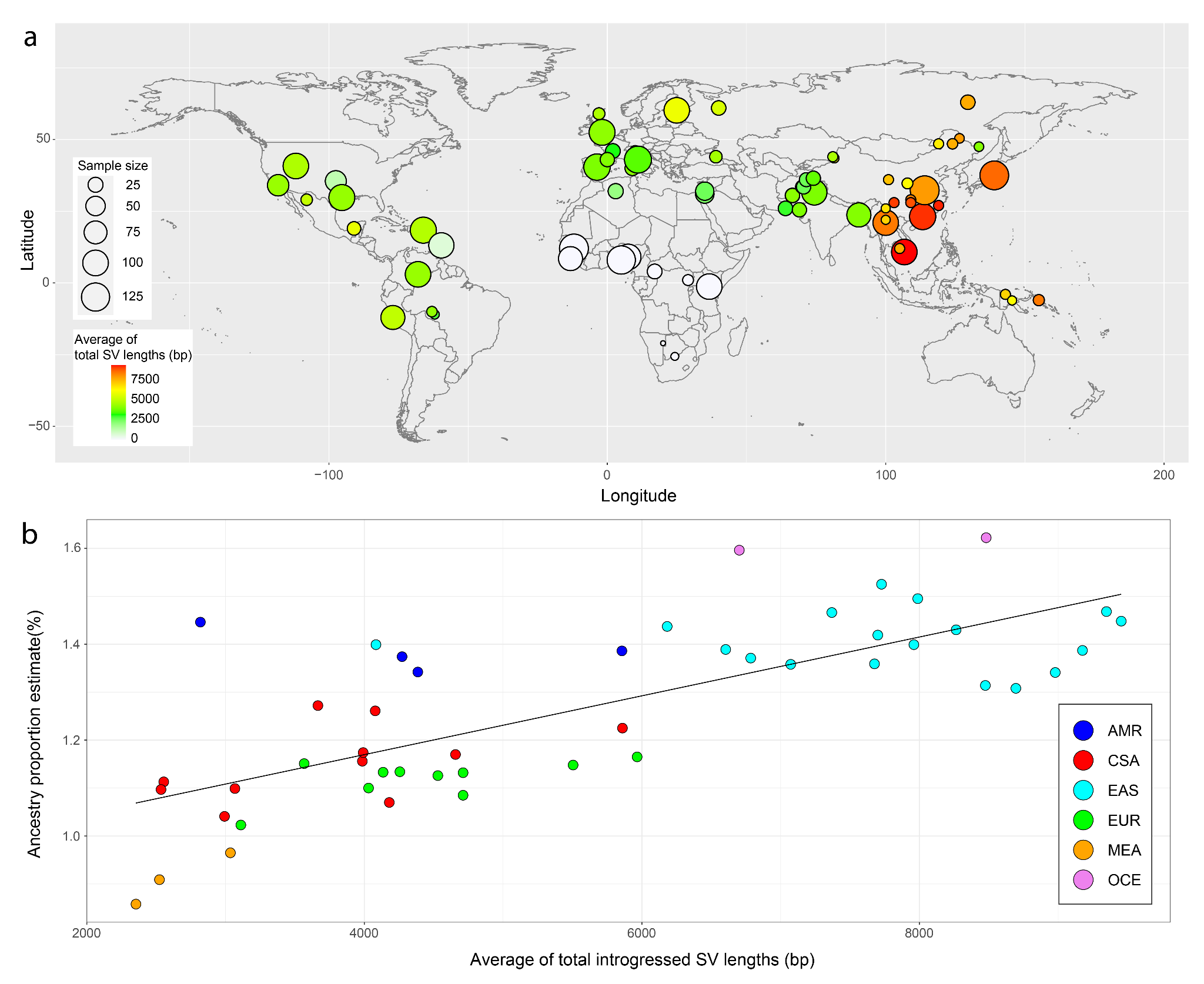

**Supplementary Figure 1. Global distribution of total length of introgressed structural variants (SVs) per individual.** (a) Global distribution of total SV length introgressed from Neanderthal in 3,332 present-day human genomes from the 1000 Genomes Project and the Human Genome Diversity Project. The size and color of the circles represent the sample size and the average of introgressed SV length per population, respectively. (b) Correlation between the average and the Neanderthal ancestry proportion from Sankararaman et al. 2016 ^1^. Each dot represents a population with fill colors indicating its superpopulation designation (AMR: America, CSA: Central and South Asia, EAS: East Asia, EUR: Europe, MEA: Middle East, and OCE: Oceania). Only populations present in both our data and Sankararaman et al. 2016 (n = 48) were used for this comparison.

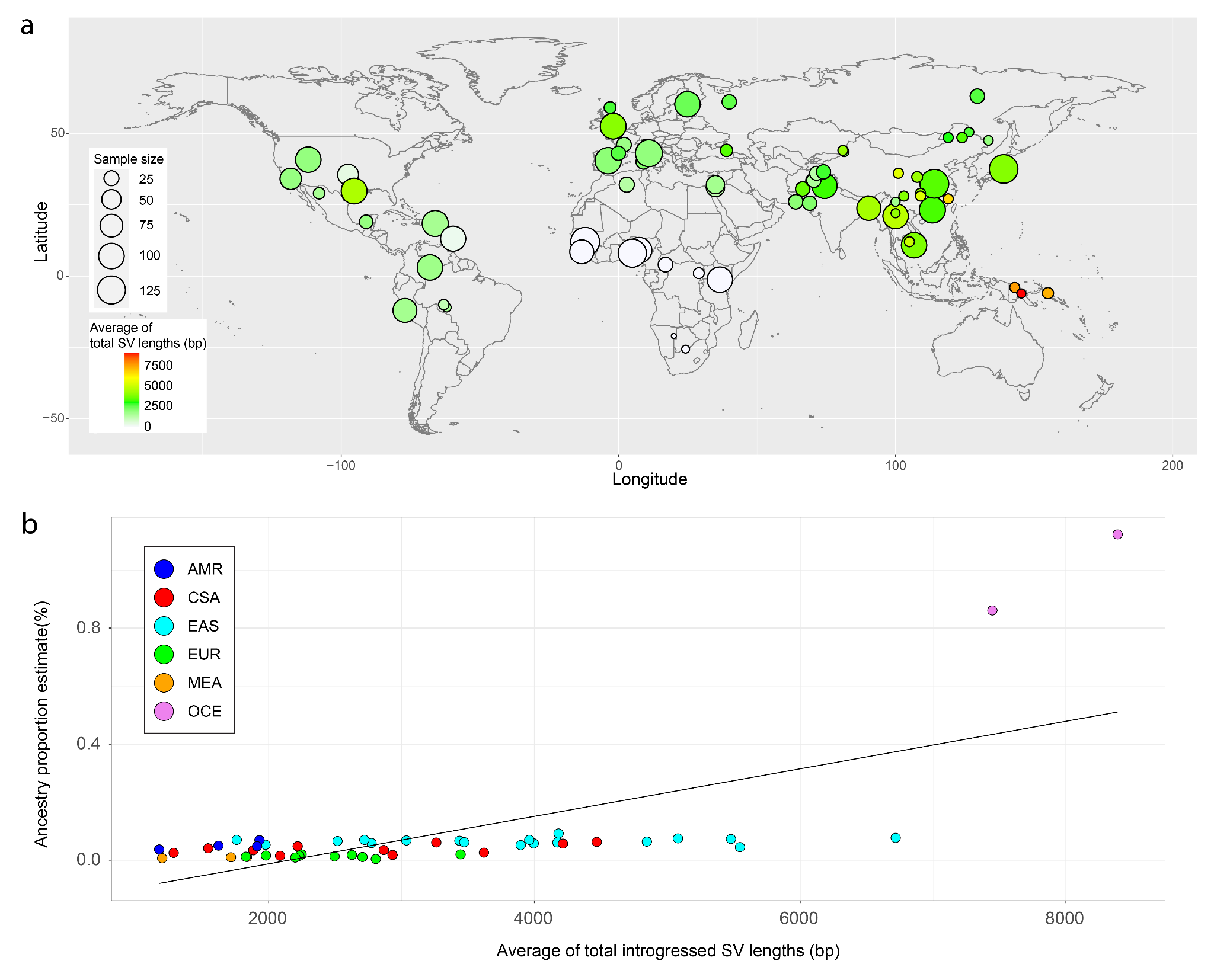

**Supplementary Figure 2. Global distribution of total length of introgressed structural variants (SVs) per individual.** (a) Global distribution of total SV length introgressed from Denisovan in 3,332 present-day human genomes from the 1000 Genomes Project and the Human Genome Diversity Project. The size and color of the circles represent the sample size and the average of introgressed SV length per population, respectively. (b) Correlation between the average and the Neanderthal ancestry proportion from Sankararaman et al. 2016 ^1^. Each dot represents a population with fill colors indicating its superpopulation designation (AMR: America, CSA: Central and South Asia, EAS: East Asia, EUR: Europe, MEA: Middle East, and OCE: Oceania). Only populations present in both our data and Sankararaman et al. 2016 (n = 48) were used for this comparison.

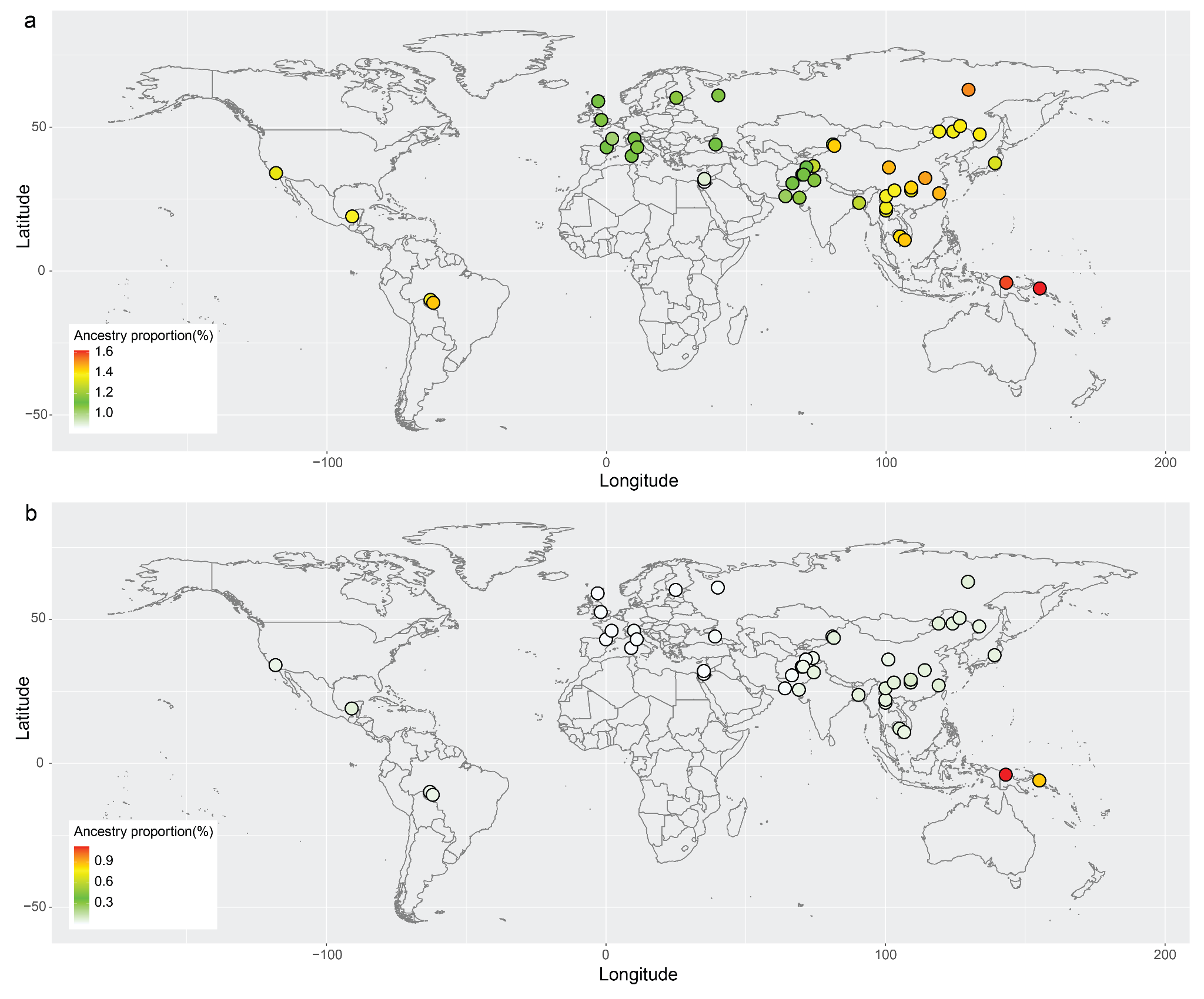

**Supplementary Figure 3. Global distribution of archaic hominin ancestry proportions based on single-nucleotide variants (SNVs).** (a) Global distribution of Neanderthal ancestry proportion. (b) Global distribution of Denisovan ancestry proportion. The ancestry proportions were obtained from Sankararaman et al. 2016 ^1^. Only populations present in both our data and Sankararaman et al. 2016 (n = 48) were used for this visualization.

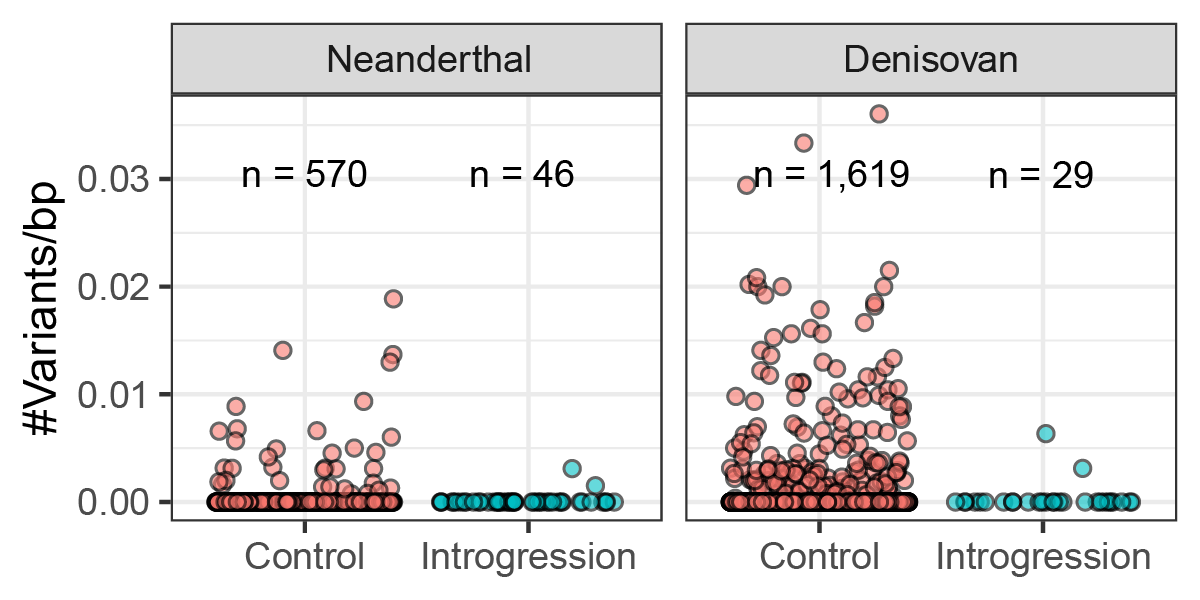

**Supplementary Figure 4. The number of variants per base pair within introgressed insertions (“Introgression”) compared to that of non-introgressed insertions (“Control”).**

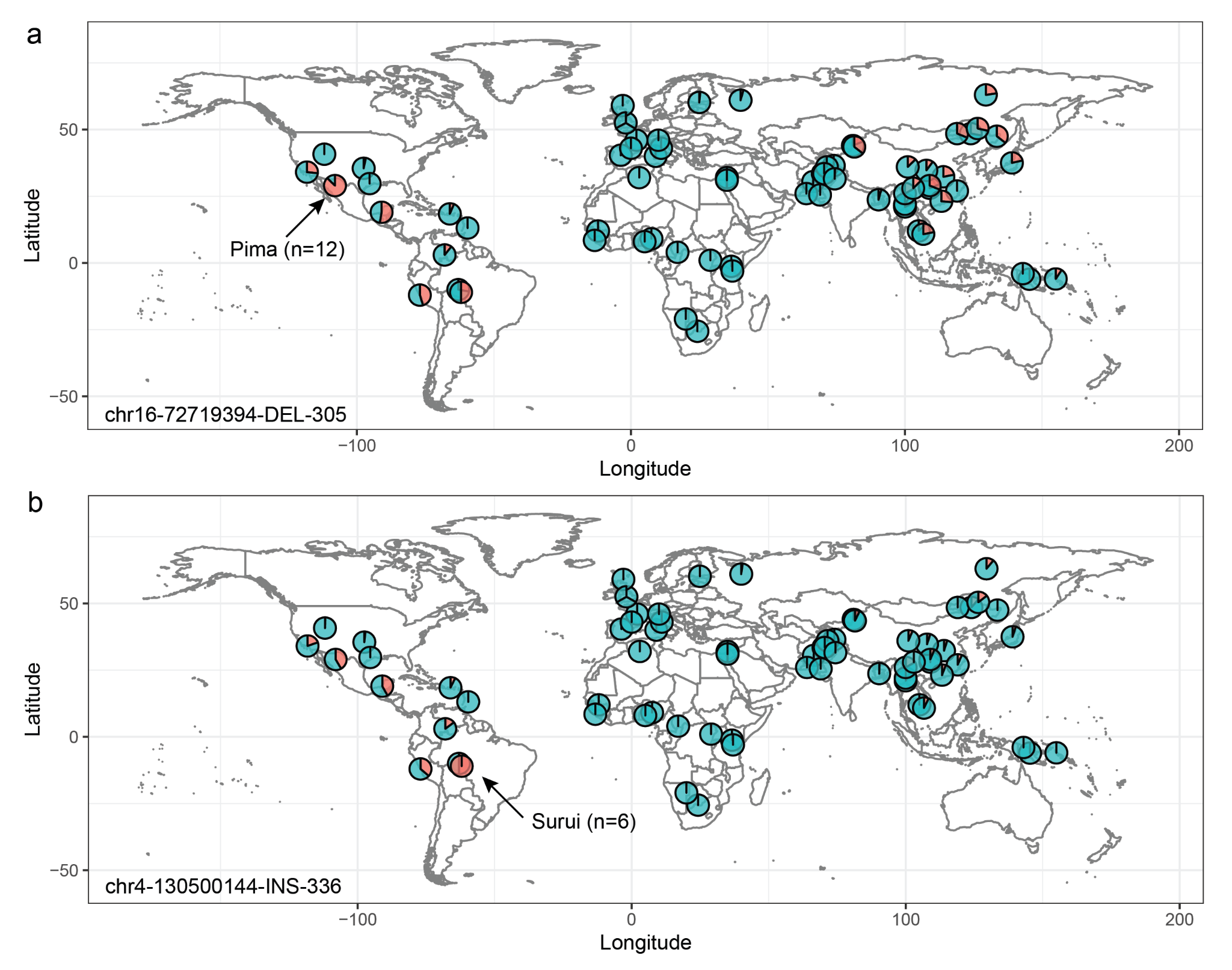

**Supplementary Figure 5.** **Examples of global allele frequency distribution of introgressed structural variations.** (a) A Neanderthal-derived 305 bp deletion on chromosome 16 (chr16-72719394-DEL-305) and (b) A Denisovan-derived 336 bp insertion on chromosome 4 (chr4-130500144-INS-336).

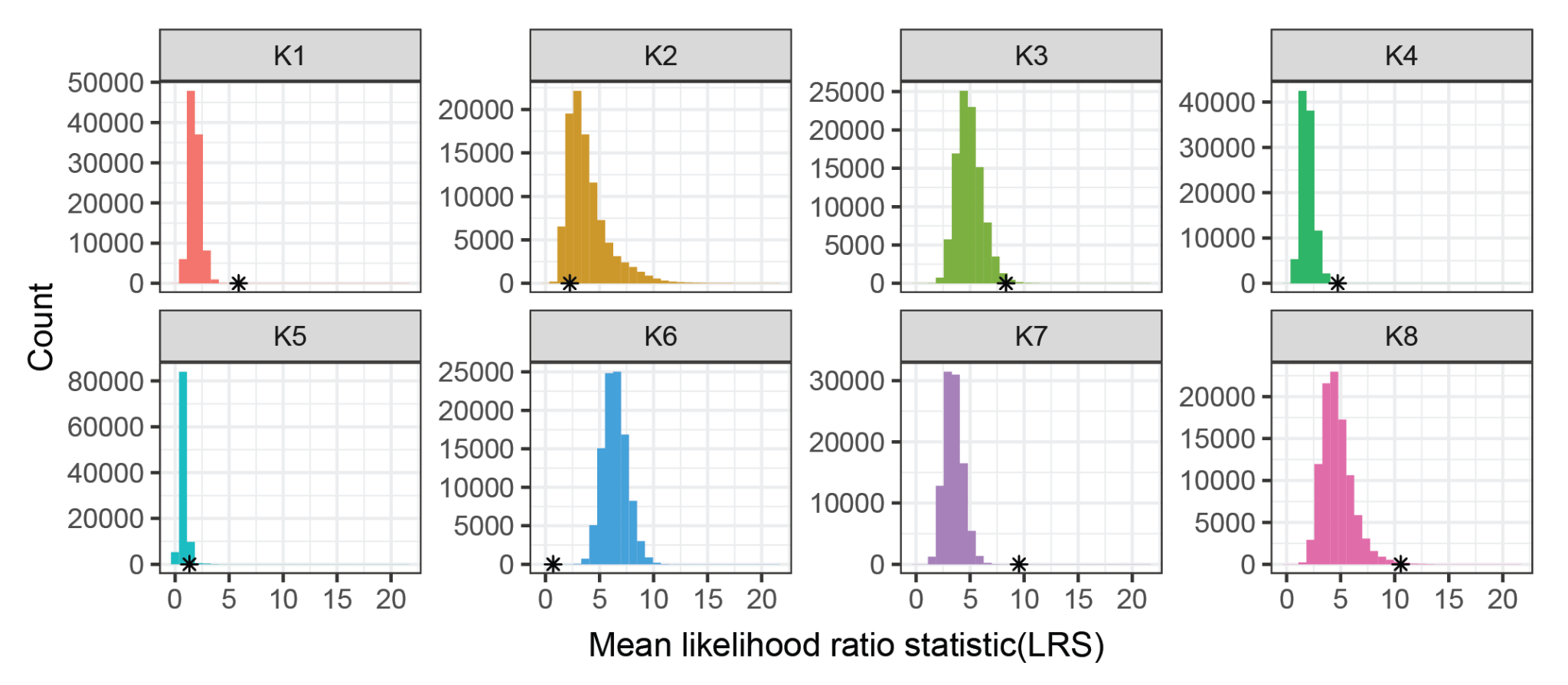

**Supplementary Figure 6.** Selection signatures of the introgressed SVs for each ancestry component under the *K* = 8 model. The asterisk (*) indicates the mean likelihood ratio statistic of the introgressed SVs.

Supplementary Figure 7

**Supplementary Figure 7.** **Individual ancestry proportions estimated by Ohana with *K*=8**. AFR: Africa, AMR: America, CSA: Central and South Asia, EAS: East Asia, EUR: Europe, MEA: Middle East, and OCE: Oceania.

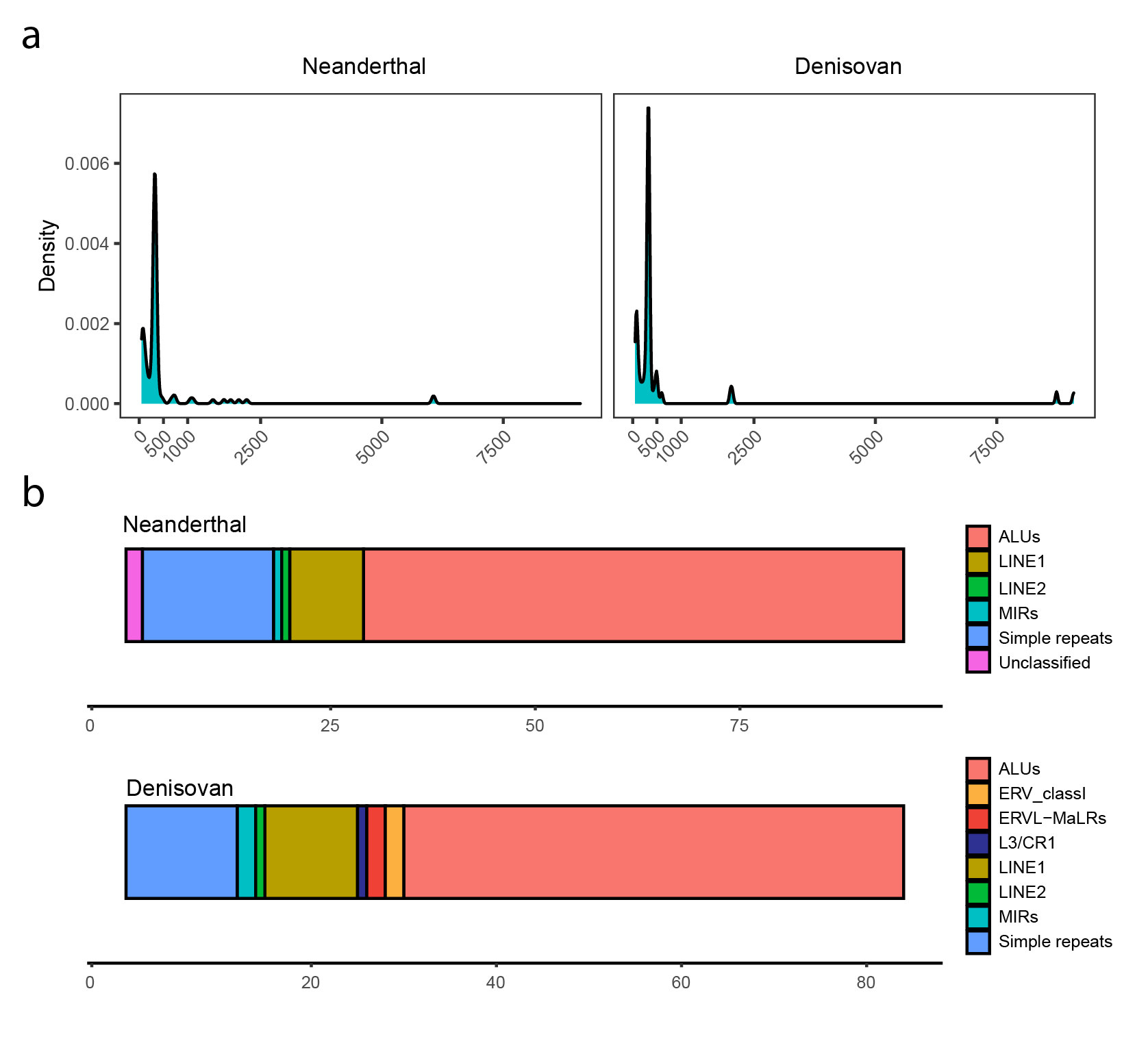

**Supplementary Figure 8. Repeat element annotation of the introgressed insertions.** (a) Size distribution of the introgressed insertions. (b) The number of repeat elements annotated in the introgressed insertions.

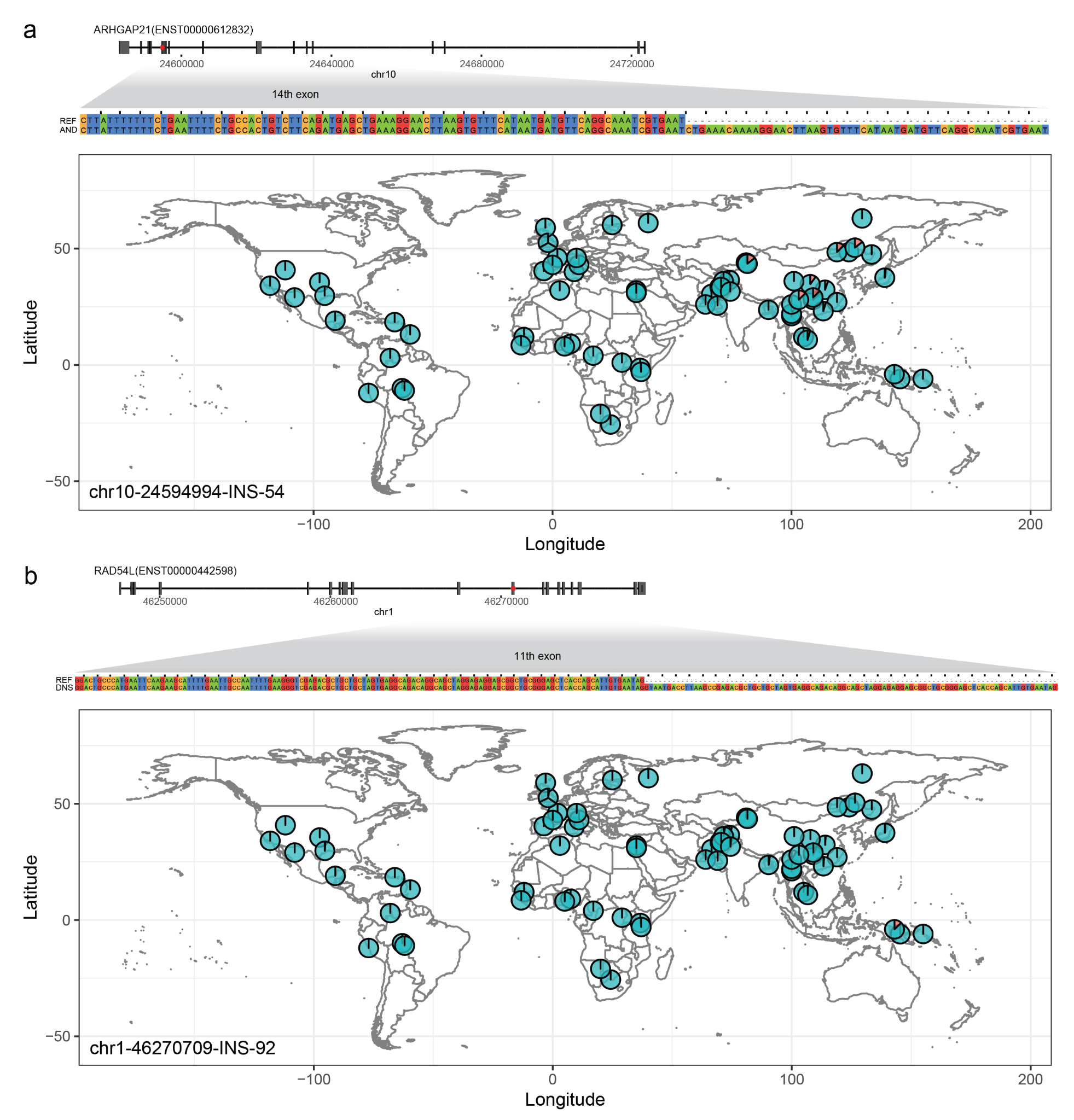

**Supplementary Figure 9.** **Global allele frequency distribution of two frameshift insertions introgressed from (a) Neanderthal (chr10-24594994-INS-54) and (b) Denisovan (chr1-46270709-INS-92).** The sequence of insertion is represented at the top of the allele frequency distribution, compared to the GRCh38 reference sequence. The red asterisk indicates the position of the variant in the context of gene structure. REF: reference, AND: Neanderthal, and DNS: Denisovan.

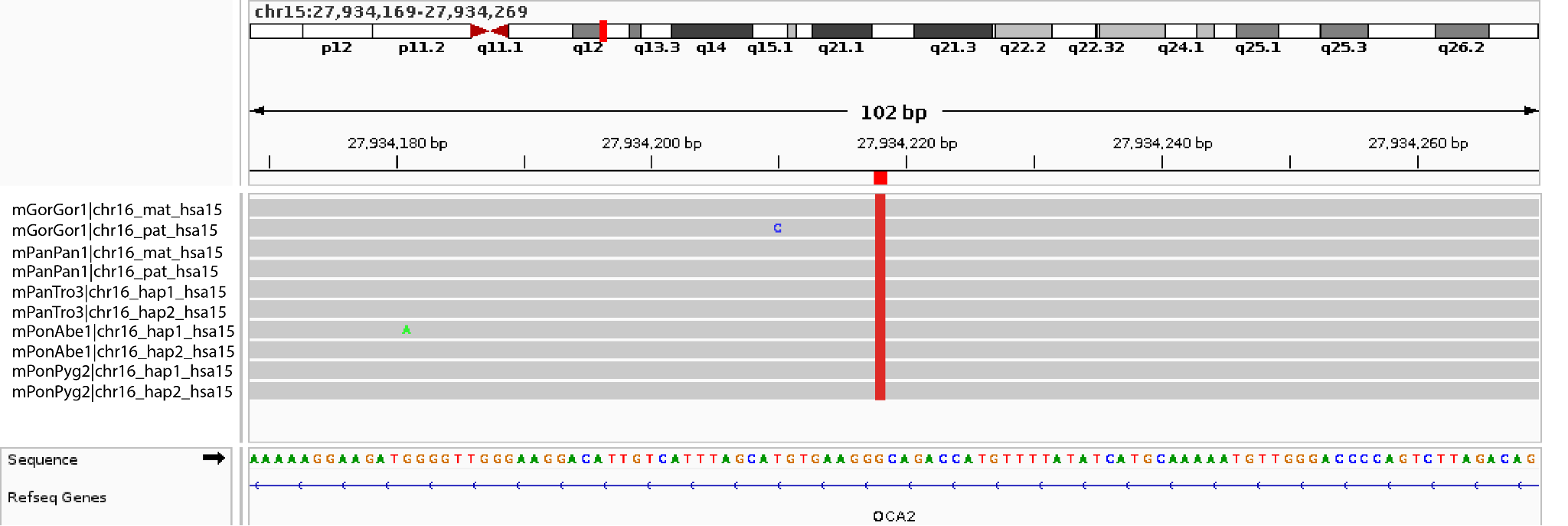

**Supplementary Figure 10. Alignments of great ape genomes around the breakpoint of the Denisovan-derived *Alu* insertion in *OCA2*.** Alignments of great ape genomes from Yoo et al. 2025 ^2^ were visualized for the 100 bp region spanning the breakpoint (chr15:27,934,169-27,934,269). The vertical red line indicates the *Alu* insertion breakpoint based on the GRCh38 reference genome (chr15:27,934,218). mGorGor1: Gorilla, mPanPan1: Bonobo, mPanTro3: Chimpanzee, mPonAbe1: Sumatran orangutan, and mPonPyg2: Bornean orangutan.

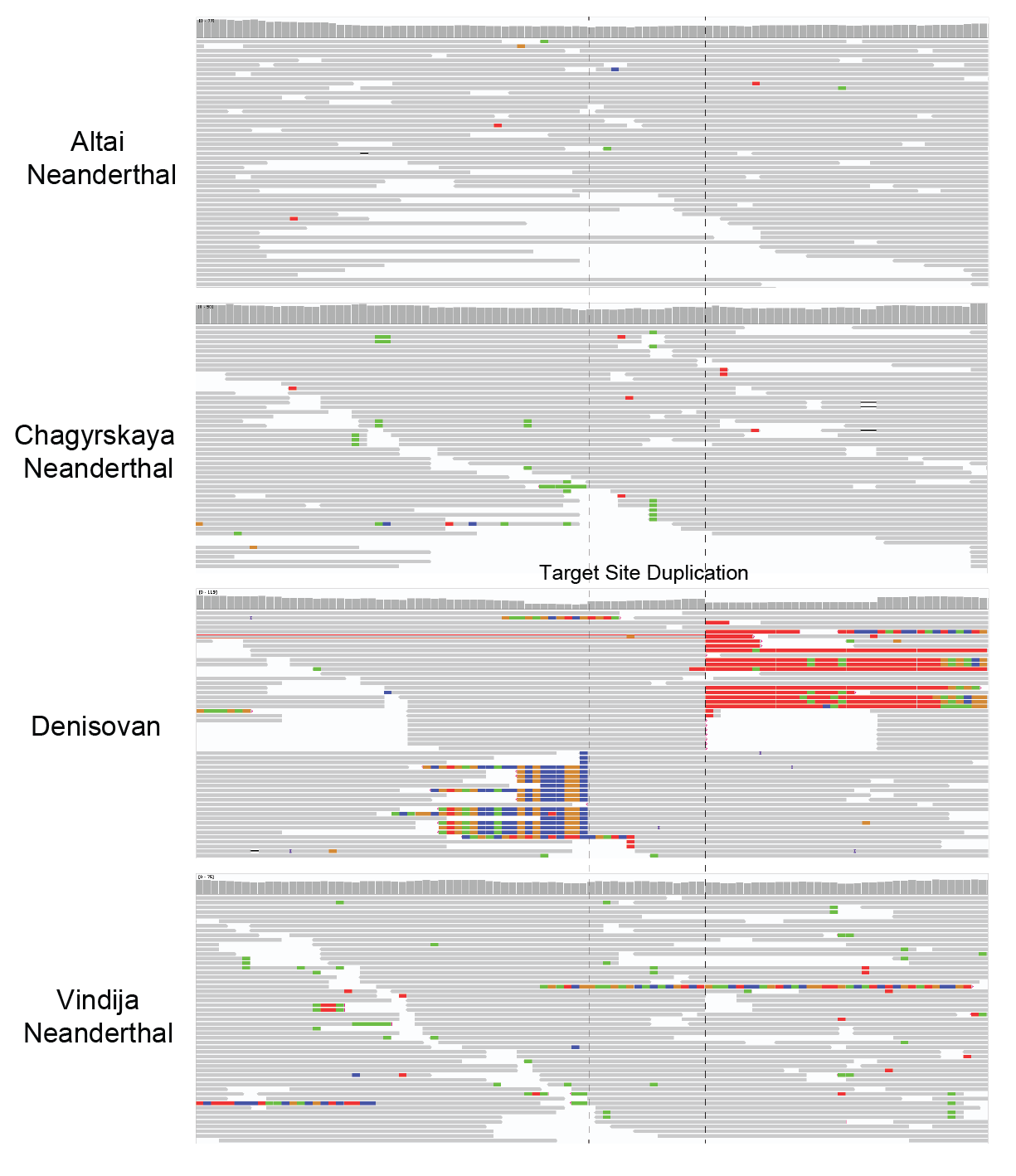

**Supplementary Figure 11. Reads pileups of archaic hominin genomes around the breakpoint of Denisovan-derived *Alu* insertion in *OCA2*.** Reads pileups of the high-coverage archaic hominin genomes were visualized for the 100 bp region spanning the breakpoint based on the GRCh38 reference genome (chr15:27,934,169-27,934,269). The bar chart at the top of each reads pileup represents per-base coverage depth.

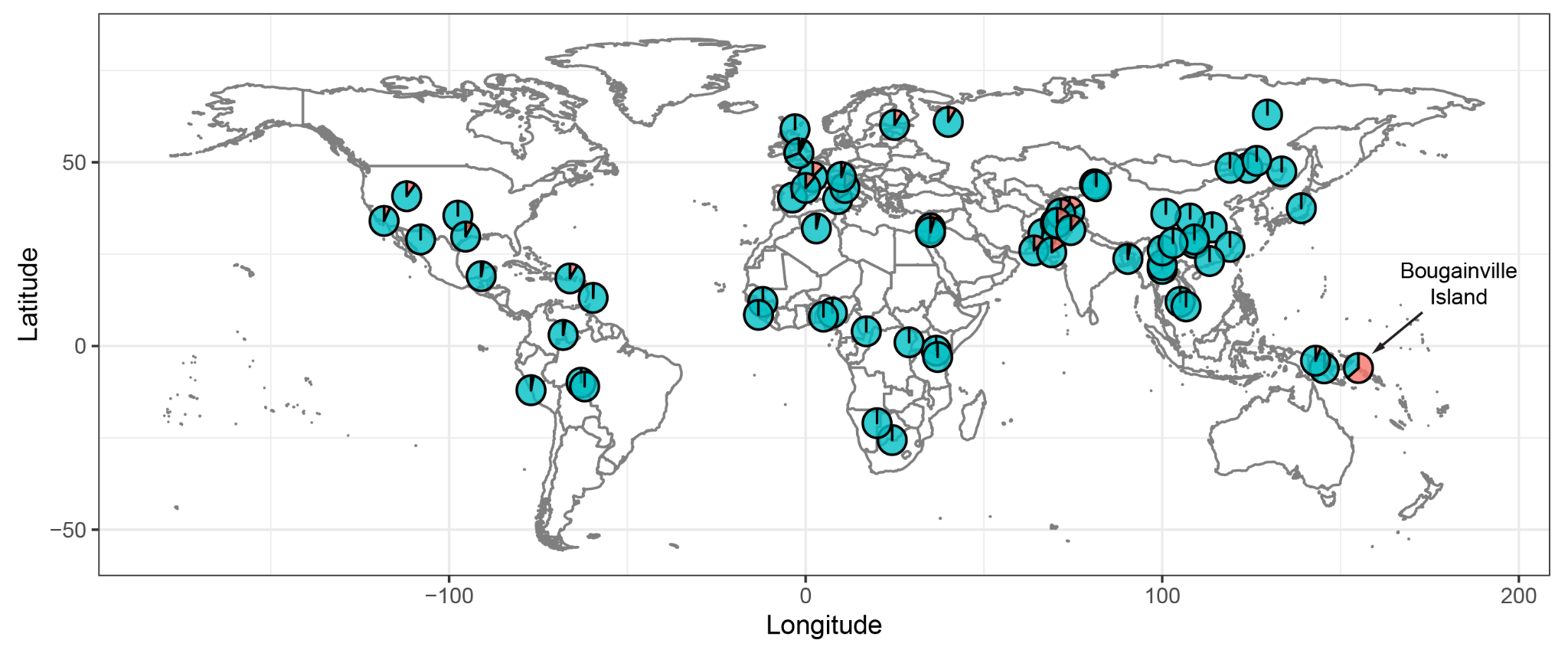

**Supplementary Figure 12. Global distribution of Denisovan-derived *Alu* insertion in *OCA2*, based on the 1000 Genomes Project and the Human Genome Diversity Project datasets.** The red color indicates the allele frequency of the *Alu* insertion in each population.

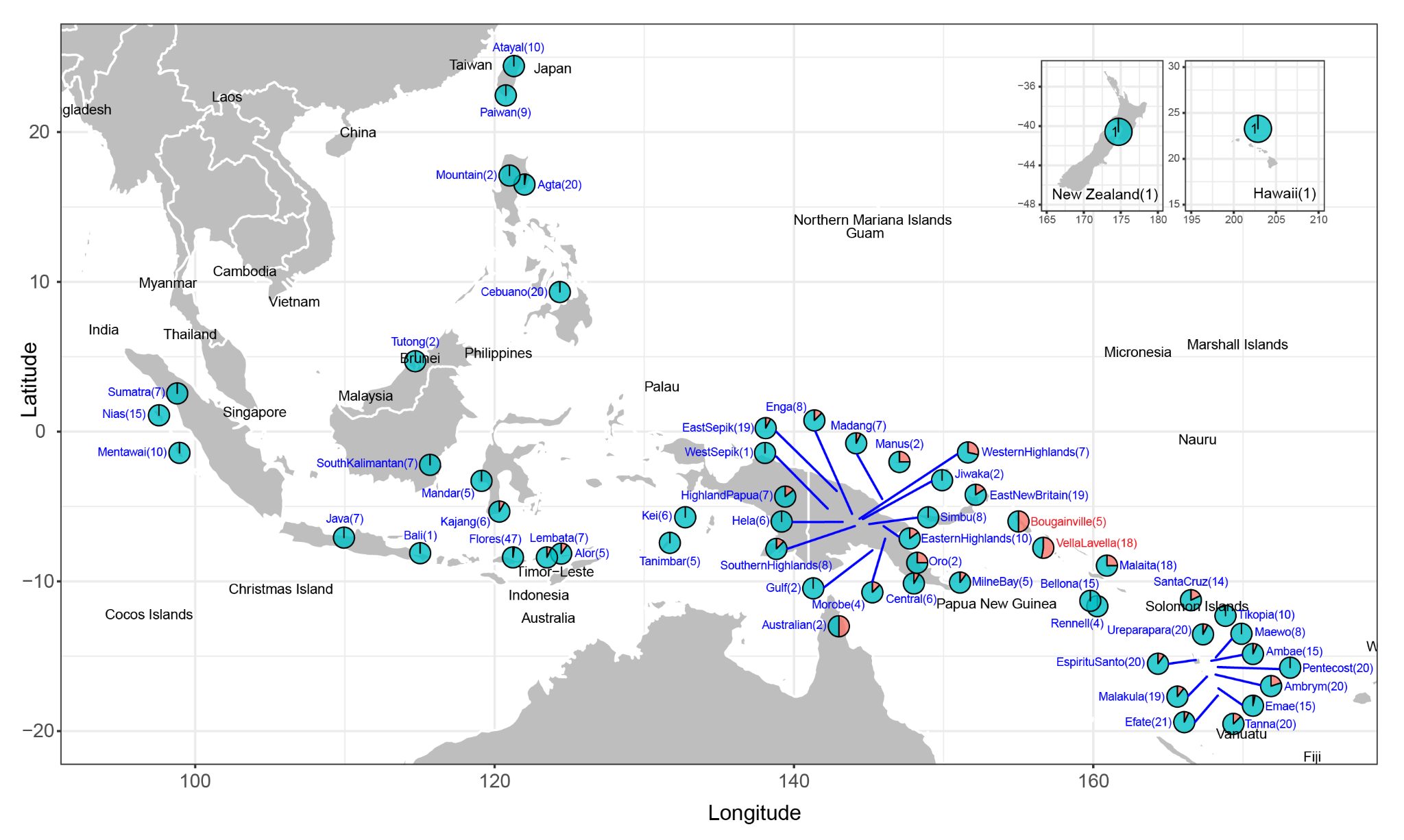

**Supplementary Figure 13. Regional distribution of Denisovan-derived *Alu* insertion in *OCA2* across Southeast Asia and Oceania.** This figure provides a zoomed-out view of Figure 2B, encompassing a broader geographic range, based solely on the Southeast Asian and Oceanian dataset. Numbers in parentheses represent the sample size with non-missing genotypes (genotype quality > 30) for each region.

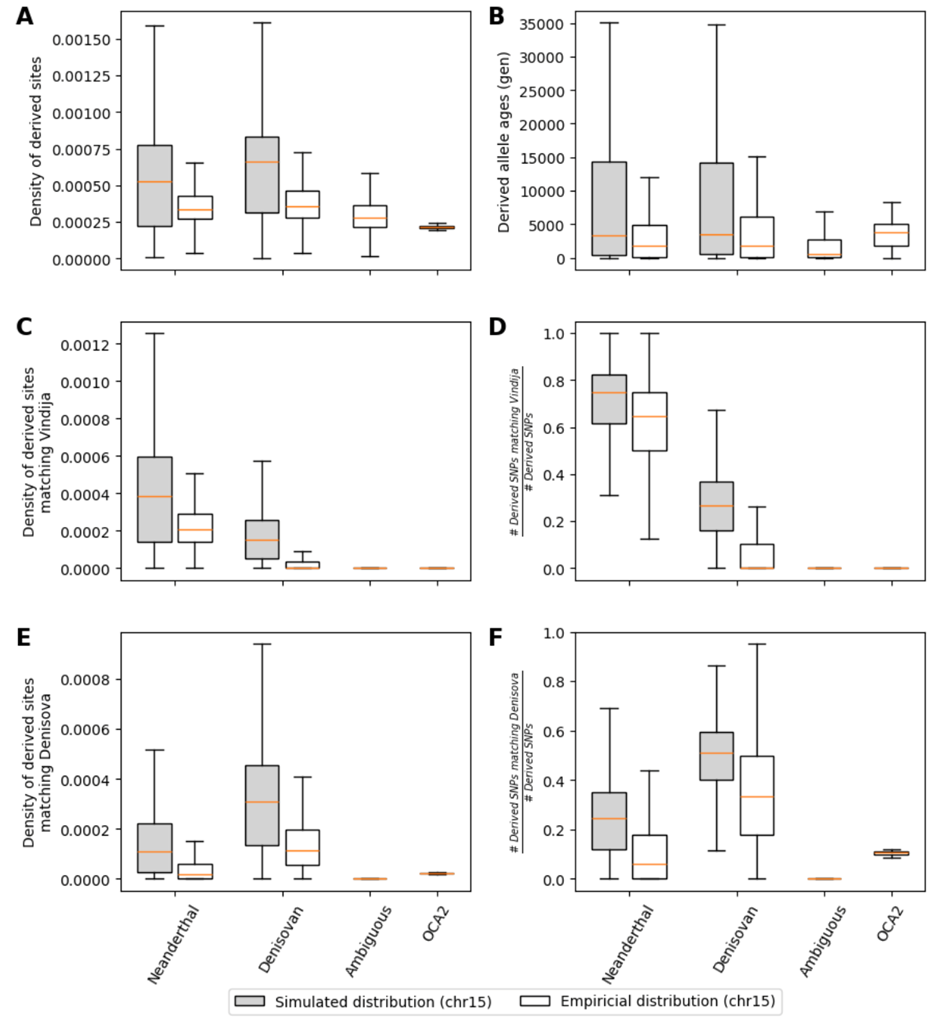

**Supplementary Figure 14. Empirical (white) and simulated (grey) distributions of derived alleles in introgressed segments compared to those of Denisovan-like haplotypes at the *OCA2* insertion locus.** Both empirical and simulated data show (a) comparable densities of derived alleles, (b) similar derived allele ages, similar derived allele densities matching the Vindija Neanderthal genome (c and d) and the Denisovan genome (e and f) to the Denisovan-like haplotypes overlapping the *OCA2* insertion.

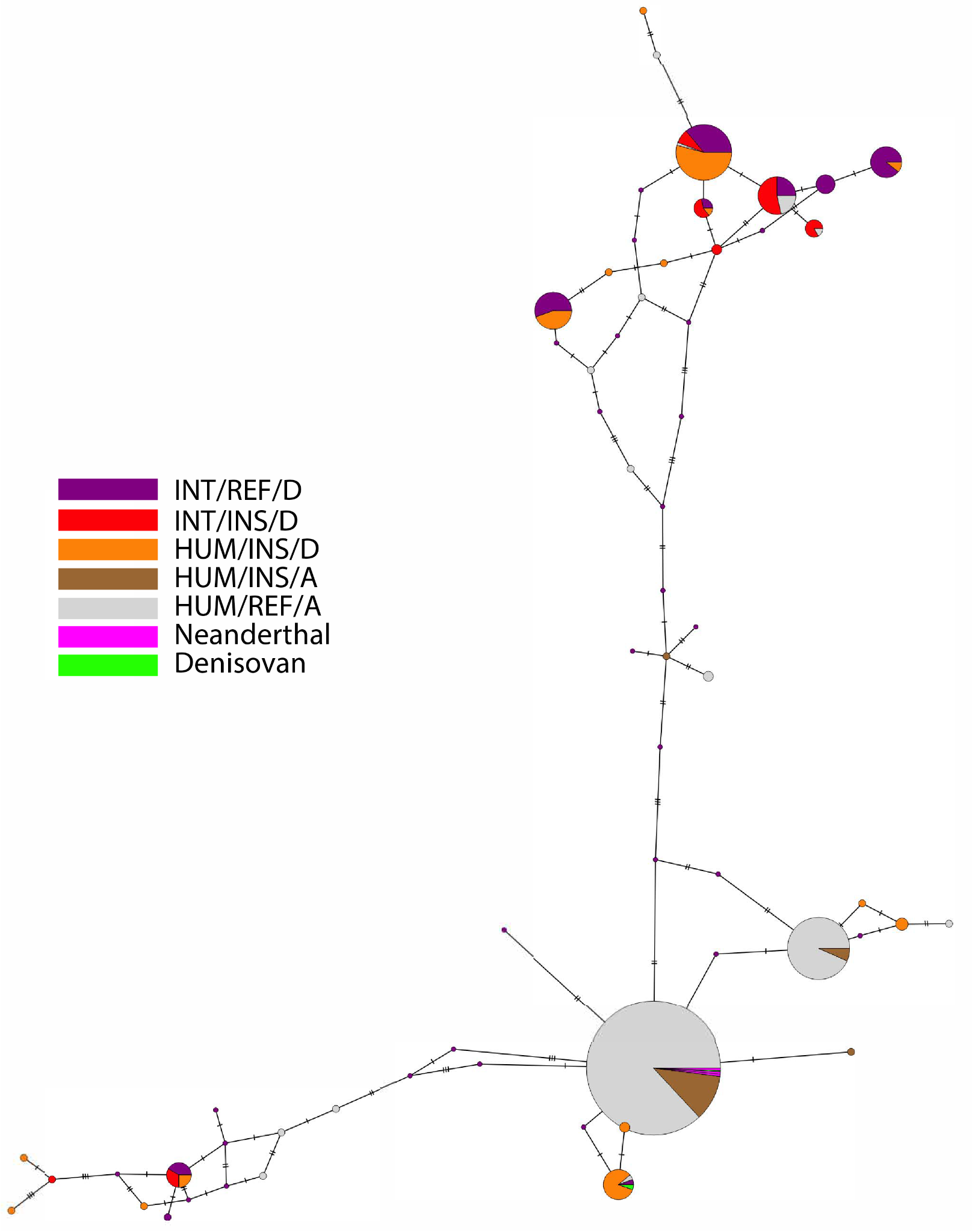

**Supplementary Figure 15. Minimum spanning tree haplotype network constructed from 24 introgression-informative sites identified by hmmix and the *Alu* insertion.** Haplotypes with the *Alu* insertion and the derived allele at chr15:27,934,733 (HUM/INS/D, orange) cluster closely with the Denisovan haplotypes (green) and introgressed haplotypes (INT/REF/D; purple and INT/INS/D; red), whereas haplotypes with the *Alu* insertion but carrying the ancestral allele at chr15:27,934,733 (HUM/INS/A, brown) do not.

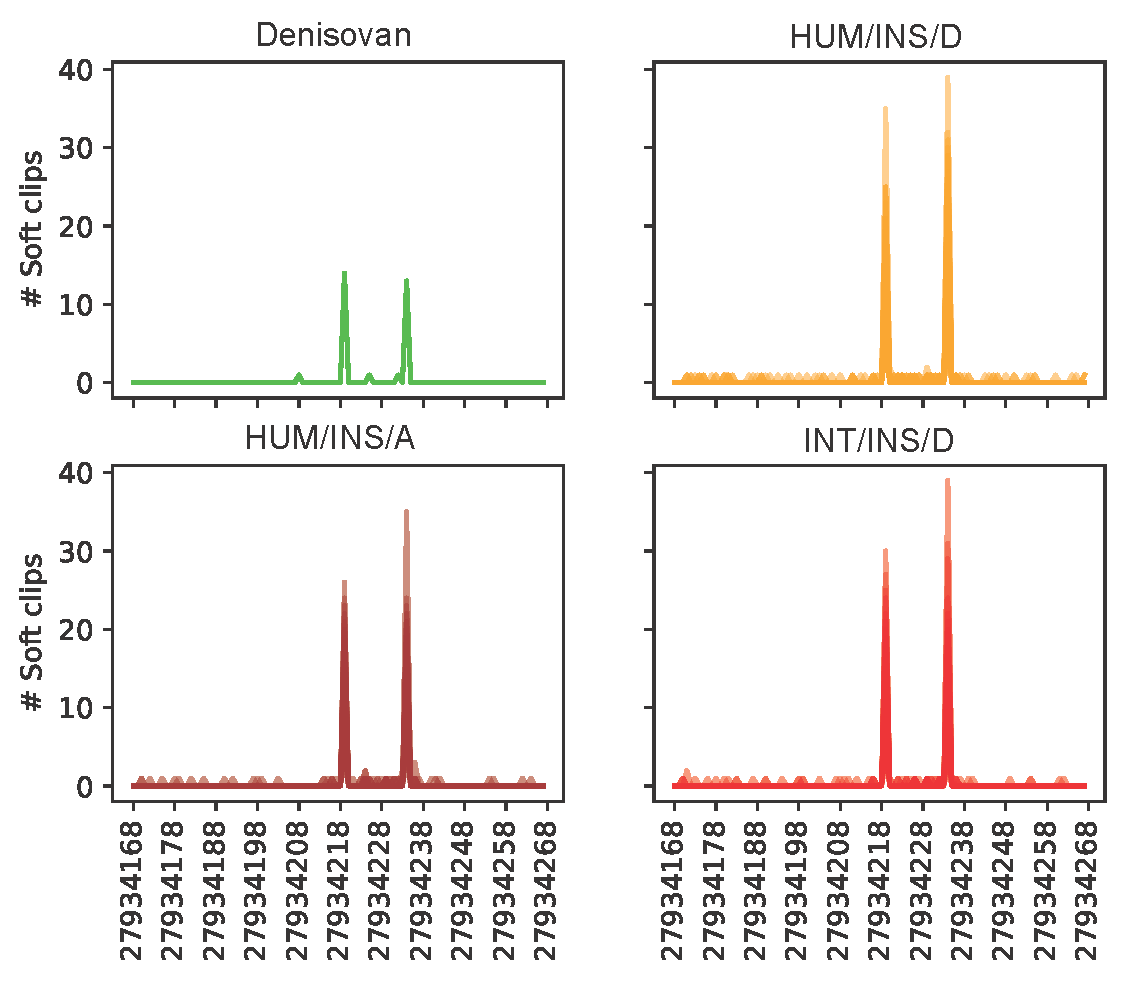

**Supplementary Figure 16. Breakpoints of soft-clipped reads aligned to the *Alu* insertion in the Denisovan genome, present-day human haplotypes with the *Alu* insertion (HUM/INS/D and HUM/INS/A), and predicted introgressed haplotypes with the *Alu* insertion (INT/INS/D).** All haplotypes represent the same breakpoint, supporting a common origin of the *Alu* insertion. The x-axis represents coordinates based on the GRCh38 reference genome.

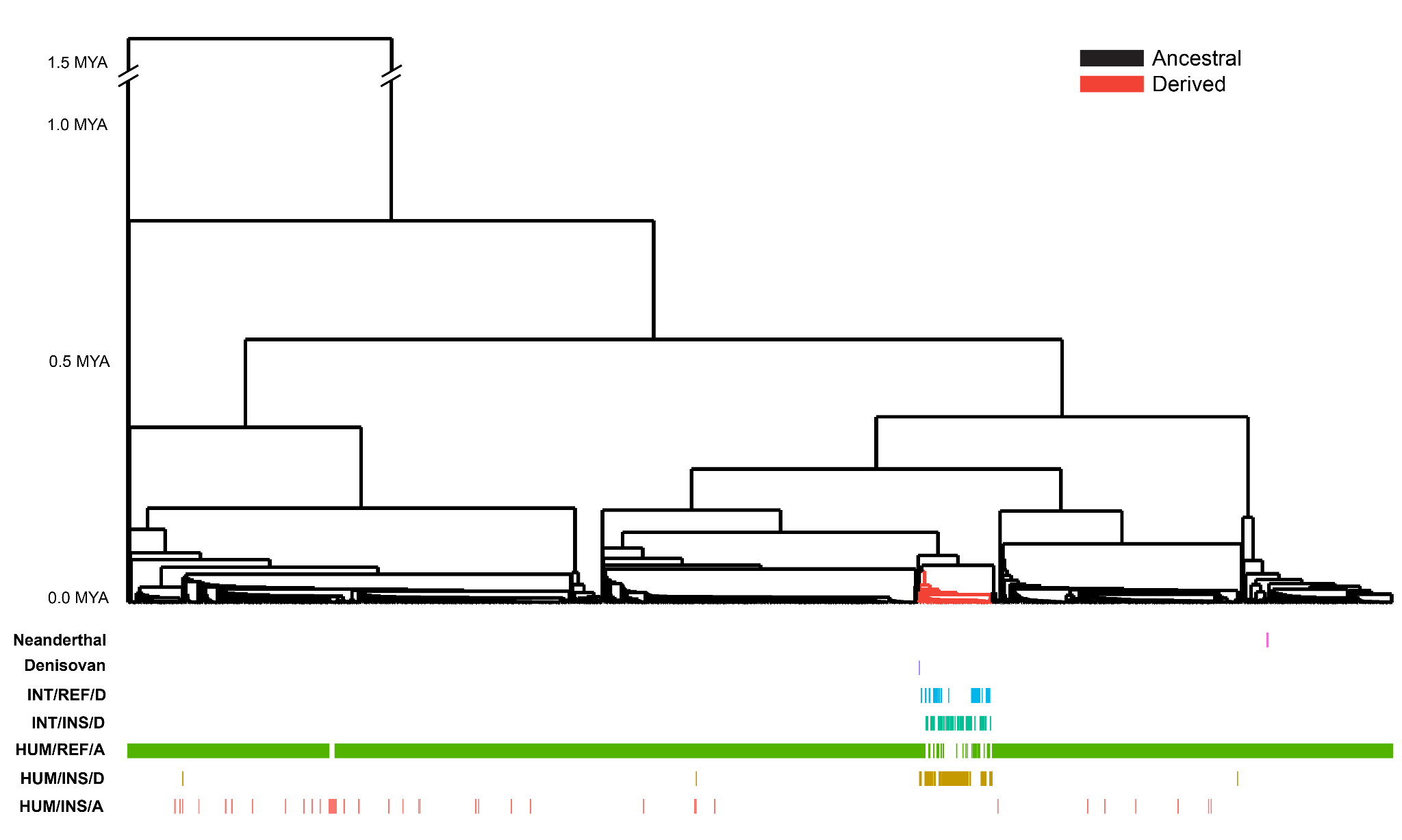

**Supplementary Figure 17. Marginal tree at chr15:27,934,733 inferred from the ancestral recombination graph using RELATE** ^3^**.** Group annotations correspond to those defined in Figure 3B.

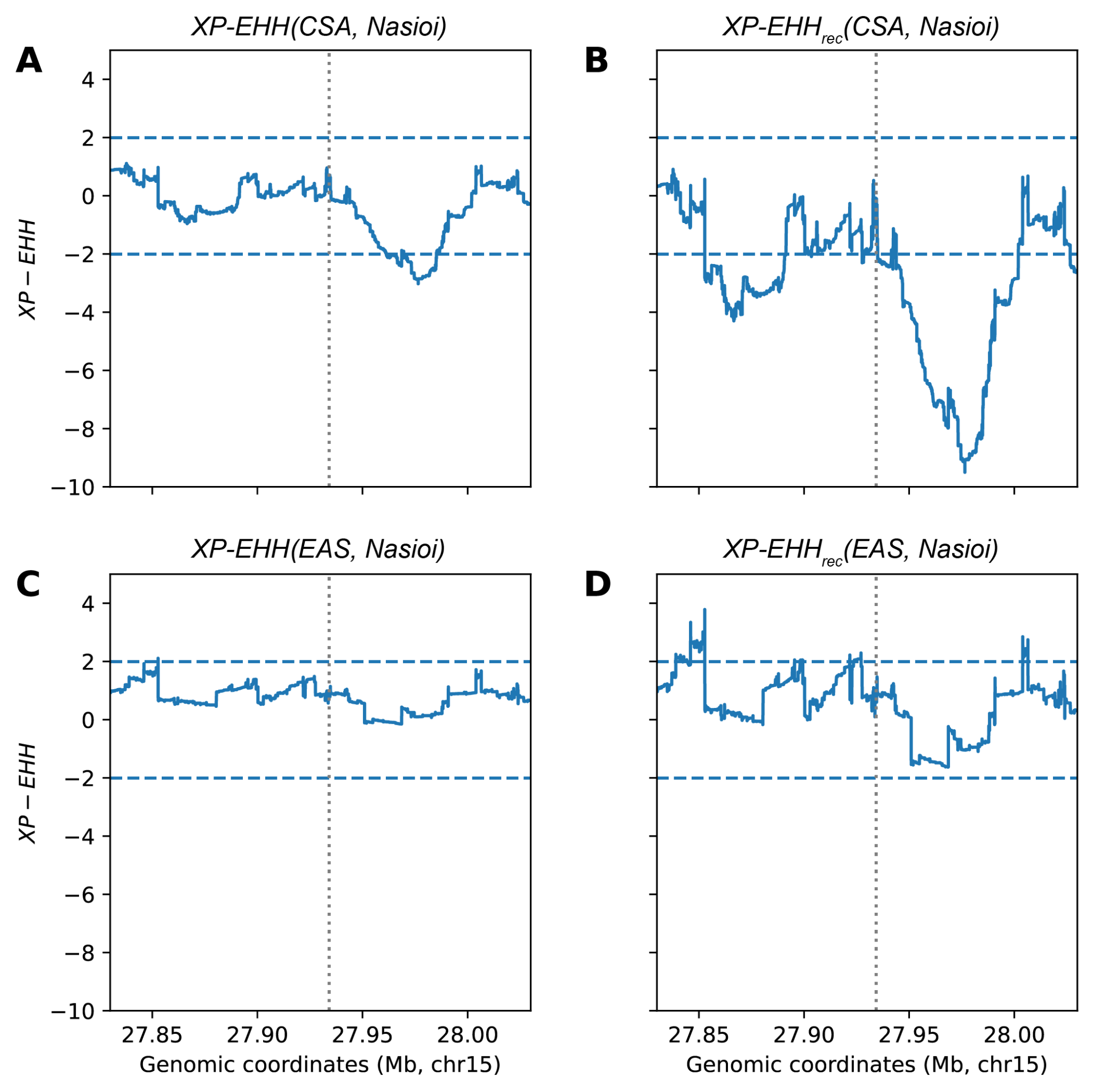

**Supplementary Figure 18. XP-EHH values of the Nasioi population of Bougainville around the *Alu* insertion using the Central and South Asian (CSA) and East Asian (EAS) superpopulations as a reference.** (a) When normalizing raw XP-EHH values relative to the genomic background, only a short stretch of approximately 40 kb downstream of the *Alu* insertion locus reaches significance (mean *XP-EHH(CSA, Nasioi)* = -0.66 in a 100 kb window). (b) When normalizing raw XP-EHH values relative to sites in the top 1% in terms of recombination rate, the $\pm$ 50 kb region becomes a significant outlier (mean *XP-EHH_rec_(CSA, Nasioi)* = -3.55 in a 100 kb window). (c) When normalizing raw XP-EHH values relative to the genomic background, no significant signal (> 2 or < -2) was detected at the *Alu* insertion site (mean *XP-EHH(EAS, Nasioi)* = 0.65 in a 100 kb window). (d) When normalizing raw XP-EHH values relative to sites in the top 1% in terms of recombination rate, no significant signal (> 2 or < -2) was detected at the *Alu* insertion site (mean *XP-EHH(EAS, Nasioi)* = 0.29 in a 100 kb window).

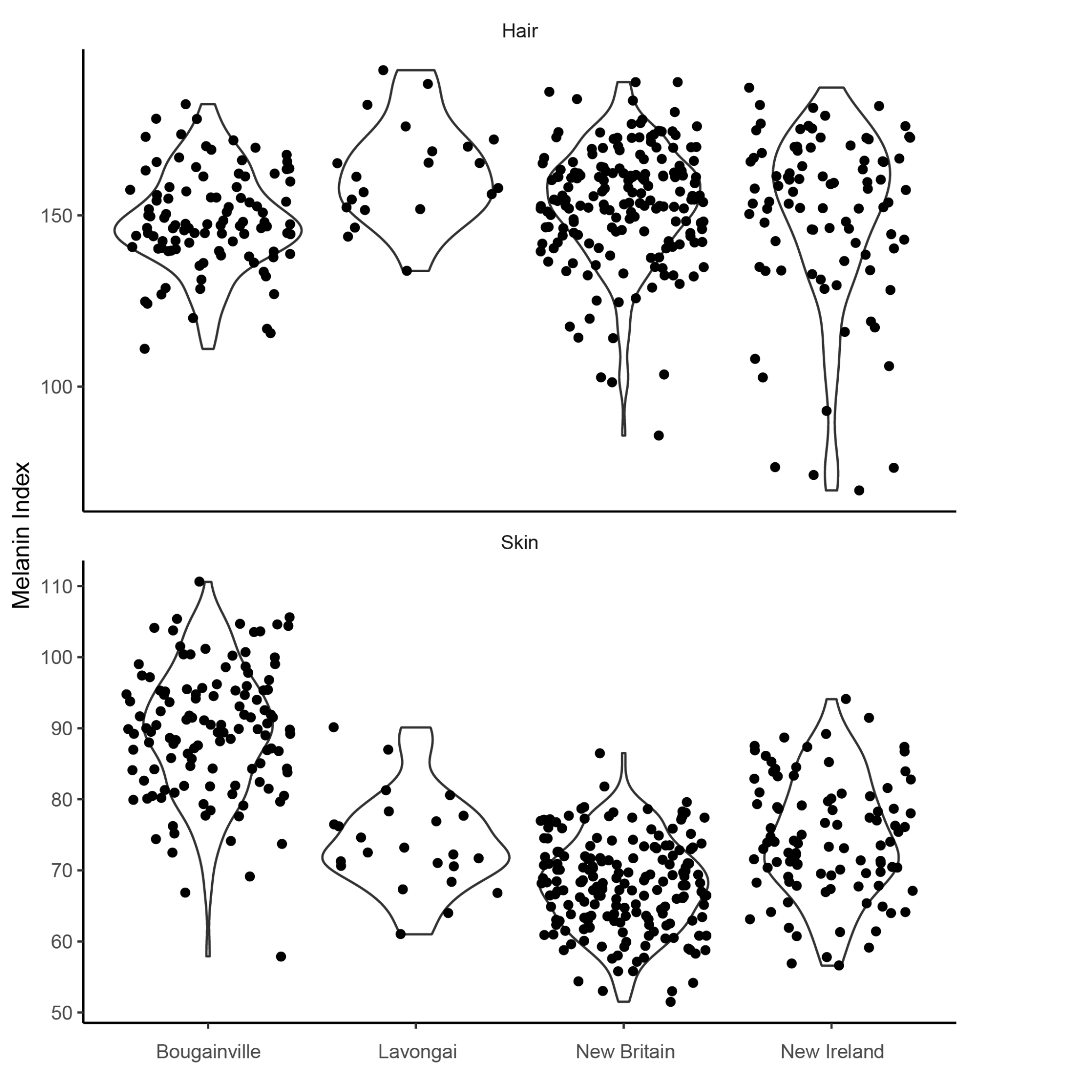

**Supplementary Figure 19. Hair and skin melanin indices on the islands of Bougainville, Lavongai, New Britain, and New Ireland.**

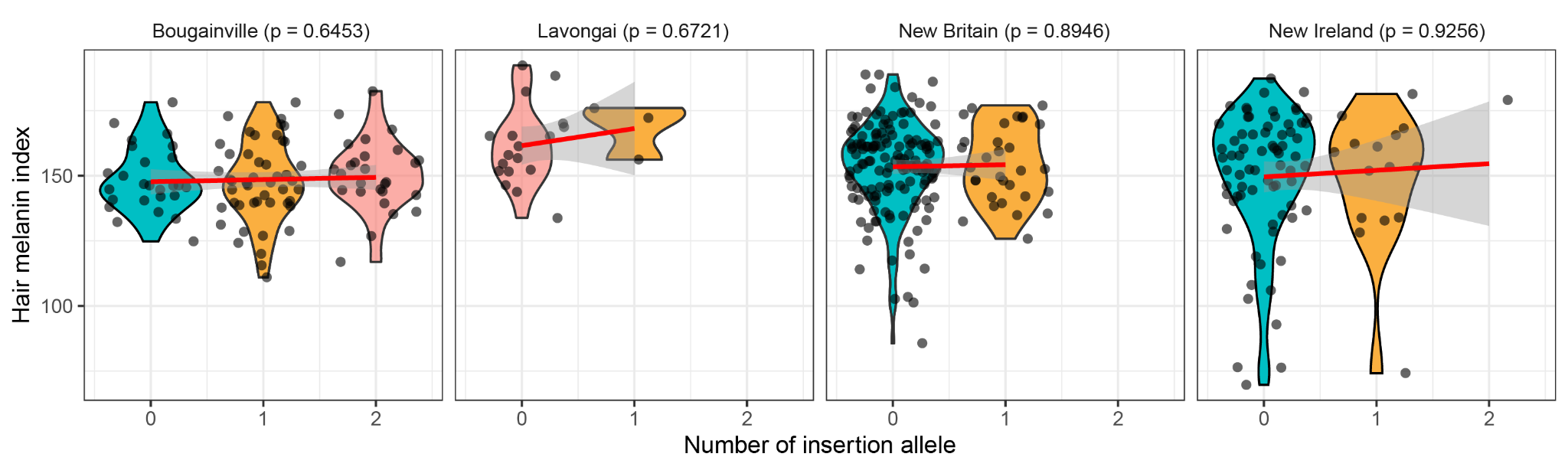

**Supplementary Figure 20. Association between the number of *Alu* insertion alleles and hair melanin index across four islands, Bougainville, Lavongai, New Britain, and New Ireland.** The red line is generated by the geom_smooth() function implemented in ggplot2 ^4^ using the formula = y ~ x.

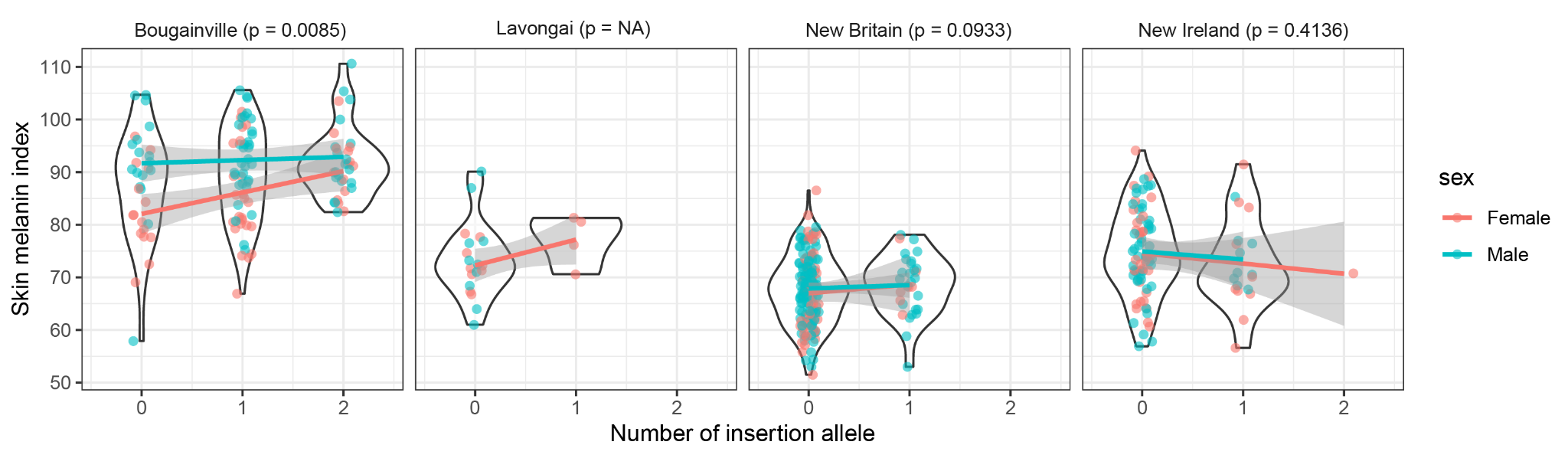

**Supplementary Figure 21. Interaction effect between sex and the number of *OCA* *Alu* insertions on skin melanin index.** A stronger association is observed in females than in males in Bougainville Island (*p-value* = 8.498 $\times$ 10^-3^). The interaction test was not available in Lavongai, as all males in this island have the same homozygous reference genotype. The trend line is generated by the geom_smooth() function implemented in ggplot2 ^4^ using the formula = y ~ x.

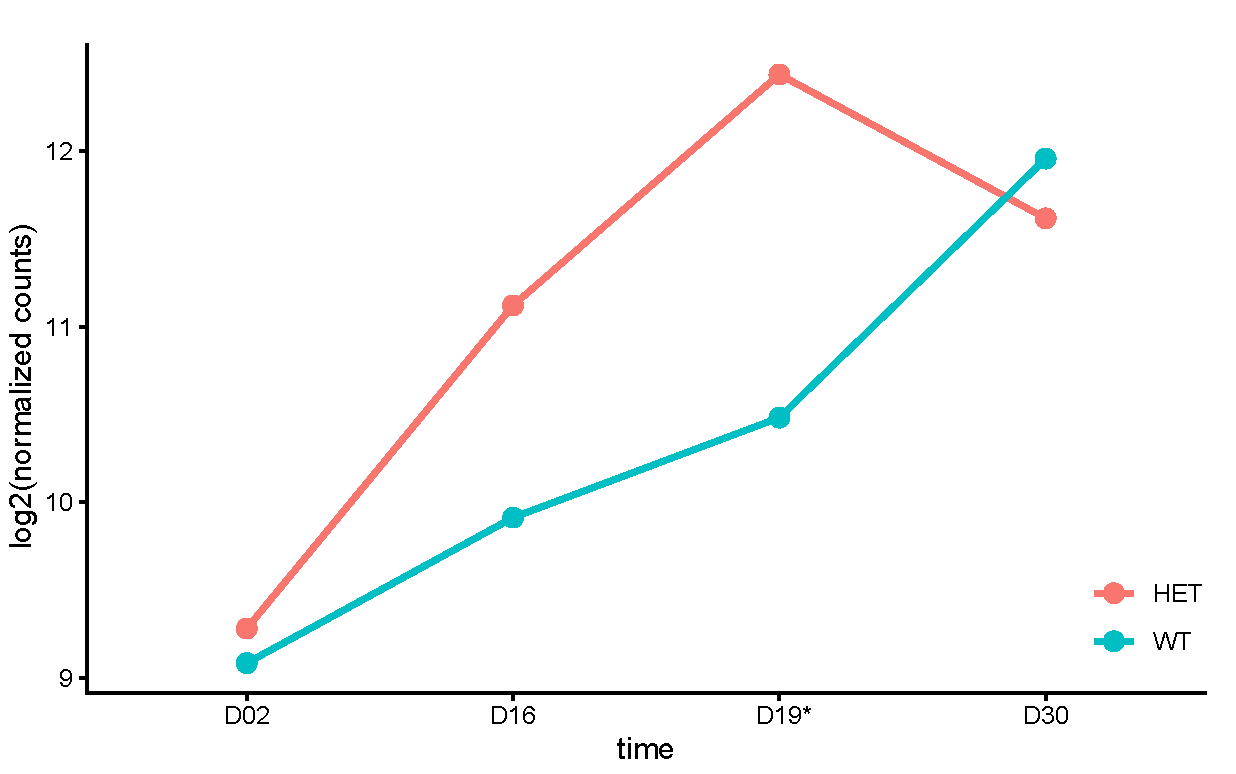

**Supplementary Figure 22. *OCA2* RNA-seq expression across time points, day 2, 16, 19, and 30.** The asterisk (*) indicates the time point where HET (heterozygous) melanocyte expression is significantly higher than that of WT (wild-type) (*p-value* < 0.05).

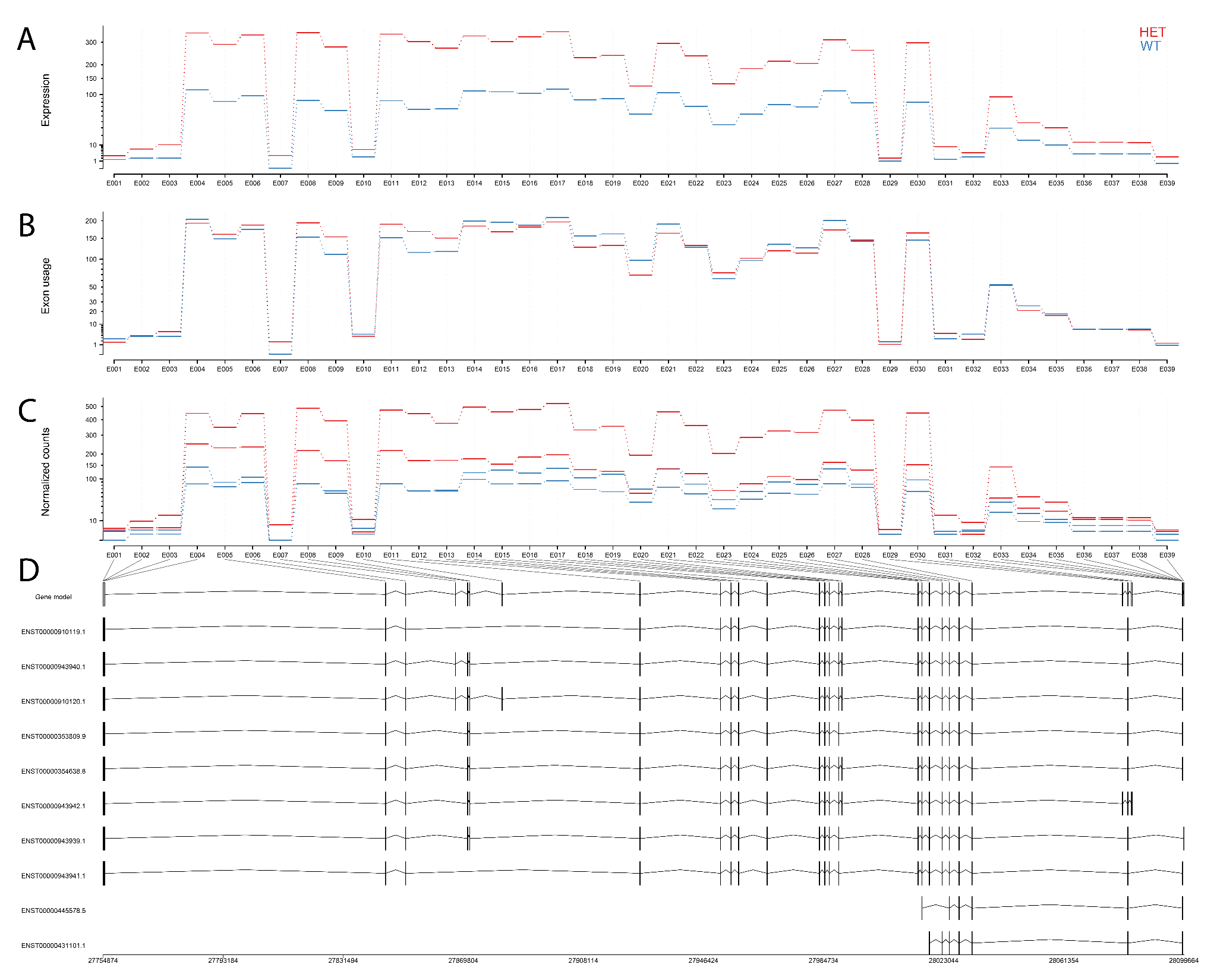
 **Supplementary Figure 23. Result from differential exon usage analysis of *OCA2*.** (a) Fitted expression estimates for HET (heterozygous) and WT (wild-type) melanocytes. (b) Fitted expression estimates of each exon for HET and WT melanocytes. (c) Normalized counts for each sample. (d) Transcript models tested.

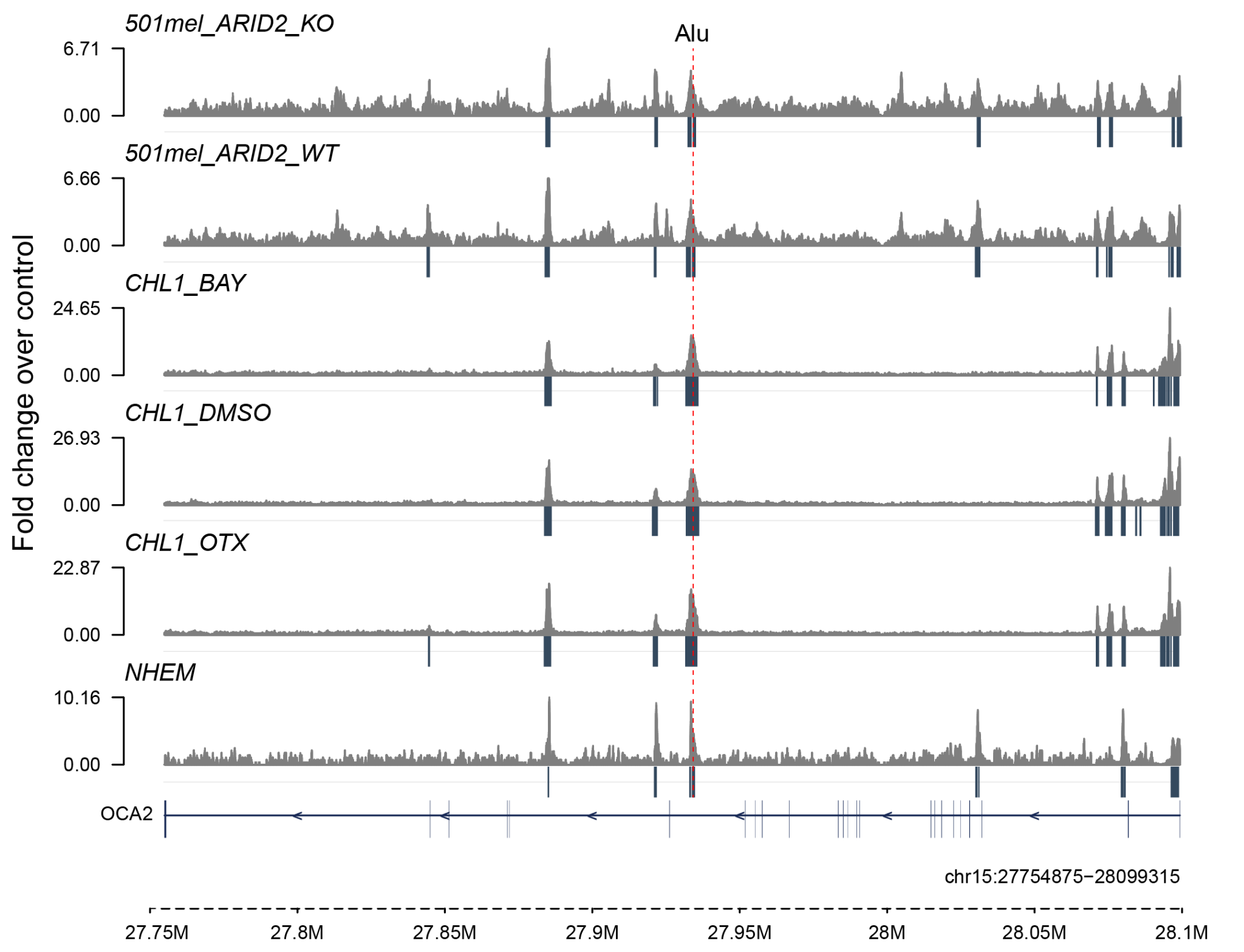

**Supplementary Figure 24. H3K27ac peaks from melanoma and melanocyte cell lines (501 mel, CHL1, and NHEM).** The y-axis shows fold change relative to the input control. The black bars under the fold change indicate H3K27ac peak intervals. The dashed red line indicates the genomic position of the *Alu* insertion. 501mel_ARID2_KO: ARID2 knock-out cells (SAMN18806624, SAMN18806625), 501mel_ARID2_WT: wild-type cells (SAMN18806626, SAMN18806627), CHL1_BAY: CHL1 cells treated with BAY 1238097 (SAMN06470131, SAMN06470146), CHL1_DMSO: CHL1 cells without treatment (SAMN06468140, SAMN06470136), CHL1_OTX: CHL1 cells treated with OTX-015 (SAMN06470126, SAMN06470141), NHEM: Normal human epithelial melanocytes (SAMN42682903).

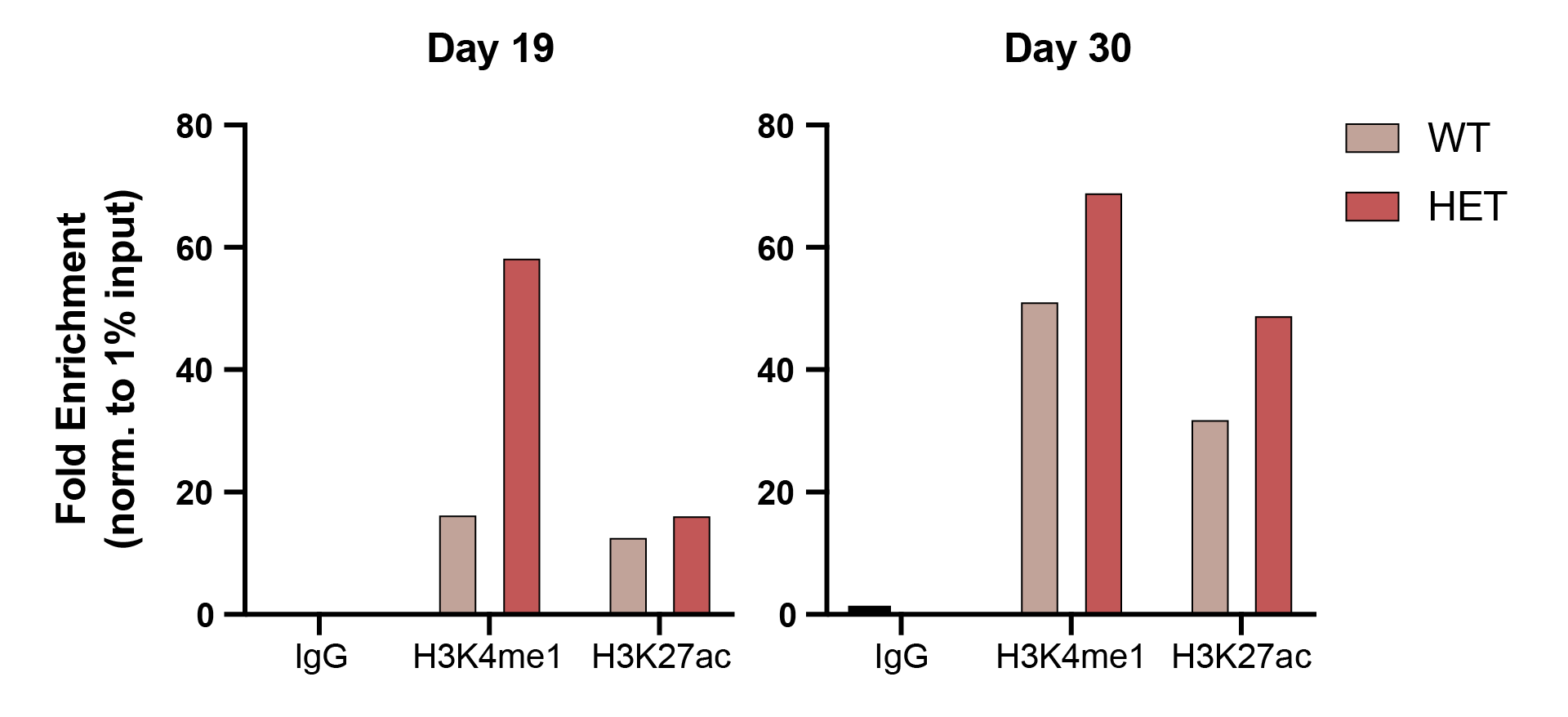

**Supplementary Figure 25. Histone mark (H3K4me1 and H3K27ac) enrichments across two time points, day 19 and 30, using a primer set (huCh_Oca2alu-right_aluK27ac_F1 and huCh_Oca2alu-right_aluK27ac_R1).** “IgG” indicates a negative control for the enrichment assay. Fold enrichment values were normalized to 1% of input.

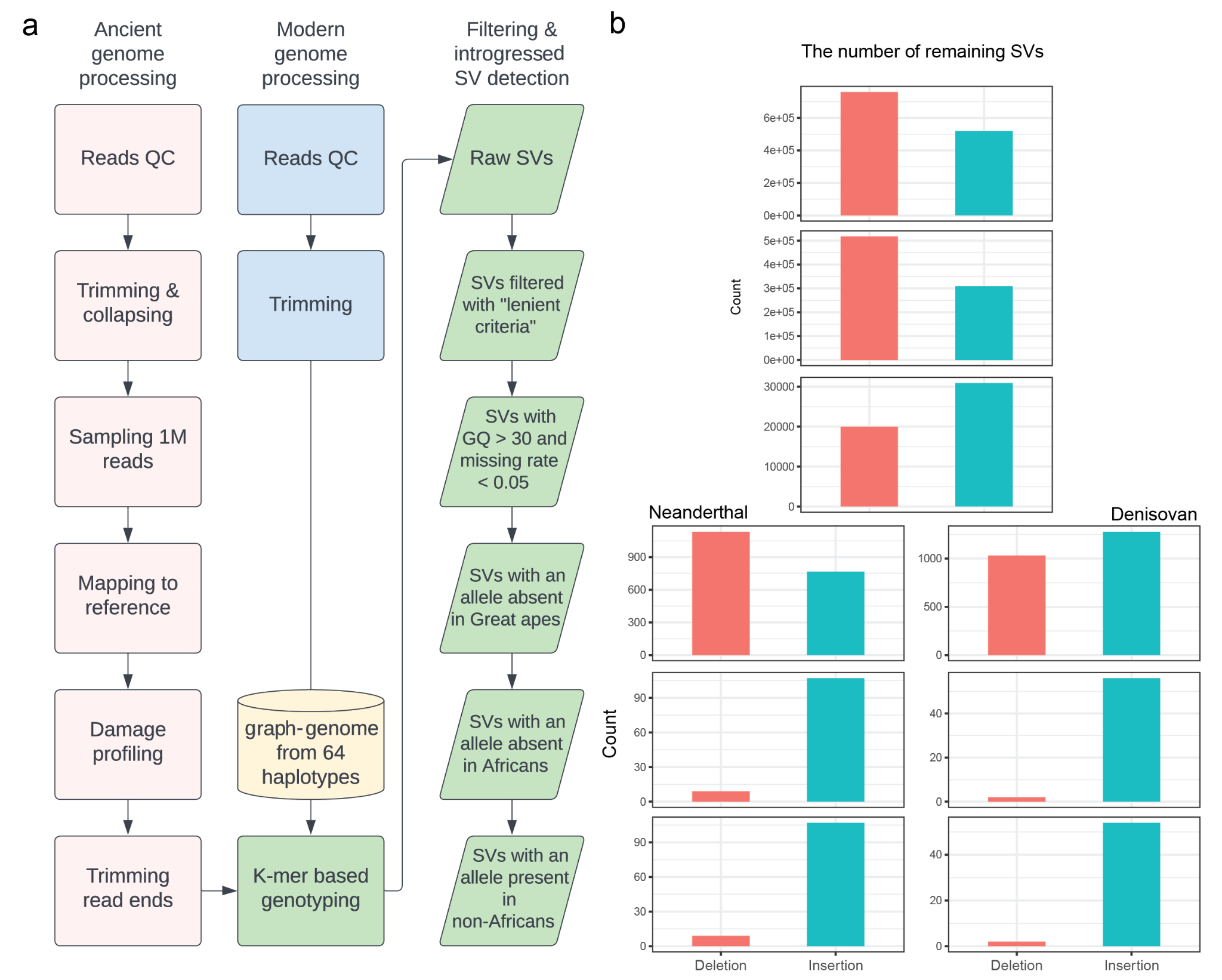

**Supplementary Figure 26. Workflow to detect introgressed structural variants (SVs).** (a) The workflow ranges from preprocessing raw sequence reads to the detection of introgressed SVs. (b) The number of remaining SVs in each step of the “Filtering & introgressed SV detection” procedure in panel (a).

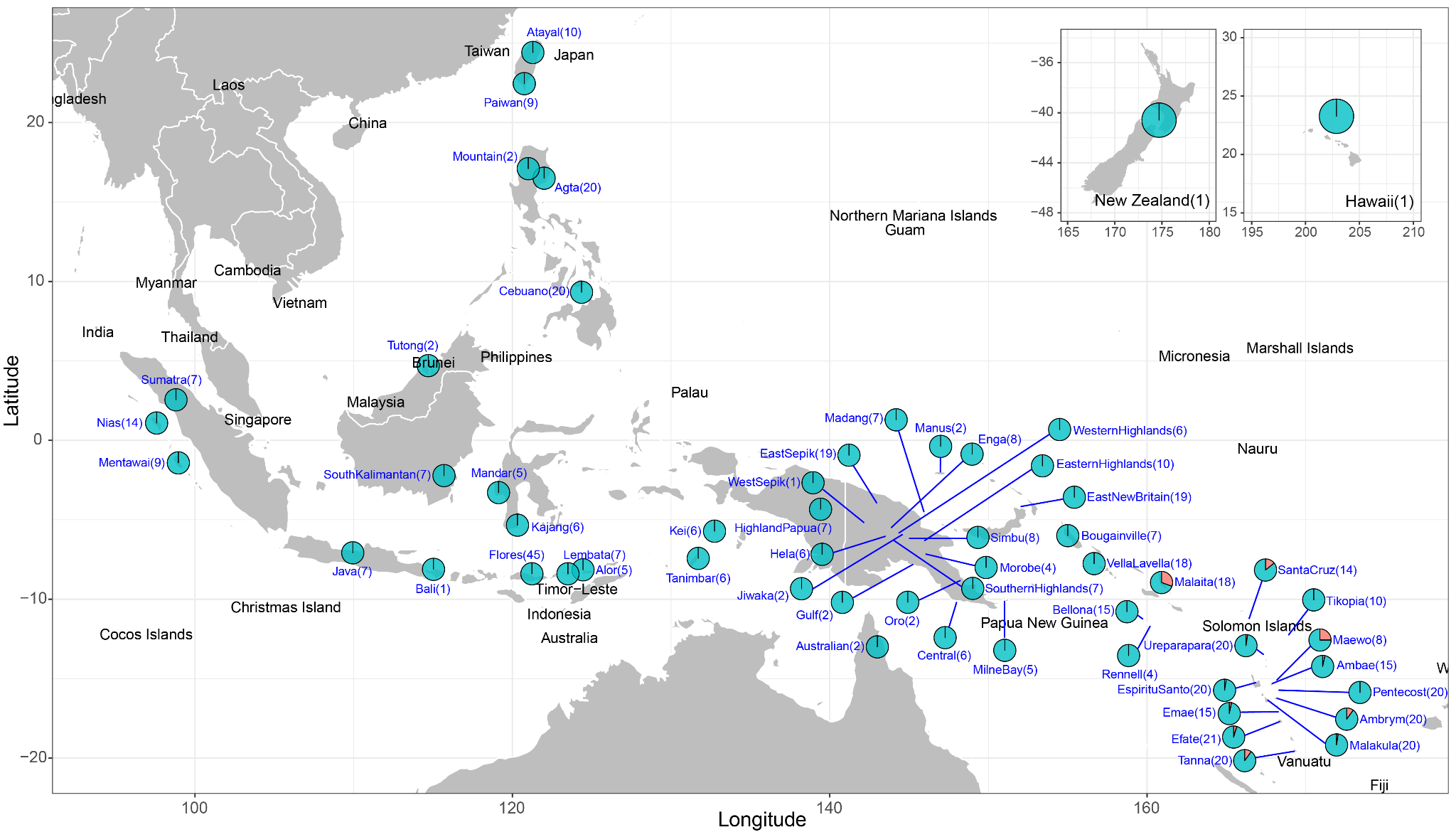

**Supplementary Figure 27. Distribution of rs387907171 in *TYRP1* around the Southeast Asian and Oceanian area.** The number in parentheses indicates the sample size with non-missing genotype (genotype quality > 30) for each region. The red color indicates the frequency of rs387907171 in each region.

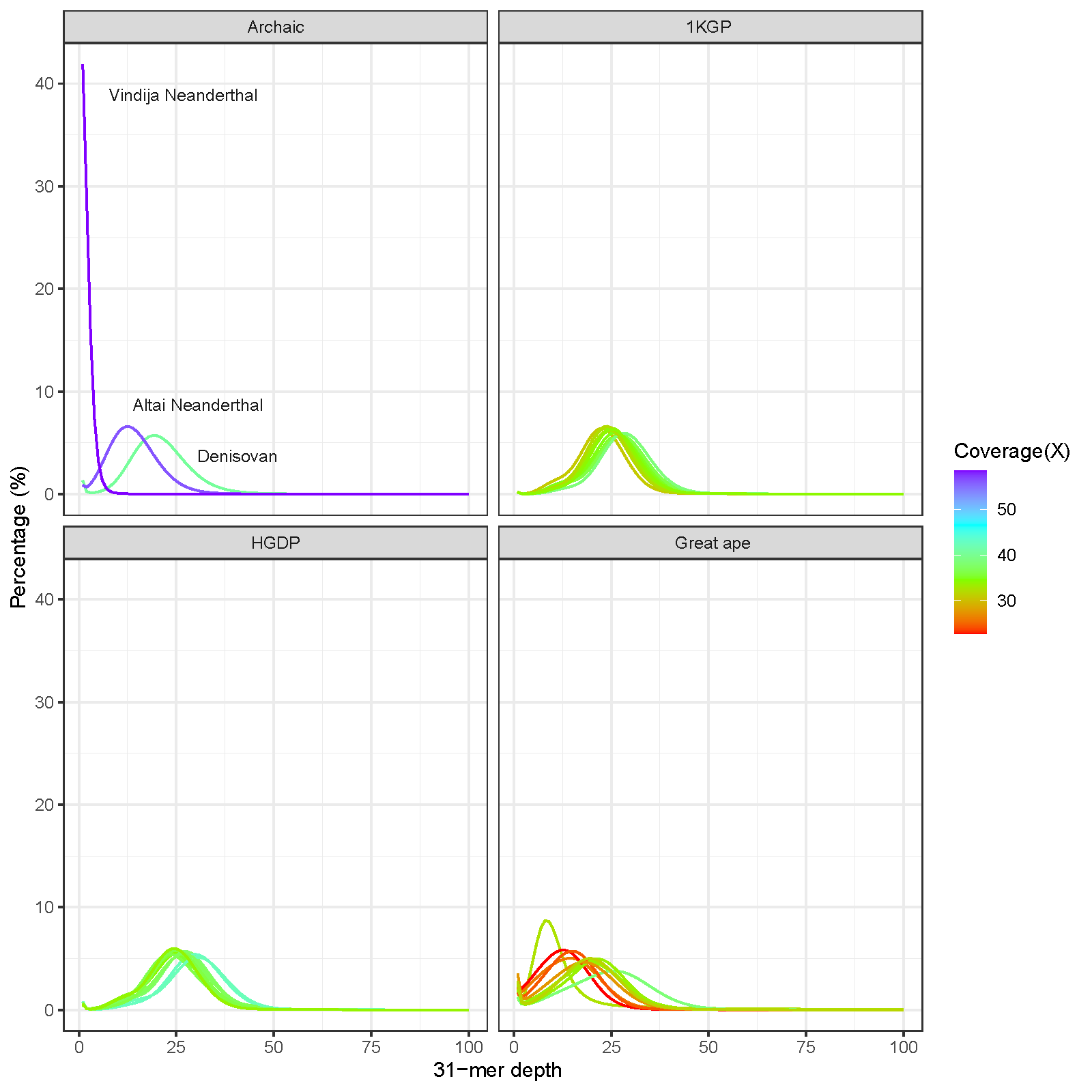

**Supplementary Figure 28.** **31-mer distributions of the genomes used in this study.** Archaic: Archaic hominin genomes, 1KGP: Randomly selected 10 individuals from the 1000 Genomes Project dataset, HGDP: Randomly selected 10 individuals from the Human Genome Diversity Project dataset, and Great ape: Randomly selected 10 individuals from the great apes dataset.

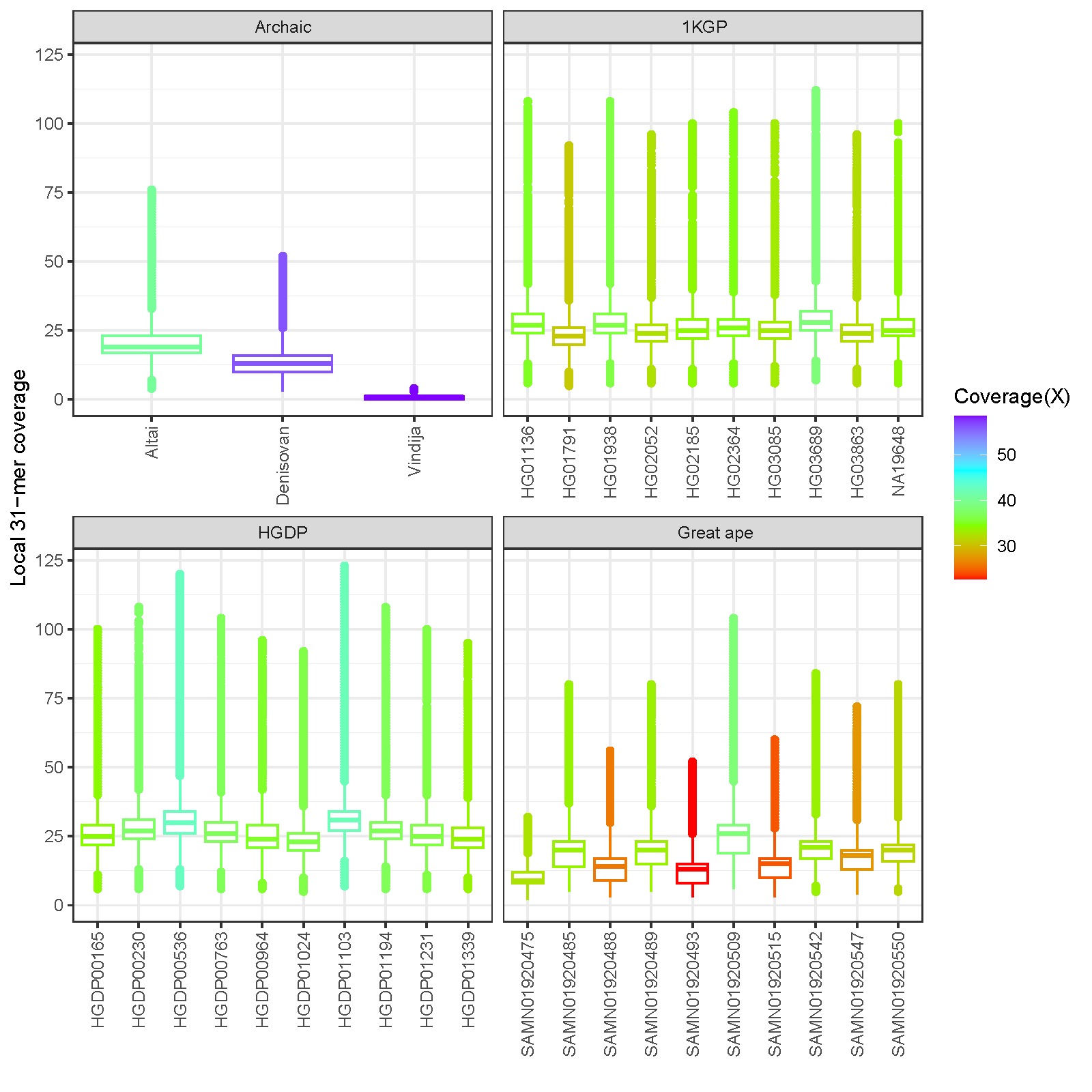

**Supplementary Figure 29.** **The number of 31-mers used for genotyping at each variant locus.** Archaic: Archaic hominin genomes, 1KGP: Randomly selected 10 individuals from the 1000 Genomes Project dataset, HGDP: Randomly selected 10 individuals from the Human Genome Diversity Project dataset, and Great ape: Randomly selected 10 individuals from the great apes dataset.

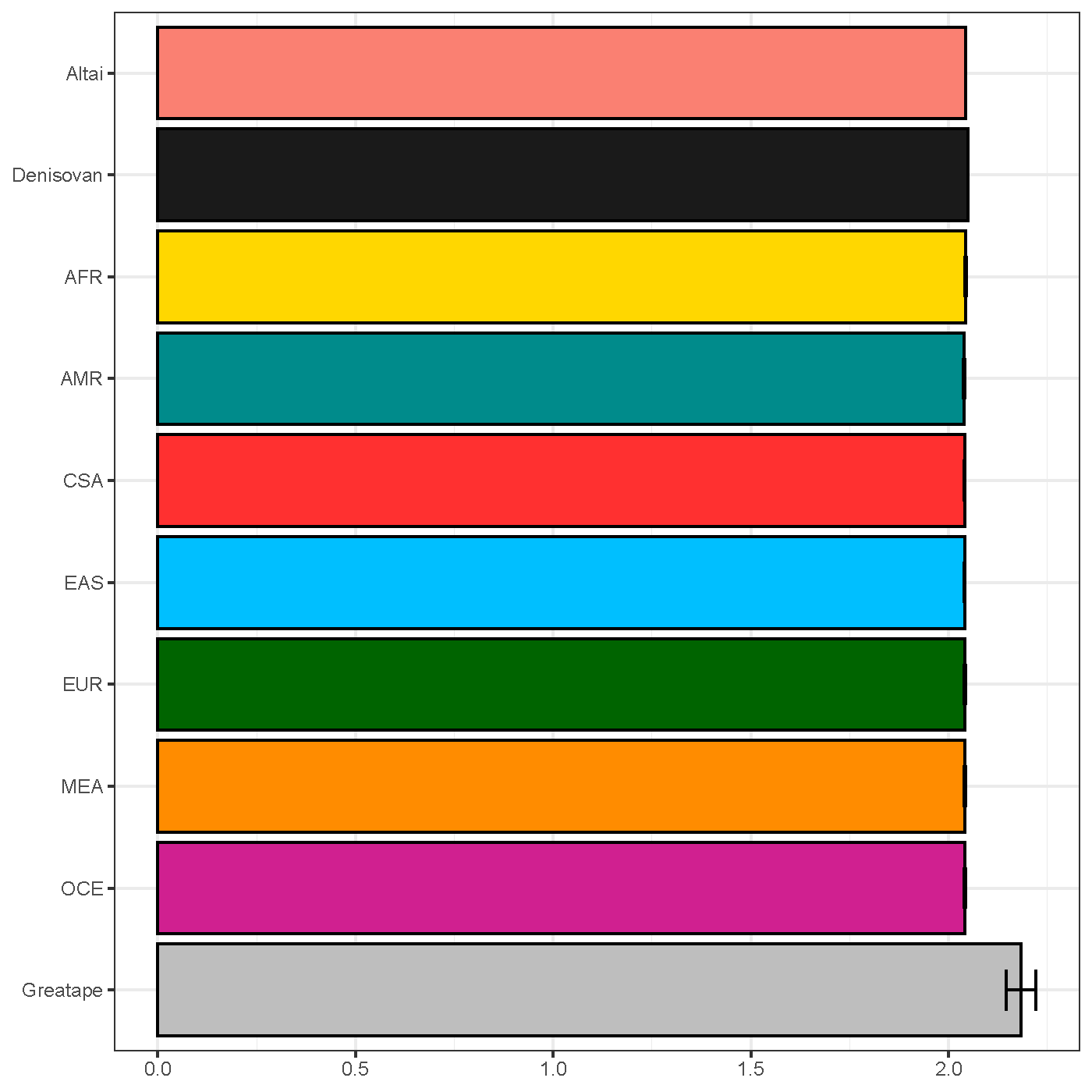

**Supplementary Figure 30. Transitions-to-transversions (Ts:Tv) ratio estimates based on PanGenie genotype calls.** The error bar indicates $\pm$ one standard deviation. Altai: Altai Neanderthal, AFR: Africa, AMR: America, CSA: Central and South Asia, EAS: East Asia, EUR: Europe, MEA: Middle East, OCE: Oceania, and Greatape: great apes.

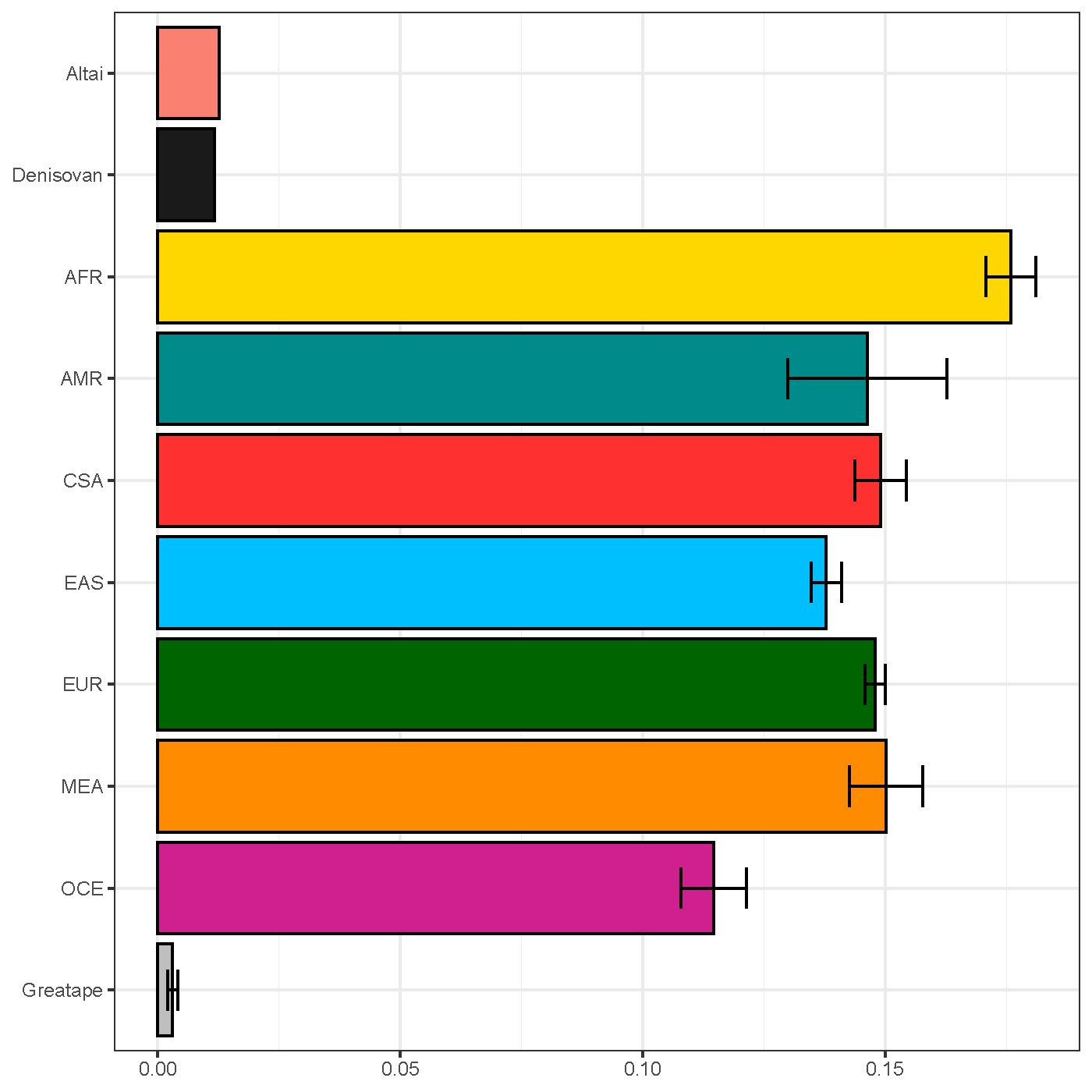

**Supplementary Figure 31. Heterozygosity estimates based on PanGenie genotype calls.** The error bar indicates $\pm$ one standard deviation. Altai: Altai Neanderthal, AFR: Africa, AMR: America, CSA: Central and South Asia, EAS: East Asia, EUR: Europe, MEA: Middle East, OCE: Oceania, and Greatape: great apes.

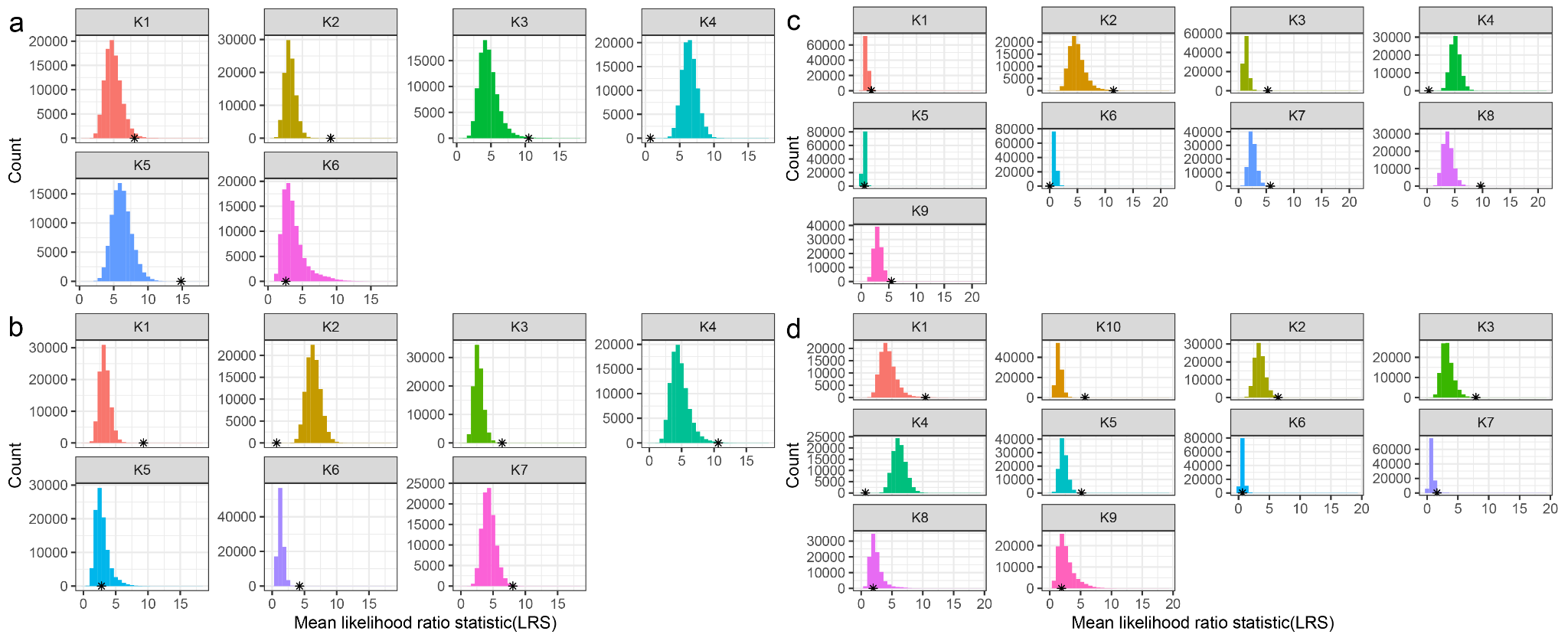

**Supplementary Figure 32. Overall selection signatures of introgressed structural variants (SVs) for each ancestry component under (a) K=6, (b) K=7, (c) K=9, and (d) K=10 models.** The asterisk (*) indicates the mean likelihood ratio statistic of the introgressed SVs.

Supplementary Figure 33

**Supplementary Figure 33.** **Individual ancestry proportions estimated by Ohana with (a) K=6, (b) K=7, (c) K=9, and (d) K=10**. AFR: Africa, AMR: America, CSA: Central and South Asia, EAS: East Asia, EUR: Europe, MEA: Middle East, and OCE: Oceania

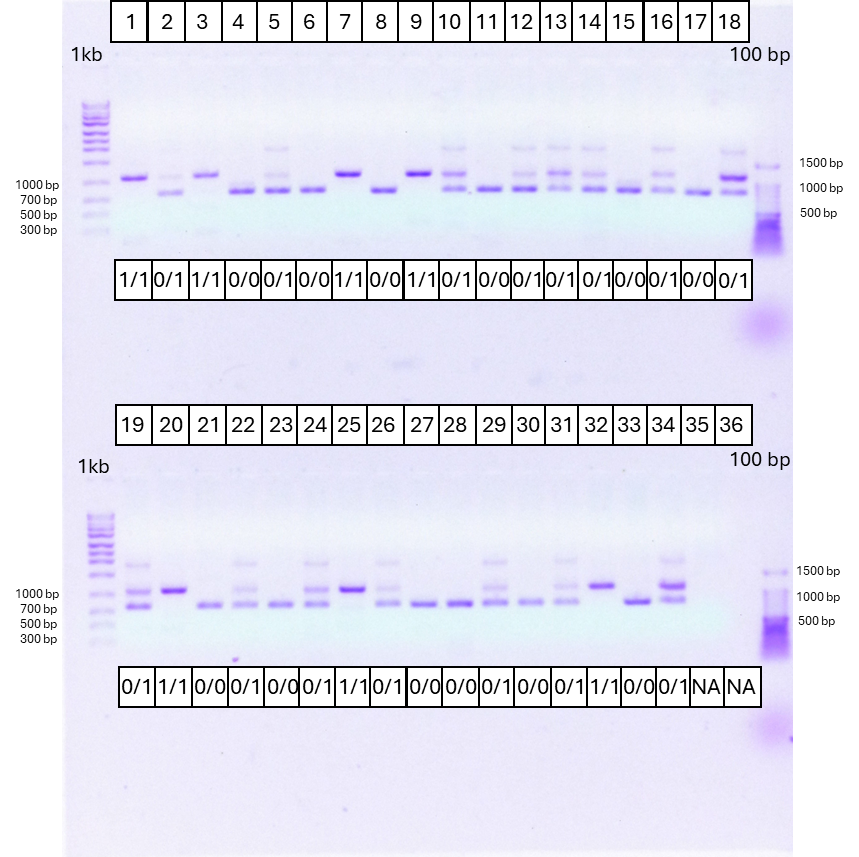

**Supplementary Figure 34. Representative gel image for genotyping PCR of *OCA2* *Alu* insertion.** Well #33 is a control for the homozygous wild-type. Well #34 is a control sample for the heterozygote. Well #35 is a negative control. No sample was loaded in well #36. 0/0: no *Alu* insertion, 0/1: heterozygote, 1/1: homozygote for the *Alu* insertion.

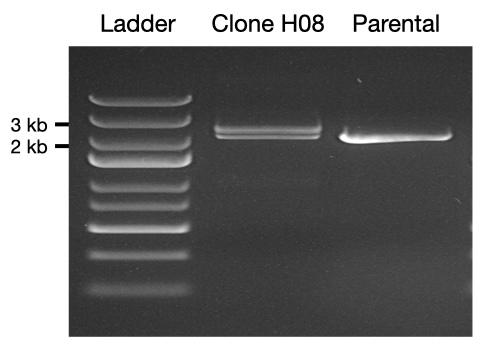

**Supplementary Figure 35. Confirmation of CRISPR-mediated insertion of a 332 bp *Alu* element into KOLF2.1J hiPSCs.** Gel electrophoresis of PCR amplicons spanning the CRISPR target site shows three lanes: Lane 1 contains a 1 kb DNA ladder; Lane 2 contains clone H08, which displays two bands - 2077 bp representing the wild-type (WT) allele and 2409 bp representing the *Alu* insertion (INS) allele - indicating a heterozygous *Alu* insertion; Lane 3 contains parental KOLF2.1J cells, showing a single 2077 bp band consistent with a biallelic WT genotype and absence of INS. Sequencing of clone H08 by both Sanger and NGS confirmed the presence of WT and INS alleles. The WT allele in clone H08 also contains an unintended 11 bp deletion at the CRISPR cut site, likely resulting from non-homologous end joining (NHEJ) repair.

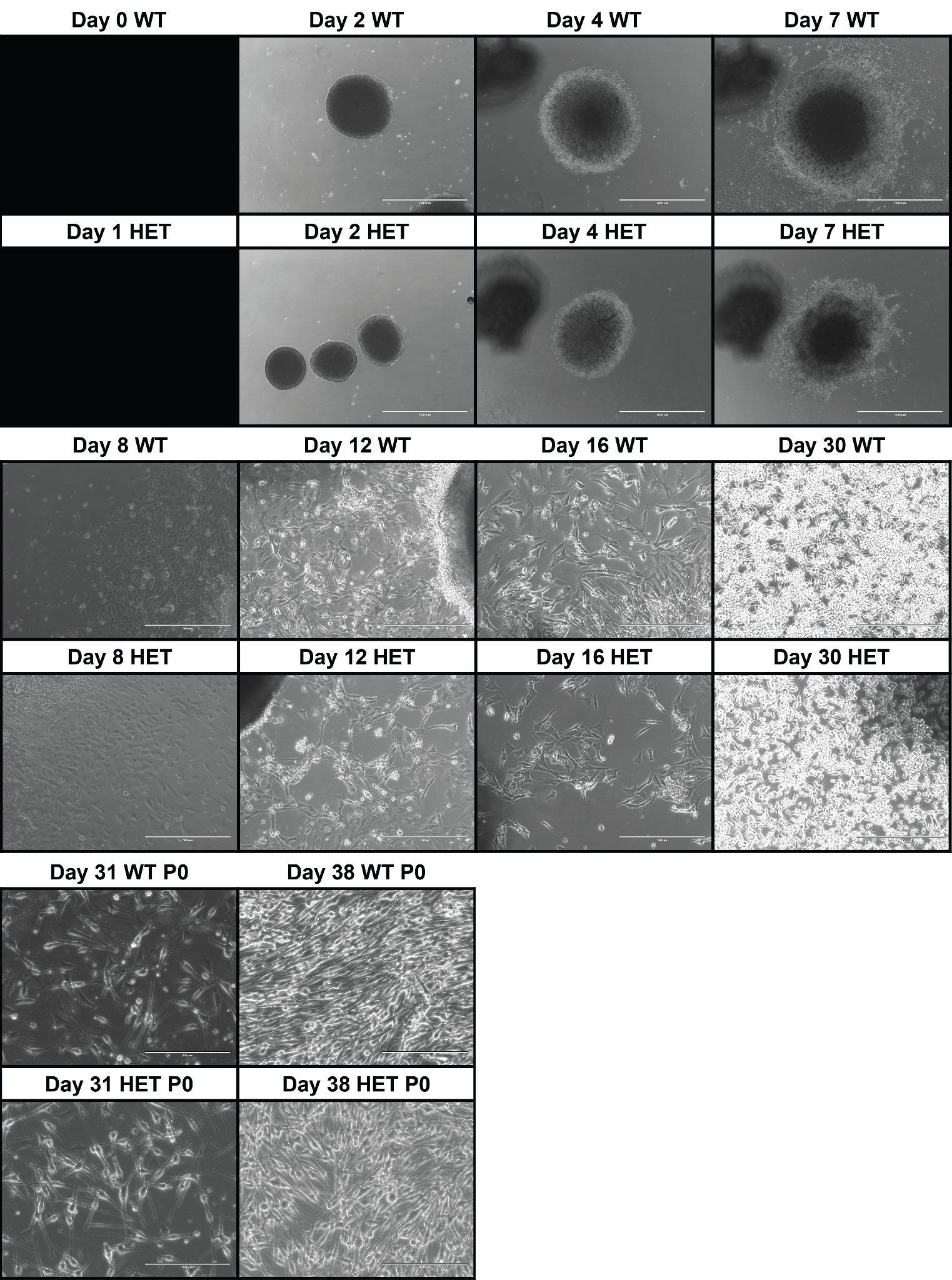

**Supplementary Figure 36. Microscopy images of wild-type (WT) and heterozygote (HET) cells for Denisovan-derived *Alu* insertion in *OCA2* at eight time points during differentiation and passage 0 (P0).**

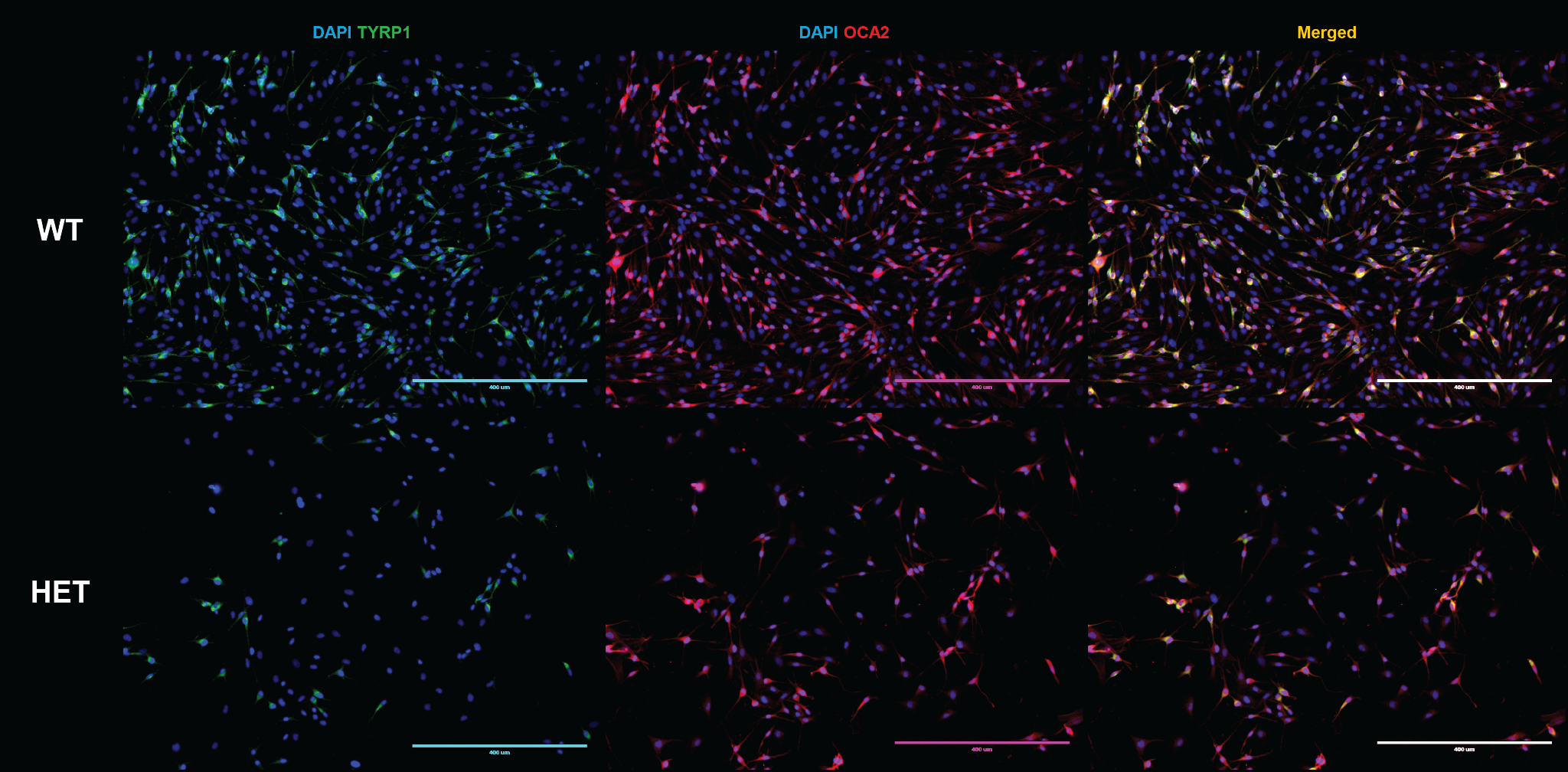

**Supplementary Figure 37. Immunofluorescence staining of *TYRP1* and *OCA2* in wild-type (WT) and heterozygote (HET) cells for Denisovan-derived *Alu* insertion in *OCA2* (scale bar: 400 µm)**

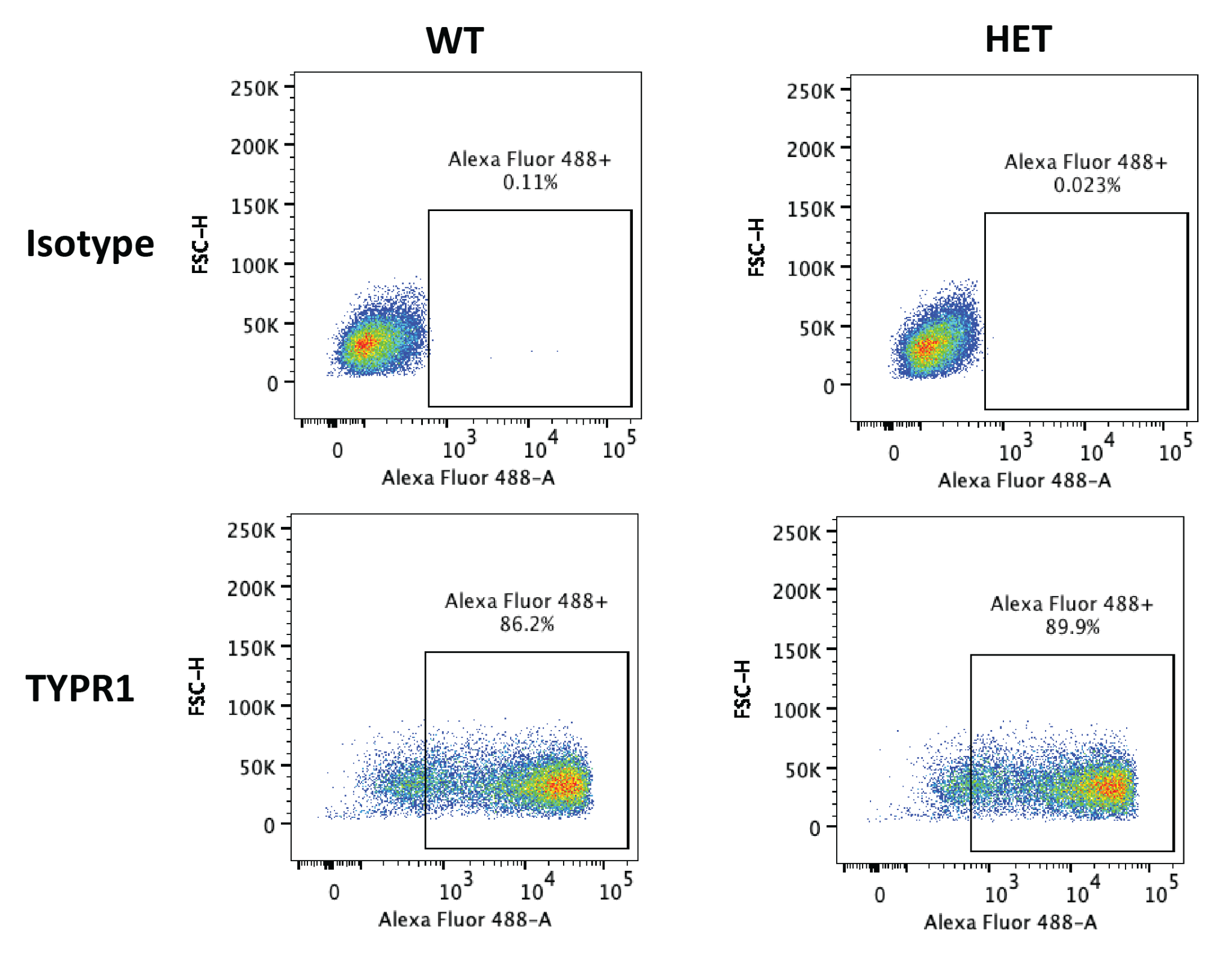

**Supplementary Figure 38. Flow cytometry analysis of *TYRP1* expression in differentiated KOLF2.1J cells at day 30.** The isotype control represents antibodies lacking specificity to the target antigen, *TYRP1*, and is used to assess background fluorescence. Over 85% of the differentiated cells exhibit positive staining for *TYRP1*, a marker of mature melanocytes, indicating successful lineage specification.
