## Supplementary figures and images for "A Denisovan-derived *Alu* insertion in *OCA2* contributes to pigmentation diversity in present-day Melanesians"

### Figure S7

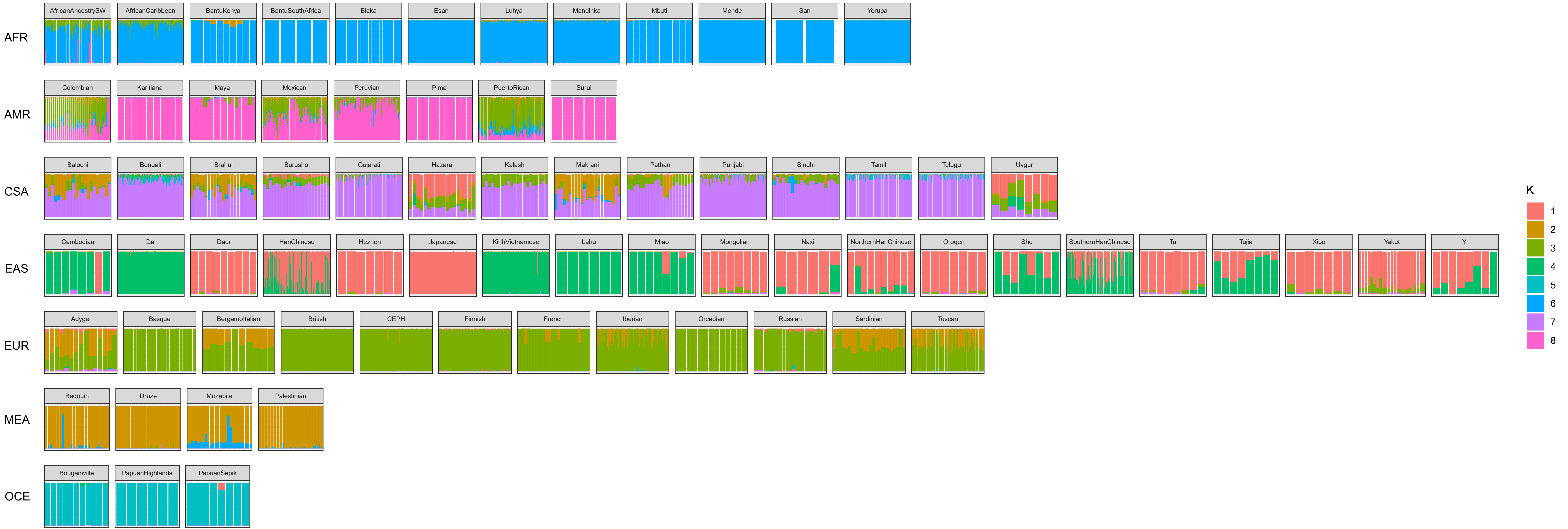

### Figure S33

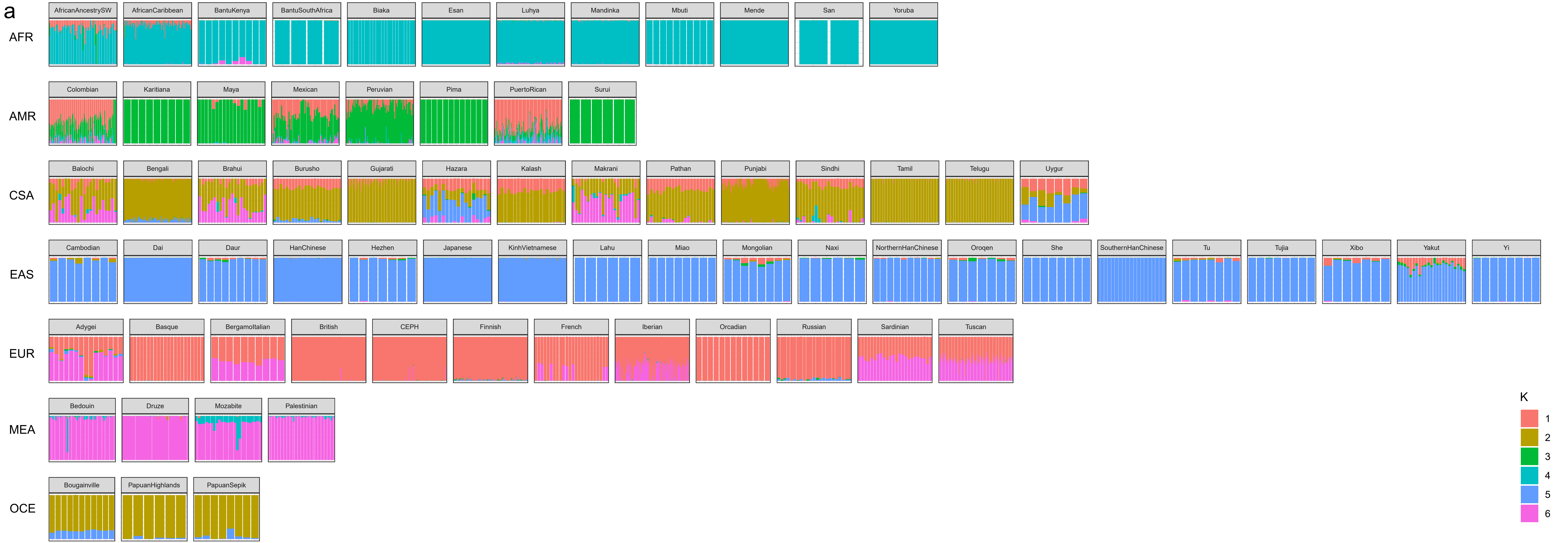

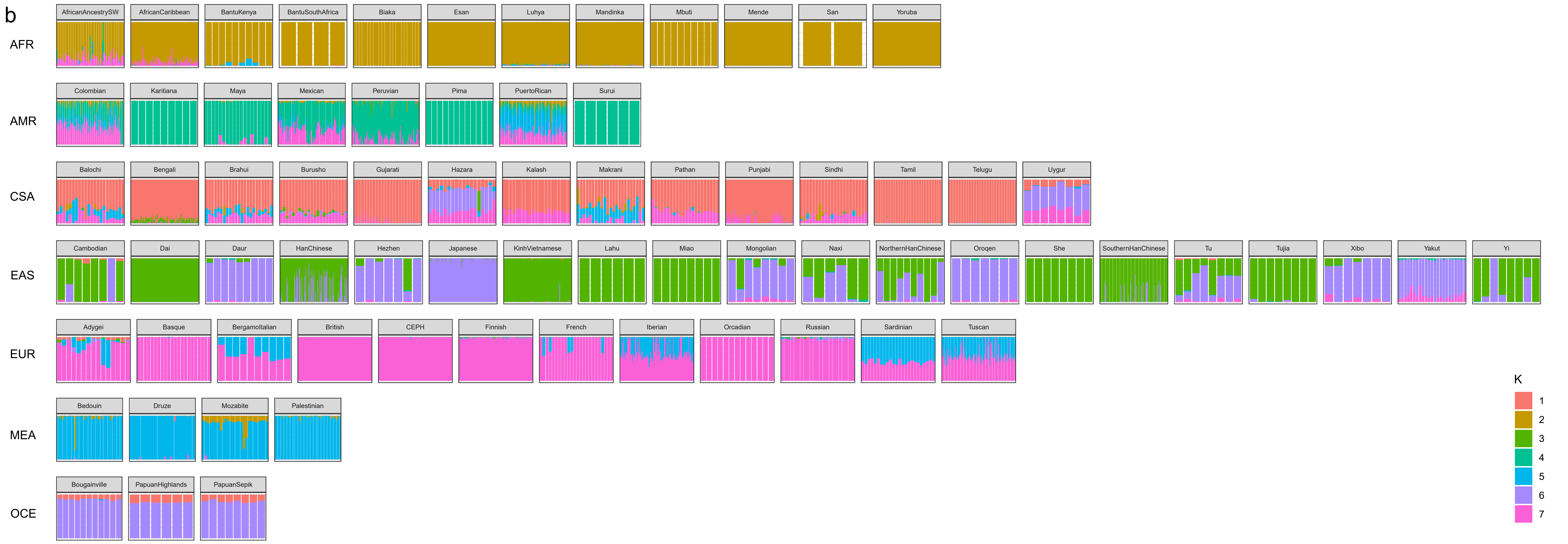

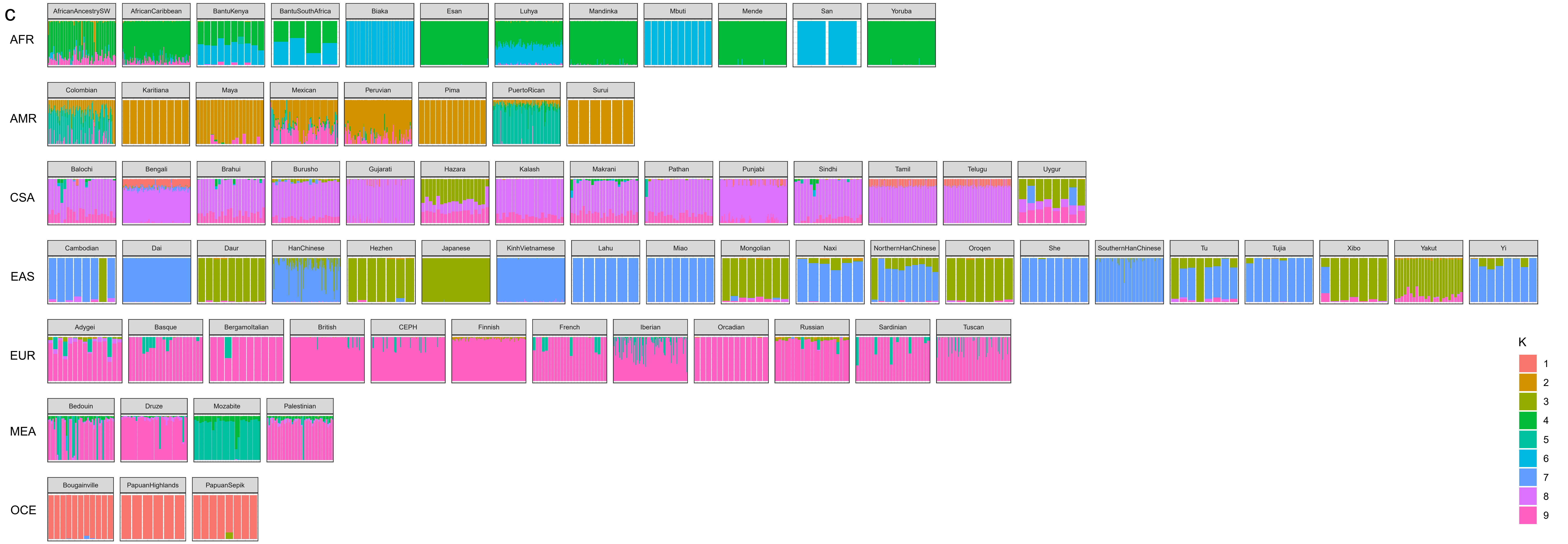

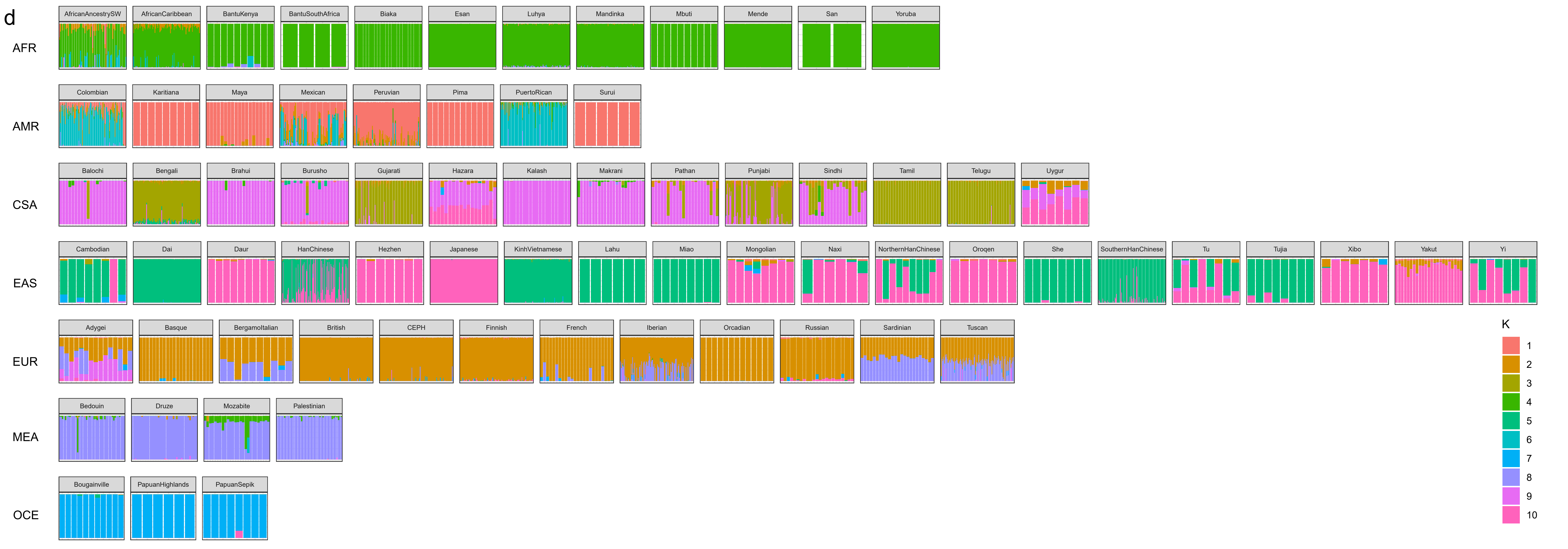
